# A Human Brain Map of Mitochondrial Respiratory Capacity and Diversity

**DOI:** 10.1101/2024.03.05.583623

**Authors:** Eugene V. Mosharov, Ayelet M Rosenberg, Anna S Monzel, Corey A. Osto, Linsey Stiles, Gorazd B. Rosoklija, Andrew J. Dwork, Snehal Bindra, Ya Zhang, Masashi Fujita, Madeline B Mariani, Mihran Bakalian, David Sulzer, Philip L. De Jager, Vilas Menon, Orian S Shirihai, J. John Mann, Mark Underwood, Maura Boldrini, Michel Thiebaut de Schotten, Martin Picard

## Abstract

Mitochondrial oxidative phosphorylation (OxPhos) powers brain activity^1,2^, and mitochondrial defects are linked to neurodegenerative and neuropsychiatric disorders^3,4^, underscoring the need to define the brain’s molecular energetic landscape^5–10^. To bridge the cognitive neuroscience and cell biology scale gap, we developed a physical voxelization approach to partition a frozen human coronal hemisphere section into 703 voxels comparable to neuroimaging resolution (3×3×3 mm). In each cortical and subcortical brain voxel, we profiled mitochondrial phenotypes including OxPhos enzyme activities, mitochondrial DNA and volume density, and mitochondria-specific respiratory capacity. We show that the human brain contains a diversity of mitochondrial phenotypes driven by both topology and cell types. Compared to white matter, grey matter contains >50% more mitochondria. We show that the more abundant grey matter mitochondria also are biochemically optimized for energy transformation, particularly among recently evolved cortical brain regions. Scaling these data to the whole brain, we created a backward linear regression model integrating several neuroimaging modalities^11^, thereby generating a brain-wide map of mitochondrial distribution and specialization that predicts mitochondrial characteristics in an independent brain region of the same donor brain. This new approach and the resulting MitoBrainMap of mitochondrial phenotypes provide a foundation for exploring the molecular energetic landscape that enables normal brain functions, relating it to neuroimaging data, and defining the subcellular basis for regionalized brain processes relevant to neuropsychiatric and neurodegenerative disorders.

## Main

Functional neuroimaging techniques capture dynamic electrical, metabolic and hemodynamic brain energy states^12–15^ but offer only indirect measures of the underlying subcellular bioenergetic processes. All basal and activity-dependent brain processes depend on cellular energy transformation, or bioenergetics, involving ATP synthesis by oxidative phosphorylation (OxPhos) within trillions of respiring mitochondria^16^. There are 100-1,000s of these organelles per neuron or glial cell^17^. Mitochondria molecularly specialize to meet specific cellular demands, direct subcellular activities^5,6,17–19^, and provide the necessary energy to support brain activity. Beyond energetics, mitochondria also participate in other critical functions including cell-cell signaling^20^ and regulation of neuronal excitability^21^, neurotransmitter release^22^ and modulation of inflammatory processes^23,24^. As a result, mitochondria must play a vital role in several supporting functions distributed across the brain’s large-scale networks. Recent mechanistic studies underscore mitochondria’s influence on cognition^25,26^ and behaviors^27,28^. However, mitochondria are typically examined at the sub-micron scale in cell biology, representing a major methodological and conceptual scale gap with systems neuroscience, which operates at the scale of millimeters when the whole brain is imaged by magnetic resonance imaging (MRI) at conventional field strength. Therefore, a major knowledge obstacle keeping us from resolving the energetic forces that power and direct complex human brain dynamics is the spatial distribution of energy-transforming mitochondria across brain structures.

Preliminary work has begun bridging neuroimaging with microscopic anatomy, providing breakthroughs for cellular imaging interpretation of human neuroimaging findings^29–31^. Such advances aspire to inform our understanding of brain development, cognition, mood, and the mechanisms underlying various neuropathologies. To connect cognitive neuroscience and cell biology, here we develop a method to physically voxelize frozen human brain tissue, systematically profile mitochondrial molecular and biochemical diversity across a coronal brain section, and provide an algorithm for predicting mitochondria distribution, density and OxPhos capacity based on MRI data.

### Brain voxelization

To systematically map mitochondrial distribution, diversity, and molecular specialization across the human brain, the first challenge is to physically partition frozen brain tissue at a spatial resolution comparable to MRI. This would enable mapping molecular/biochemical profiles into standard neuroimaging stereotaxic space (Fig. 1a-g).

**Fig. 1.**
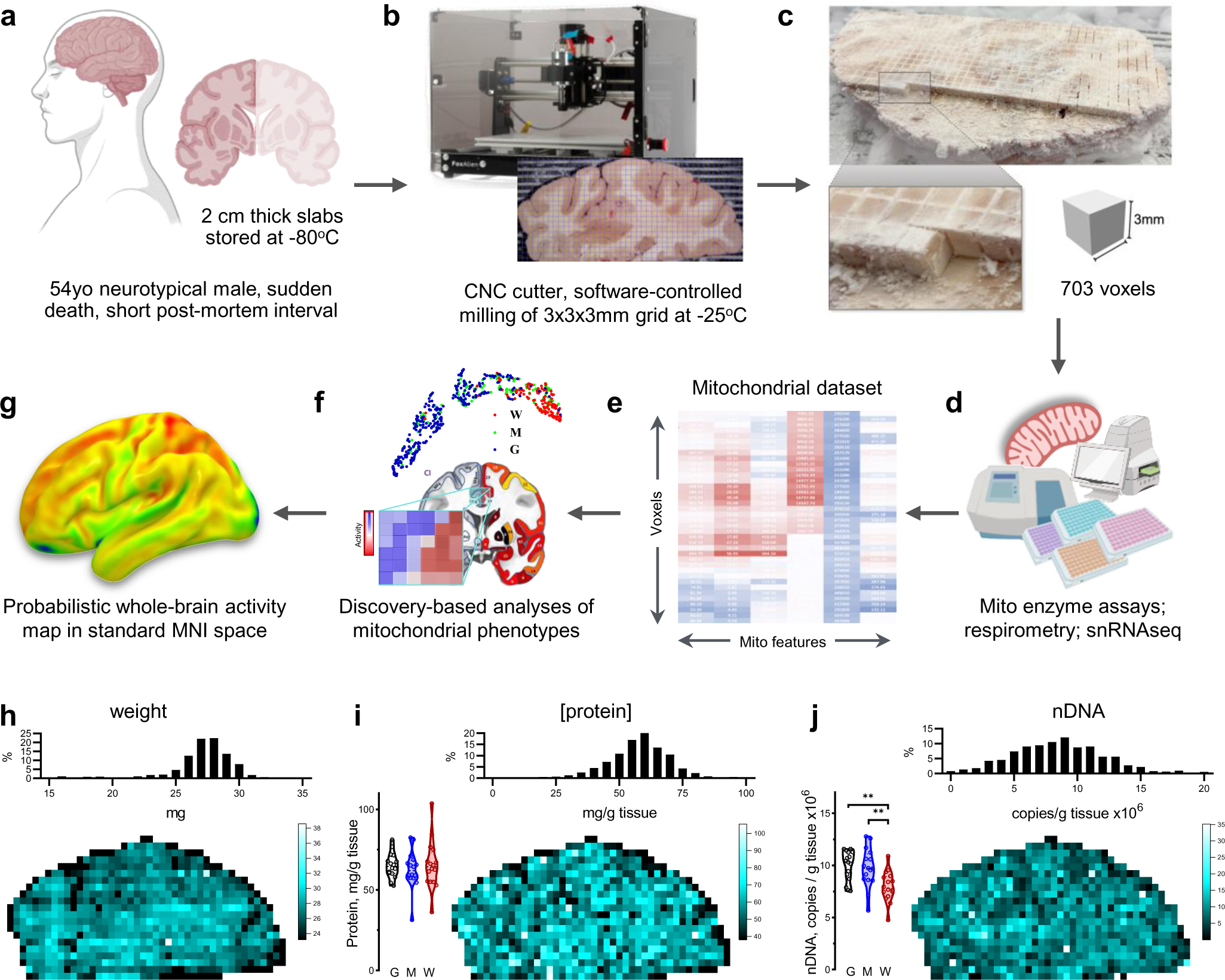
Overall strategy of brain voxelization and mapping. **a,** Postmortem brain tissue was sectioned into ∼2 cm thick coronal slabs, flash frozen in refrigerant HFC-134a and stored in −80°C freezer for 10 years (see Methods). **b,** A slab located at stereotactic Montreal Neurological Institute (MNI) coordinates 15.51 mm posterior to the center of the anterior commissure^63^, was mounted on a computer numerical control (CNC) cutter operated in a −25C freezer room. **c,** The top surface was cleaned and leveled, and a square 3×3 mm grid was milled with a 0.4 mm drill bit to the depth of 3 mm. Brain voxels were manually collected, the surface cleaned once more, and several 50 µm cryosections were collected for histological evaluation. **d,** Each of the >700 samples was weighed, homogenized and ran through an array of biochemical tests, generating matrixes of mitochondrial features linked to voxel coordinates on the brain slice. **e,** Each voxel was characterized based on relative activities and abundance of mitochondrial complexes. **f,** Dimensionality reduction and clustering analysis was performed to identify different mitochondria phenotypes. Maps of mitochondrial features were registered onto a standard MNI space and correlated with average MRI readouts for the same brain regions. **g,** Finally, MRI data was used to predict mitochondrial features and to extend mitochondrial maps to the whole brain. **h-j**. Basic properties of collected voxels, including their (h) weight, (i) protein content and (j) nuclear DNA content. Histograms of each parameter values are shown on top and mapping of values on the brain slice at the bottom. Bar graphs on the left show repeated measurements of the corresponding values in control gray (G), white (W) and mixed (M) matter samples from the occipital lobe of the same brain that were used as normalization controls in the assay plates (see Extended Data Fig. 2).

To address this challenge, we selected a postmortem neurotypical brain (male, 54 years old) with short postmortem interval (PMI = 8 hours, storage at −80°C for 10 years), with negative toxicology for psychoactive medication, drugs and alcohol, negative neuropathology, and negative history of neuropsychiatric disorder (see *Methods*). A coronal slab of the whole right hemisphere, including cortical and subcortical anatomical structures, was voxelized while maintained frozen at −25°C to preserve enzyme activities and molecular integrity during processing. To physically partition the frozen brain slab at 3 mm isotropic resolution comparable to MRI, we programmed a computer numerical control (CNC) cutter to engrave a 3×3 mm square grid at 3 mm depth, followed by manual collection and indexing of a total of 703 samples (Extended Data Fig. 1&2, Supplementary Video 1).

### Mitochondrial phenotyping of brain voxels

To perform mitochondrial phenotyping at unprecedented scale on hundreds of physical human brain voxels, each sample was first randomly assigned to a well across 96-well plates (later deconvolved using a custom algorithm, Extended Data Fig. 3 and *Methods*), weighed, homogenized and subjected to quality control and basic characterization of total protein concentration and nuclear DNA content. These parameters demonstrated relatively uniform quality across the brain section (Fig. 1h-j). The average voxel weight was 26.2±4.5 mg (mean±SD; predicted weight = 27 mg from 3×3×3 mm voxel dimensions at aqueous volumetric density). Such small tissue samples require high-sensitivity assays for both mitochondrial density and enzymatic activities of the respiratory chain (also known as the electron transport chain, ETC)^5^. Mitochondrial density was assessed using a combination of two mitochondrial markers: citrate synthase (CS) activity and mitochondrial DNA (mtDNA) density. OxPhos enzymes reflecting energy transformation capacity were indexed using three markers: complex I (NADH-ubiquinone oxidoreductase), complex II (succinate dehydrogenase, SDH) and complex IV (cytochrome *c* oxidase, COX) which transport electrons and generate the life-giving transmembrane potential that ultimately powers ATP synthesis by the OxPhos system^20^.

For robustness, OxPhos enzymatic activities were quantified using two independent assays in different laboratories: a miniaturized colorimetric assay optimized for brain tissue^5^ and frozen-tissue respirometry^32,33^ (Extended Data Fig. 3). To increase technical accuracy across the dataset, standard reference samples from the occipital lobe of the same brain (grey matter (GM), white matter (WM), and voxels of mixed (M) grey/white composition) spanning the spectrum of possible activities were assayed in duplicate within each batch (eight 96-well assay plates) and used to correct for potential batch effects. The result is a uniform MRI-resolution dataset of mitochondrial activity profiles across the entire coronal section of the human brain (Extended Data Fig. 4). We excluded voxels with either too little tissue (samples located at the brain section’s edges) or enzymatic activities below the detection limit (mainly from the WM), resulting in ∼10% of the voxels missing at least one measure. All mitochondrial features (CI, CII, CIV, CS and mtDNA) were determined in 633 voxels, requiring 27,820 individual samples analyzed by colorimetric, respirometry, biochemical and qPCR assays, including replicates, controls and standards (see Methods).

### Maps of brain mitochondrial density and OxPhos capacity

Mitochondria specialize molecularly and functionally, exhibiting a wide range of energy transformation capacities^18^. To assess mitochondrial “quality” and functional specialization, we previously developed a simple linear formula where OxPhos activities are divided by mitochondrial mass, yielding an index of OxPhos capacity on a per mitochondrion basis, known as the mitochondrial health index (MHI^34^). Here, with a substantially larger brain mitochondrial biochemistry dataset at hand, we discovered that the distributions of individual mitochondrial metrics were left-skewed (Extended Data Fig. 5).

When spheroid organelles have normally distributed radii, metrics related to the spheroid’s surface area follow a square-root normal distribution, whereas those related to spheroid’s volume are cube-root normalized. A well-established example of such dependence is observed with dense core secretory vesicles in which a similarly left-skewed distribution of neurotransmitter content is normalized by a cube-root transformation^35^. Although mitochondrial morphology is more complex than a sphere, volumetric transformation of mitochondrial content (CS^1/3^ and mtDNA^1/3^) and OxPhos capacity (CI^1/2^, CII^1/2^, CIV^1/2^) parameters adequately normalized the mitochondrial features (i.e., decreased CV, skewness, and K-S distance). This transformation also allowed us to deconvolve the normal distributions of mitochondrial features for both GM and WM voxels, yielding a final parametric dataset of individual mitochondrial features (Extended Data Fig. 5).

As expected, the transformed mitochondrial content markers were strongly correlated (Pearson’s r^2^=0.46, p<0.0001, Extended Data Fig. 6a), providing a basis to integrate CS^1/3^ and mtDNA^1/3^ as *MitoD*, representing mitochondria tissue density for each brain voxel (Fig. 2a-c,e; Extended Data Fig. 7a). Mitochondrial DNA copy number (mtDNAcn), which reflects the number of mtDNA genomes per cell nucleus, is also presented (Extended Data Fig. 7b) but is influenced by cellularity and therefore comparison between brain regions cannot be straightforward^5^. Activities of OxPhos enzymes measured by colorimetric and respirometry assays were also highly correlated (Extended Data Fig. 6b-d), and subsequently averaged to integrate them as a robust measure of tissue respiratory capacity (TRC, Fig. 2c,d), representing the *direct quantification of mitochondrial OxPhos capacity per mg of brain tissue*.

**Fig. 2.**
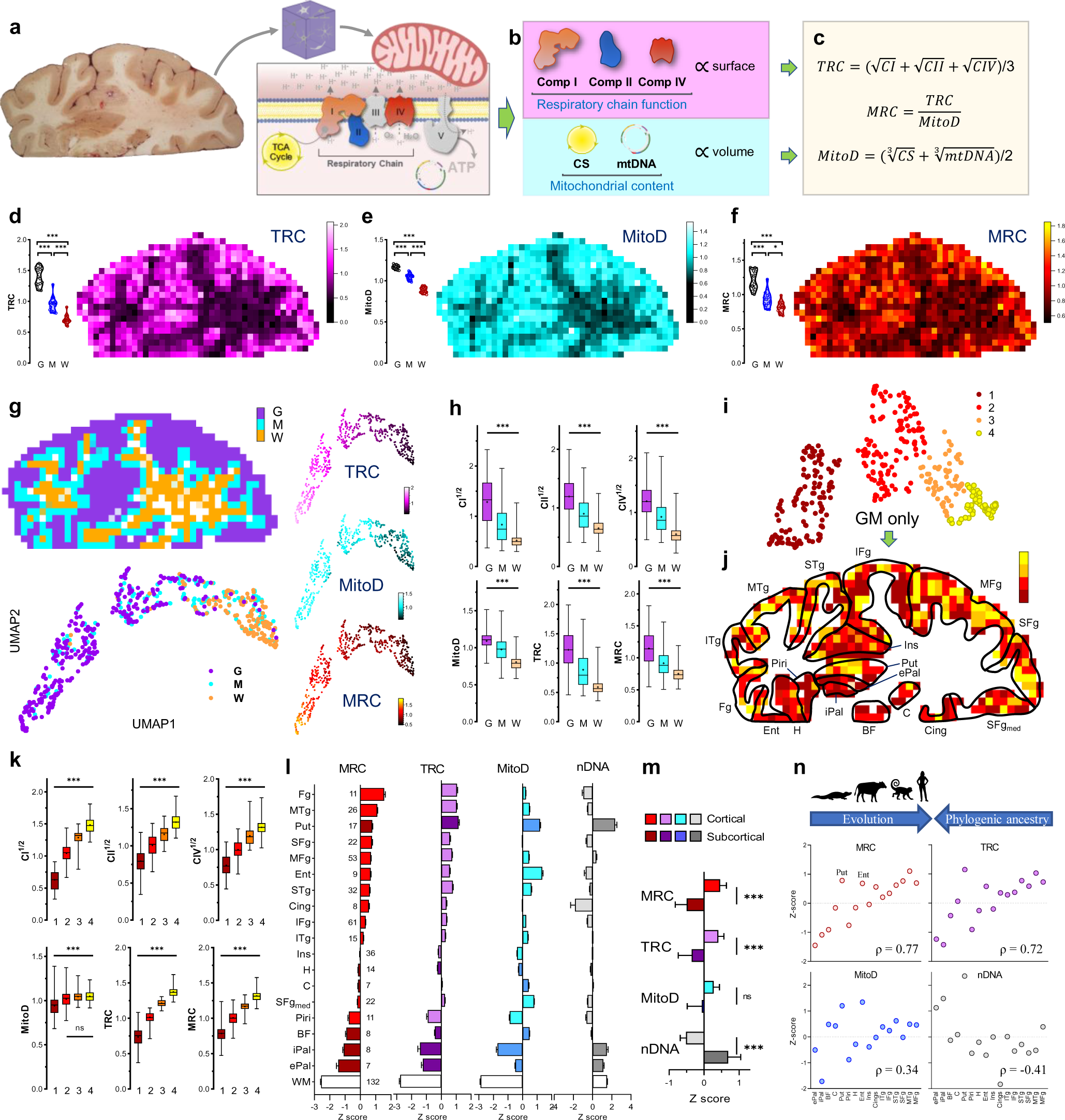
Mitochondrial density and respiratory complexes activity assays. **a,** Human brain slab before voxelization and schematic of the inner mitochondria membrane with respiratory chain (RC) complexes I-V. **b,** If organelle radii are normally distributed, activities of RC complexes that are proportional to the membrane surface should follow square root normal distribution. Citrate synthase (CS) and mtDNA levels are proportional to mitochondria volume and thus should follow a cube root normal distribution. **c,** Formulas for Tissue Respiratory Capacity (TRC), mitochondria density (MitoD) and Mitochondria Respiratory Capacity (MRC). Note that complex activities are normalized by tissue weight during sample preparation. **d-f,** Maps of TRC (d, derived from Extended Data Fig. 4c,d), MitoD (e, average of two panels on Extended Data Fig 4a), and MRC (f). Bar graphs on the left of each panel show distributions of repeated measures of control gray, white and mixed matter samples on different assay plates. **g,** Based on MNI space location (Extended Data Fig. 7), 633 voxels that had all 6 mitochondrial features (CI, CII, CIV, MitoD, TRC and MRC) were divided into gray (G, n=325), white (W, n=132) and mixed (M, n=176) matter clusters. Uniform Manifold Approximation and Projection (UMAP) algorithm was applied for dimension reduction. UMAP plots correlating clusters with physical location of each voxel (left) or z-score values of TRC, MitoD and MRC in each voxel (right). **h,** Bar (25 - 75 percentile) and whiskers (min and max) plots of mito features in voxels mapped to GM, WM and mixed tissue in MNI space. ***-all groups are significantly different from each other by one-way ANOVA with Tukey’s post-hoc test (p<0.001). **i,** UMAP plot and clusters of mitochondrial features of GM voxels. **j,** Mapping of voxel UMAP clusters on specific brain areas (see below for abbreviations). **k,** Comparison of mitochondrial features in four GM clusters identified on i. ***-significantly different from each other by one-way ANOVA with Tukey’s post-hoc test (p<0.001); for MitoD, only cluster 1 is different from all other clusters. **l,** Comparison of mitochondrial features in groups of voxels mapped into specific brain areas in the MNI space. Subcortical areas are shown in darker shades. **m**, Average z-scores of mitochondrial features in cortical and subcortical areas. ***-significantly different by t-test (p<0.01). **n,** Comparison between measured values and the known phylogenetically organization of main brain areas derived from comparative anatomy studies. Spearman’s rho is shown at the bottom-right. ePal, external pallidum; iPal, internal pallidum; BF, basal forebrain; C, caudate; Put, putamen. Areas also present in mammals with some close equivalent in reptiles: Piri, piriform cortex; H, hippocampus. Areas also present in mammals but not in reptiles: Ent, entorhinal cortex; Ins, insula; Cing, cingulate cortex. Neocortex in primates: ITg, inferior temporal gyrus; STg, superior temporal gyrus; SFg, superior frontal gyrus. Neocortex in humans: MTg, middle temporal gyrus; MFg, middle frontal gyrus.

Reference GM voxels exhibited significantly lower values for all mitochondrial metrics than reference WM voxels, with intermediate values in voxels of mixed GM/WM composition (Fig. 2d-e, Extended Data Fig. 4, bar graphs on the left of each panel). This is in agreement with prior neuroimaging-based estimates of brain energy metabolism^5–10^. The data distributions for reference GM and WM voxels exhibited no overlap, highlighting their divergence. As a result, the signal-to-noise ratio to detect differences between GM and WM areas was high, reflected in large effect sizes for all mitochondrial metrics (Hedges’ g=3.5-10.7 for GM *vs.* WM). Compared to the protein and cellularity measures that were homogenously distributed across the section (Fig. 1i-j), the maps of mitochondrial features (Fig. 2d-e, Extended Data Fig. 7) showed clear heterogeneity that overlapped with brain anatomy (Fig. 2a; Extended Data Fig. 1h,i).

To create a volumetric analogue of MHI that reflects tissue-specific specialization of mitochondria for OxPhos and energy transformation, we computed an additional metric that we labeled the *mitochondrial respiratory capacity (MRC)* by expressing each voxel’s TRC relative to MitoD (MRC=TRC/MitoD) (Fig. 2c,f). If each mitochondrion across the brain section had the same OxPhos energy transformation capacity, then MRC should have similar values in all brain areas. Conversely, a difference in MRC would indicate the presence of mitochondria with different OxPhos capacity, or degrees of specialization for OxPhos^34^. Our data establish a variation in MRC both between GM and WM reference samples, and across anatomical structures within the coronal section (Fig. 2f). Thus, beyond the expected variation in mitochondrial density between GM and WM, these results reveal the extent to which maximal respiratory capacity, on a per-mitochondrion basis, varies across the human brain.

### Brain-area specific differences in mitochondrial features

We next registered the coordinates of each voxel to the standard Montreal Neurological Institute (MNI) space (https://www.bic.mni.mcgill.ca/ServicesAtlases/ICBM152NLin2009), the most widely used reference neuroanatomical space for MRI research. Each voxel was manually annotated as GM (n=325), WM (n=132), or mixed (n=176) by an anatomy expert (M.T.d.S.) and labeled with its stereotaxic location (Extended Data Fig. 8). Anatomical identity of the voxels was further confirmed by staining adjacent thin tissue sections for Nissl substance, labeling neurons and glia, and for neuronal nuclear marker NeuN labeling neurons only (Extended Data Fig. 1h,i).

Projecting all annotated voxels with the full set of mitochondrial parameters in a Uniform Manifold Approximation and Projection (UMAP) space produced clusters identified visually that map clearly onto gross voxel composition (GM and WM, Fig. 2g). Conversely, when UMAP clusters were used to predict the origin of the voxels, they mapped GM and WM voxel locations with 64% and 83% accuracy, respectively (chance level = 51% [GM] and 21% [WM], Extended Data Fig. 9a). Furthermore, GM and WM voxels exhibited large differences in all mitochondrial features (g=1.8-2.6; Fig. 2h). Compared to the GM, the WM contained significantly less mitochondrial mass and total OxPhos capacity (MitoD and TRC), but WM voxels exhibited a disproportionally low TRC relative to MitoD. As a result, the MRC reflecting mitochondria specialization for energy transformation on a per-mitochondrion basis, was lowest in the WM voxels (Extended Data Fig. 9b). The difference between GM and WM voxels in all mitochondrial features was replicated in an independent sample from the right occipital lobe (Fig. 2d-f, bar graphs).

Further analysis focused on GM voxels resulted in four UMAP clusters with distinct mitochondrial activity profiles (Fig. 2i-k; Extended Data Fig. 9c). Notably, we observed that despite having similar MitoD (Fig. 2l,m), compared to subcortical brain regions, cortical GM areas exhibited higher TRC (p<0.01, t-test; g=1.1) and MRC (p<0.01; g=1.4). An intriguing exception is the putamen, a subcortical structure with the highest TRC and MitoD. This could be attributed to the high density of terminal synapses/neurites and axonal projections in this area^36^, including large axonal arbors from the tonically active substantia nigra pars compacta dopaminergic neurons^37^, as well as the presence of numerous cell bodies indicated by the highest nDNA content in the putamen. With evolution, the putamen has undergone an evolutionary exaptation that may also account for this difference^38^. Conversely, we find high cell density and low MRC in the internal and external pallidum, as well as among several WM regions.

In comparative anatomy, the presence or absence of structures across species allow for the phylogenic classification of major human brain structures^39–42^. The brainstem, pallidal, and striatal areas are uniformly acknowledged as the oldest, while cortical areas represent the most recent evolutionary development. Our results show that both TRC and MRC correlated strongly with the estimated phylogenic age of brain regions. Mitochondria in regions that emerged later in the evolutionary timeline – moving from reptilian to mammalian and further to primates and humans – exhibited higher MRC (rho=0.72-0.75, p>0.01; Fig. 2n). This enzymatic specialization aligns with the hypothesis that an evolutionary gradient exists with an increased metabolic demand^12^. Our data show that this demand is met by a specialized mitochondrial phenotype with high OxPhos capacity per mitochondrion.

### Brain regions and cell-type specific mitochondrial phenotypes

Specialized mitochondrial phenotypes (i.e., mitotypes) map either to specific brain areas^5^ or to specific cell types^6,43^, which differ in their abundance among different brain areas (WM *vs*. GM; cortical *vs*. subcortical). To profile cell-type specific molecular mitochondrial phenotypes across the human brain, we performed single-nucleus RNA sequencing (snRNAseq) on individual nuclei recovered from voxel homogenates prepared for mitochondrial enzymatic measurements presented above. The lysis-oriented chemistry and aggressive mechanical disruption required for mitochondrial enzyme assays contrast with the gentler approaches typically required to generate high-quality preparations for snRNAseq^44^, a noteworthy limitation of our approach that should be corrected in future studies seeking to establish cell-type specific mitochondrial phenotypes.

We selected individual voxels from four areas: the middle temporal gyrus (MTg), hippocampus (Hipp), putamen (Put), and corpus callosum (white matter, WM), thereby covering a spectrum of brain areas with wide differences in cell type composition. Histochemical staining of neurons and glia (Nissl) or neurons only (NeuN) on 50 µm-thick brain sections confirmed the expected cellular identity of each voxel (Fig. 3a, Extended Data Fig. 10). After applying stringent data cleaning and quality control thresholds, we computationally recovered a total of 32,515 nuclei (MTg: 8,945; Hipp: 7,044; Put: 6,176; WM: 10,350) categorized into 9 broad cell types (Fig. 3b), confirming the expected differences in cell composition across each stereotactically-defined voxel (Fig. 3c).

**Fig. 3.**
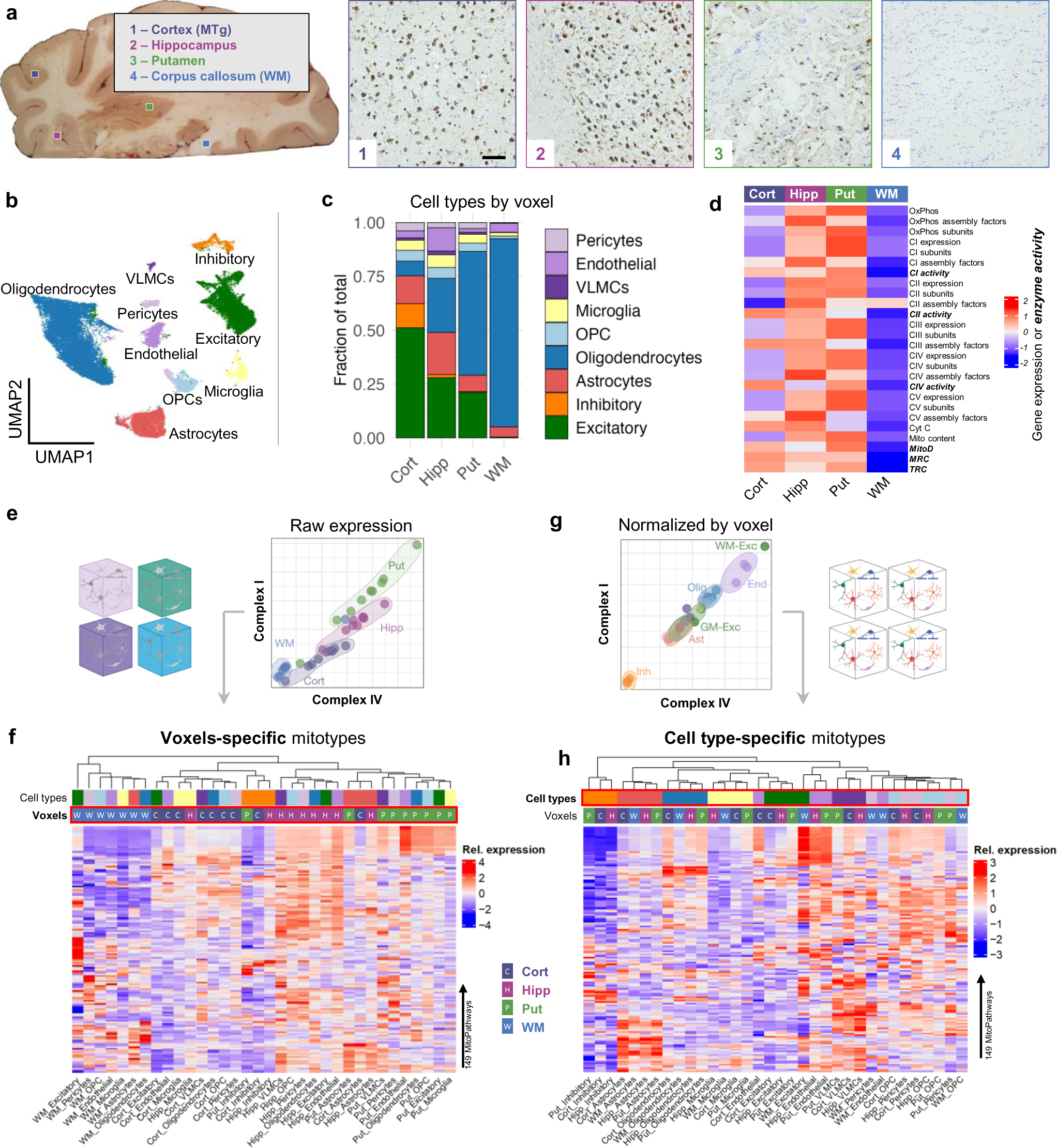
snRNAseq analysis of selected voxels. **a,** Location of voxels selected for snRNAseq analysis and their histological evaluation. Thin brain slices were stained with Nissl (blue, all nuclei) and immunolabeled for NeuN (brown, neuronal cells). Scale bar is 100 µm. **b**, UMAP plot of cell clusters identified by snRNAseq analysis of 4 voxels shows 9 major cell types. **c**, Cell type proportions in each of the 4 voxels. **d**, Heatmap of OxPhos pathway gene expression scores of each voxel (pseudobulk expression per sample), together with the activity measurements (bold). **e**, Correlations between complexes I and IV expression. Each data point represents pseudobulk expression of OxPhos subunits in each voxel/cell type and is color-coded by the voxel type. **f**, Heatmap of 149 mitochondrial pathway expression scores (see Supplementary File 1). Each column represents raw expression scores in a voxel/cell type color coded in the first two rows (same colors as in *a* for voxels and *c* for cell types). Note clustering of the samples by their voxel type. **g**, Correlations between complexes I and IV expression after z-score normalization of expression levels within each voxel type to account for region-specific differences. Each data point is color coded by its cell type. **h**, Heatmap of mitochondrial pathways expression scores in voxels/cell types after expression was normalized within each voxel. Clustering of samples by their cell type is now obvious.

Mitochondrial specialization arises from the regulated expression of ∼1,136 validated mitochondrial genes/proteins from MitoCarta3.0 (Ref ^19^). Using pseudo-bulk transcriptomes (i.e. mimicking bulk RNAseq), we recovered 1,118 nuclear-encoded mitochondrial genes (not mtDNA) and calculated OxPhos scores for each voxel and OxPhos subunit^5^. As in our molecular and biochemical results, the WM voxel exhibited lower mtDNA-encoded genes than GM voxels (Fig 3d; logFC < −1.5, adjusted p<0.001). Overall, relative gene expression of OxPhos complex subunits corresponded well with measured enzymatic activities of CI, II and IV in all brain regions (Fig 3d), providing convergent evidence for the observed enzymatic data. One exception was a lower expression of Complex I and II subunits in the cortex that was not reflected in the activity measures. The putamen had the highest scores for both OxPhos expression and enzyme activity, despite the low abundance of recovered neurons, suggesting that either i) the recovered nuclei do not faithfully represent cell types present in the brain areas, or ii) there are region-specific drivers of enzymatic mitochondrial phenotypes independent of cell type composition, such as local activity or functional connectivity patterns.

To investigate how different cell types may underlie mitochondrial phenotypes by brain region, we compared mRNA abundance of two OxPhos complexes (Complex I and IV) for matching cell types in each of the four voxels. The raw transcriptional profiles of OxPhos complexes clustered by voxel rather than cell type (Fig. 3e, Supplementary File 1), indicating that the major driver of the differences in OxPhos transcriptome for each cell type is the brain region in which it resides. We then extended this analysis to 149 mitochondrial pathways (MitoCarta3.0), composite multi-gene scores that de-emphasize the influence of individual genes to create more stable estimates of functionally relevant mitotype specialization. Again, we found that transcriptional profiles were more similar between different cell types from the same brain region rather than between similar cell types from different regions (Fig. 3f). Thus, most of the variance in brain mitotypes is attributable to regional variations.

Interestingly, removing the voxel-to-voxel variation in the overall expression levels by z-scoring mRNA levels within each voxel rather than across the whole dataset revealed cell-type-specific mitotypes (Fig. 3g). For instance, relative to other cell types, inhibitory neurons had the lowest expression of OxPhos complexes, while endothelial cells showed the highest expression. Similar robust clustering by cell type rather than voxel location was observed when all mitochondrial gene expression profiles were normalized within each sample (Fig. 3h).

Overall, our single-nucleus transcriptome analyses indicated that while mitotype differences between human brain cell types exist and are conserved across the brain regions examined, global drivers of mitochondrial gene expression influencing all cell types across brain regions drive the most significant mitochondrial variation, consistent with a topological understanding of brain functional organization^45–47^.

### Associations of brain mitochondrial phenotypes with neuroimaging modalities

We next asked whether mitochondrial phenotypes may be reflected in standard neuroimaging modalities, including T1 and T2, fMRI-BOLD, and diffusion-weighted imaging from hundreds of individuals (Extended Data Table 1, n = 1,870 multi-modal brain MRIs from healthy adults aged 18-35 years^11^). Such correlations would allow us to use neuroimaging data to predict mitochondrial features, thereby extending the map of mitochondrial OxPhos capacity to the whole brain and to eventually use neuroimaging data to derive personalized MRI-based maps of mitochondrial phenotypes.

As described above, each voxel from our frontoparietal section was registered to its corresponding resolution-matched voxel in the standard MNI brain atlas. Accurate mapping of each voxel was manually verified. Samples with a sum of GM and WM probabilities less than 70% were discarded to avoid partial volume effect contamination. The remaining 539 voxels with all mitochondrial features measured and correctly mapped into the stereotaxic space were randomly divided into two groups to train the model (80%, or 431 of voxels) and to test the model (20%, 108 voxels) (Extended Data Fig. 11a). Each mitochondrial feature (CI, CII, CIV MitoD, TRC and MRC) was regressed onto 22 structural, functional, and diffusion-based neuroimaging metrics using a stepwise linear regression model^48^. Some parameters, such as the orientation dispersion (OD) index, a proxy for neurite complexity^49^, correlated positively with all mitochondrial metrics (Extended Data Table 1). On the other hand, some MRI metrics did not correlate with mitochondrial OxPhos, while others correlated differently with specific features with either a positive or negative linear regression weights (Extended data table 1).

Using linear equations from the regression model, the prediction accuracy of mitochondrial features in out-of-sample voxels (20% of samples) based solely on MRI metrics ranged between r^2^=0.26-0.36 (all p<0.0001, Fig. 4a). In comparison, prediction of the null model with scrambled voxels returned accuracies less than 0.1% (p>0.7, Extended Data Fig. 11c). Collapsing the test samples into their respective brain regions, the observed-to-expected correlations were 2-3 times stronger, emphasizing the robust regional variation in mitochondrial phenotypes (Fig. 4b). The out-of-sample accuracy of the model was also tested by using the MRI-based model to predict the mitochondrial markers in the occipital lobe from the same donor brain (not used in the prediction model). When compared across the six mitochondrial features, the model yielded significant agreement between observed and predicted values (r=0.75, Fig. 4h). The predicted and observed mitochondrial feature maps, projected on the coronal slice examined biochemically, showed remarkable similarities (Extended Data Fig. 11d), lending additional support to the model.

**Fig. 4.**
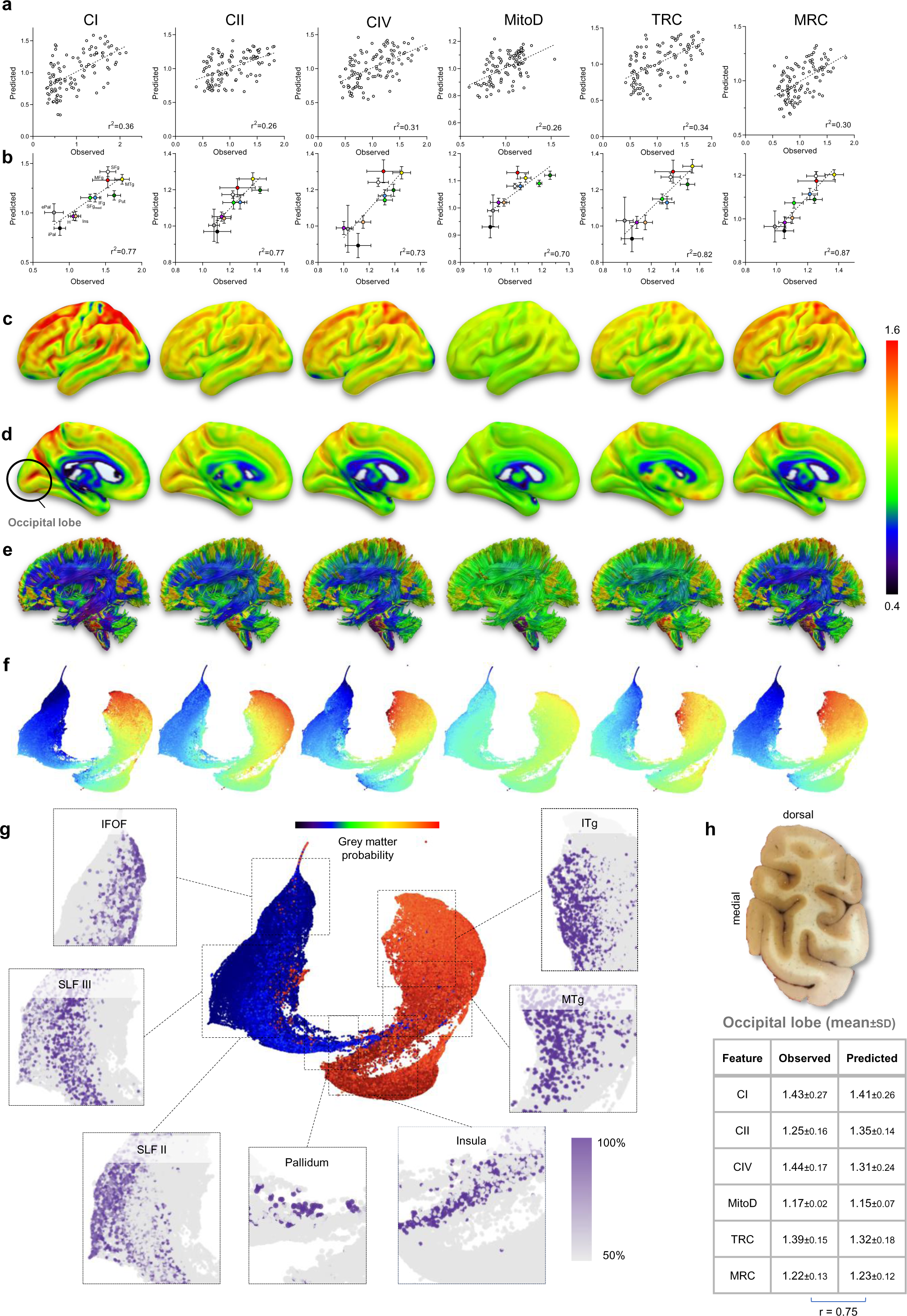
Projected mitochondrial activities of the whole brain. **a,** Scatterplots of 20% out-of-sample prediction of mitochondrial profiles (see Extended Data Fig. 11). **b,** Correlations between observed and predicted mitochondrial features for brain areas rather than individual samples. Both ‘learning’ and ‘testing’ samples were used. n**=**5 voxels (ePal and iPal), 7 (MTg), 9 (MFg), 11 (IFg_med_) 15 (H, Put and SFg), 27 (Ins), 33 (IFg). Slopes of all linear regressions on *a* and *b* are different from zero (p<0.0001; Pearson’s r^2^ is shown on each graph). **c-e,** Prediction of whole brain values for the lateral (c), medial (d) surfaces and the white matter connections (**e,** lateral view). 3D maps are available at https://identifiers.org/neurovault.collection:16418. **f,** UMAP embedding of the whole brain at 1 mm^3^ resolution, colors indicate mitochondrial activity profiles. **g,** UMAP plot of the whole brain with colors indicating the probability of each point being located in the white or gray matter. Insets show probabilities of voxels being in specific white matter (inferior fronto-occipital fasciculus, superior longitudinal fasciculus) or gray matter (pallidum, insula, middle temporal gyrus, inferior temporal gyrus) brain structures (p < 0.0001 Bonferroni corrected for multiple comparisons). **h,** An image of the occipital lobe brain slab before milling (top) and summary table of mitochondrial features in the pooled gray matter OL samples used as loading controls (see Extended data Fig. 3). Predicted values were generated by averaging MRI-based model predicted mitochondrial metrics in 10 randomly selected GM voxels in the MNI space. Observed and predicted datasets are not different by 2-way ANOVA (F_1,144_=0.69, p=0.41); Pearson’s r is shown below the table.

Encouraged by the ability of the model to predict out-of-sample mitochondria characteristics based on MRI features, we next expanded it to calculate mitochondria density and OxPhos activities at the scale of the whole brain. The resulting cortical and WM tracks maps revealed regional specialization for all mitochondrial features (n= 1,827,243 voxels at 1mm^3^ resolution; Fig. 4c-e). Mitochondrial specialization is particularly large for MRC and the OxPhos enzymes, and comparatively more modest for the mitochondrial content (MitoD), suggesting that different brain areas are expected to vary less in their mitochondrial content and substantially more in their mitochondrial specialization for ATP synthesis (i.e. MRC). Accordingly, our neuroimaging-based estimates of MRC across the brain showed a significant correlation with maps estimating brain evolution: evolutionary expansion^50^ and evolutionary variability^51^ (Extended Data Fig. 12), consistent with the region-based analysis shown in Fig. 2n.

Finally, we generated a UMAP representation of the predicted brain-wide mitochondrial phenotypes (anatomical annotation confidence >80%), which further confirmed the striking contrast between GM and WM regions across all mitochondrial features (Fig. 4f,g). Again, the magnitude of gradients was especially large for CI-IV activities and MRC, reflecting the predicted regional specialization of mitochondria. Furthermore, the multivariate UMAP space mapped relatively well with the anatomical/stereotaxis coordinates of cortical and subcortical GM regions and WM tracts, including regions not presented in our initial brain section. Thus, these results establish an approach to use standard MRI parameters to predict biochemical and molecular mitochondrial profiles across the human brain.

## Discussion

We have generated an atlas of mitochondrial content, enzymatic OxPhos activities, and specialization across the human brain, establishing the diversity of mitochondrial phenotypes at a resolution comparable to MRI. Enabled by a novel approach to physically voxelize frozen human brain tissue, our results reveal roles for both regional and cell type-level factors that contribute to molecular mitochondrial phenotype diversity observed throughout the human brain. Notably, the distinct mitochondrial phenotypes between the grey and white matter align with well-known regional variation in metabolic requirements across brain tissue^5–10,12–15^. This also may highlight a predisposition for mitochondrial disorders to impact cortical structures, such as stroke-like episodes that selectively affect and spread across cortical areas with a topology not restricted to arterial vascular territories^52^. These data could also suggest the existence of compensatory mechanisms within the white matter that may mitigate cortical deficits in diseases like stroke^53,54^.

The alignment of mitochondria molecularly specialized for energy transformation (high MRC) with evolutionary patterns sheds light on the underlying subcellular bioenergetic infrastructure evolved to sustain the elevated energy costs unique to humans^55^, particularly in regions associated with human-specific cognitive functions^12^. Recent work highlights how cortical brain regions recruited to perform executive functions express high levels of cell surface receptors for neuromodulators, and depend highly on glucose consumption^12^. This dependence also agrees with the neuropsychological vulnerability of evolutionary advanced cognitive functions among individuals with inherited mitochondrial diseases^56^, and with the cognitive domains affected by the age-related accumulation of defects on the mitochondrial genome^57–59^.

Rapidly accumulating evidence that neuropsychiatric and age-related brain disorders involve mitochondrial alterations has motivated basic and clinical research communities to image mitochondrial biology in the living brain. In parallel with PET-based metabolic imaging, our model predicting molecular mitochondrial phenotypes across the brain using common neuroimaging modalities^11,60^ meets this need and opens the door to exciting questions bridging the scale gap between cognitive neuroscience and mitochondrial biology.

However, some limitations of the present study should be noted. Our prediction model is based on neuroimaging data from a group template and is subject to interindividual anatomical variability. The reliance on a single neurotypical human brain sample, akin to other whole-brain microscopic initiatives^29–31^, underscores the necessity for additional postmortem data sets to capture the full spectrum of mitochondrial expression diversity across individuals of different sex and across a wide age span. The scale of this task promises to be technically challenging. Nevertheless, mapping inter-individual differences in the sub-cellular energetic architecture – in specific cell types, among different brain regions – appears crucial to advance our understanding of the energetic infrastructure that supports and possibly instructs large-scale brain dynamics in different brains^61^.

In sum, our physical voxelization approach and molecular phenotyping study establishes the spatial distribution of mitochondrial diversity within the human brain, revealing robust disparities between grey and white matter and among key cerebral regions. This work not only unveils a gradient that parallels the brain’s phylogenetic development but also provides a scalable approach to deciphering the mitochondrial underpinnings of human brain evolution, development, and disease. Bridging the scale gap from organelle to whole-brain biology and neuroimaging lays the foundation to understand the mitochondrial basis of brain function and dysfunction across a variety of contexts.

## Methods

### Donor brain selection

Donor brains were collected, evaluated, and stored by the Macedonian / NYSPI Brain collection and made available for this study through the Quantitative Brain Biology (Brain QUANT) Institute at New York State Psychiatric Institute (NYSPI). The study was reviewed and approved by the Institutional Review Board at NYSPI. The selected brain was from a 54-year-old neurotypical male who died from a myocardial infarction (PMI = 8 hours, storage time at −80°C for 10 years), had negative toxicology for psychoactive medication, drugs and alcohol, essentially negative neuropathology, and negative history for neuropsychiatric disorders, according to a validated psychological autopsy designed to obtain psychiatric diagnoses by interview of next of kin^62^. According to his relatives, the donor was right-handed, athletic, and did not smoke or drink alcohol. Coronal 2cm-thick slabs of the right hemisphere were flash-frozen in 1,1,1,2-tetrafluoroethane (R-134a, TGS, Skopje, Macedonia) and stored at −80°C. For sectioning, we chose a slab located at stereotactic Montreal Neurological Institute (MNI) coordinates 15.51 mm posterior of the center of the anterior commissure (AC), according to the Human Brain Atlas^63^.

### Tissue Voxelization and Sample collection

To perform brain partitioning while maintaining the tissue below −20°C to ensure that enzymatic activities are least affected, we developed a novel hardware/software platform. The hardware consisted of a CNC (computer numerical control) router (FoxAlien CNC Router 3018-SE V2) operated in a −25°C freezer room and controlled by a laptop computer in the adjacent +4°C room. A software routine in Igor Pro (WaveMetrics, Inc., OR) was developed to 1) easily define the parameters to clean and partition a frozen brain slab, 2) generate a G-codes used to control the CNC router, 3) randomize samples and assign them to assay plates, and 4) derandomize the samples after the assays are complete (Supplementary Files 2&3).

Using an occipital lobe slab from the selected brain, sectioning parameters (drill diameter, feed rate, spindle speed, etc.) were optimized to minimize cutting time while avoiding potential tissue damage due to local heating and maintaining reliable operation during the several hours required for brain cutting. See Extended Data Fig. 1a-c for details on the milling procedure. Once the surface of the brain slab was fully milled, it was placed on dry ice. First, four samples of shavings were collected from the surface above one white (SLF) and three gray (putamen, SFg and MFg) areas. These ‘dust’ samples were processed and quantified together with all other samples. As shown on Extended Data Fig. 2, mitochondrial activities are noticeably affected by local heating and exposure to air.

Next, shavings were gently removed from the surface with a brush and a pre-chilled scalpel and forceps were used to collect individual voxels. The scalpel was inserted into the milled cuts applying light pressure, which released the voxels from the surface of the slab without causing them to pop out. Voxels (703 in total) were then individually picked up with forceps and placed into pre-labeled and pre-weighed 0.5ml PCR tubes on dry ice. Tubes were then re-weighed to determine the weight of the tissue, transferred to 1.5 ml tubes, and stored at −80°C until future processing. To minimize potential batch effects and to blind the experimenter to the origin of the voxel, samples were scrambled using an Igor Pro routine: unique coordinates of each sample on the brain map were substituted with new, randomly generated coordinates on eight 96-well assay plates.

### Brain Histology

After voxel collection, 0.3 mm of the surface was cleaned/leveled again with a 12.7 mm flat-tip drill bit (Extended Data Fig. 1d-f). The metal plate with the slab was then mounted on a freezing microtome and 50 µm-thick cryosections were collected onto glass slides for histological evaluation. For anatomical mapping of neurons and glia, and alignment of cell populations with the collected voxels, we performed tissue staining for neuronal nuclear marker (NeuN), and Nissl as previously described^64^ with some modifications to adapt the immunostaining to frozen tissue.

In brief, for NeuN immunohistochemistry (IHC), whole hemisphere frozen sections mounted on glass slides were dried at room temperature for 1h, then fixed in 4% paraformaldehyde in phosphate buffer (PB; 0.1M; pH 7.4) for 3h at 4°C. After 3 washes in PBS at RT, sections were incubated in 3% H_2_O_2_ and 10% methanol in PBS for 10 minutes, followed by 3 washes in PBS. After 1h pre-incubation in blocking buffer (10% normal horse serum, 0.3% Triton X-100 in PBS), sections were incubated with mouse monoclonal antibodies against NeuN protein (Anti-NeuN, clone A60; 1:20K dilution; Millipore, St. Louis, MO, USA) overnight at 4°C. Sections were then washed 3 times in PBS and incubated with secondary biotinylated horse antibodies against anti-mouse IgG (1:200 dilution; Vector, Newark, CA), followed by avidin/biotinylated peroxidase complex (Vectastain ABC kit elite, Vector) for 2h at RT (1:100 Avidin,1:100 Biotin in PBS) and then 5 min in 0.05% diaminobenzidine (DAB) in 0.1M NaAc.

For Nissl stain with Cresyl violet, after NeuN immunostaining sections were dried in a desiccator overnight followed by delipidation in inceasing percentages of ethanol to 100%, followed by 100% xylene. Then, sections were rehydrated starting from xylene and ending in H_2_O, and stained in Cresyl Violet (0.1% Cresyl violet Acetate, 0.25% glacial Acetic Acid in H_2_O). The staining was differentiated in 90% EtOH + 10% glacial acetic acid. Finally, sections were placed in 100% EtOH followed by 100% xylene before coverslipping.

### Sample Preparation for Mitochondrial Assays and snRNAseq

Homogenization buffer (1mM EDTA and 50mM Triethanolamine) was added to each sample at 1:50 (g:ml) ratio. Homogenization was performed with 2 tungsten beads using a Tissue Lyser (Qiagen, cat# 85300) run at 30 cycles/sec. Samples were homogenized for 1 minute, then incubated on ice for 5 min followed by another 1-min cycle of homogenization. The beads were then removed, and samples underwent 3 freeze/thaw cycles, which was determined to result in higher mitochondrial activities. Samples were placed at −80°C for 15 minutes, then transferred to a RT water bath for 4 min. Homogenate from each sample before the freeze/thaw cycles (100-200µl) was collected for snRNAseq analysis. Protein concentration was measured in 15µL of each sample using the Pierce BCA Assay Kit (Thermo 23225) per manufacturer instructions. Absorbance at 562nm was measured on Tecan Spark plate reader. Sample protein concentration was determined based on the linear regression of an 8-point BCA standard curve (0-20µg) on a plate-by-plate basis. Samples were stored at −80°C.

### Colorimetric Assays of Mitochondrial Enzyme Activities

Enzymatic activities were quantified as previously described^5^ with several modifications. Briefly, activities of Citrate Synthase (CS), Complex I, Succinate Dehydrogenase (SDH/ complex II), Cytochrome C Oxidase (COX/ complex IV) were measured spectrophotometrically in 96-well plates using Spectramax M2 (Spectramax Pro 6, Molecular Devices). Linear slopes reflecting changes in absorbance of the reporter dyes were exported to Microsoft Excel and converted into enzymatic activities using the molar extinction coefficient and dilution factor for each assay. The assays were optimized for the amount of homogenate and substrates to produce stable readings across all samples. Three *loading control* samples - grey matter (GM), white matter (WM), and mixed tissue from the occipital lobe slab of the same brain - were run in duplicates on each plate to ensure that controls would be identical between plates run on different days. Thirty voxels of each tissue type were homogenized and pooled to produce technical replicates of the same samples.

Each sample was assayed in three technical replicates (assayed on different plates) plus the same number of negative controls, except CIV that requires only one nonspecific activity measure. Overall, 8 plates * 4 assays * (3 replicas + 2.5 negative controls) = 176 plates were analyzed by colorimetric assays. Samples were thawed in a room temperature water bath for 4 minutes on the morning of the assays. On each day, 90 samples plus 6 (duplicates of the three) loading controls were transferred into a 96-deepwell block. A 96-multichannel pipette (VIAFLO 96 # 6001) was used to load samples from the block into each of the 96-well triplicate plates and triplicate nonspecific activity plates for each assay. 10 µl of homogenate was used to measure activity for each of the enzymes. Plates were stored at 4°C before the assay. Each day, 22 plates were processed: 6 plates for 3 assays and 4 for the CIV assay.

*Citrate synthase (CS)* enzymatic activity was determined by measuring the increase in absorbance of 5,5’-dithio-bis-(2-nitrobenzoic acid) (DTNB) at 412nm at 30°C in 200μL of a reaction buffer containing: 200 mM Tris (pH 7.4), 10 mM acetyl-CoA, 10 mM DTNB, 2 mM oxaloacetic acid, and 10% w/v Triton-x-100. The rate of conversion of DTNB into NTB^2-^ ions indicates the CS enzymatic activity. To measure non-specific activity, oxaloacetate was removed from the assay mix.

*Complex I (CI)* activity was determined by measuring a decrease in the absorbance of 2,6-dichloroindophenol (DCIP). Kinetics of changes in DCIP absorbance was measured at 600nm at 30°C in 200 μl of a reaction buffer containing: 50 mM potassium phosphate (pH 7.4), 50 mg/ml bovine serum albumin (BSA), 20 mM decylubiquinone, 80 mM nicotinamide adenine dinucleotide (NADH), 20 mM DCIP, 400 µM antimycin A, 50 mM potassium cyanide (KCN). Antimycin A and KCN were used to inhibit electron flow through complexes III and IV. The negative control included CI inhibitors rotenone (500 μM) and piericidin A (200 μM).

*Complex II (CII, Succinate Dehydrogenase [SDH])* activity was determined by measuring the decrease in absorbance of DCIP at 600 nm at 30°C in 200 μl of a reaction buffer containing: 50 mM potassium phosphate (pH 7.4), 50mg/mL BSA, 500 μM rotenone, 500 mM succinate-tris, 50 mM KCN, 20 mM decylubiquinone, 20 mM DCIP, 50 mM ATP, 400 µM antimycin A. The negative control included 500 mM sodium-malonate, which inhibits CII.

*Complex IV (CIV, Cytochrome c oxidase [COX])* activity was determined by measuring a decrease in cytochrome C absorbance. The rate of conversion of cytochrome C from a reduced to oxidized state was measured at 550 nm at 30°C, in 200μL of reaction buffer containing: 100 mM potassium phosphate (pH 7.5), 10% w/v n-dodecylmaltoside and 120μM of purified reduced cytochrome C. The negative control omitted tissue homogenate to determine the auto-oxidation of reduced cytochrome C.

For each assay, activity was determined by integrating the OD change over a certain time period and then subtracting the non-specific activity. As some samples had very different OD kinetics, two separate integration times were used for each assay which ensured that samples that had a delayed start or samples that saturated quickly were all accounted for. CS activity was determined by integrating OD412 change over both 150-250 seconds and 250-500 seconds and by subtracting the non-specific activity. CI activity was determined by integrating OD600 change over 50-300 seconds and 300-600 seconds, and by subtracting the rate of NADH oxidation in the presence of rotenone and piericidin A from the total decrease in absorbance. CII activity was determined integrating OD600 change over 100-400 seconds and 400-700 seconds, and by subtracting the absorbance in the presence of malonate from the total decrease in absorbance. CIV activity was determined by integrating OD550 change over 50-150 seconds and 150-300 seconds, and by subtracting the non-specific activity from the total decrease in absorbance.

For each assay, the later integration time resulted in more consistent measures for control activities across all plates. Therefore, the activities of the control duplicate samples on each plate were averaged for each of the three controls. Each individual control value was then divided by its duplicate average to determine a correction factor. Correction factors for the duplicate control samples on each plate were then averaged. The average normalization factor across the three controls was then used to normalize the plates. Both the earlier integration time values and the later integration time values from each sample were then multiplied by the final normalization factor per plate. Non-specific activity plates were integrated over the full range of the two integration times for each assay (150-500 CS, 50-600 CI, 100-700 CII, 50-300 CIV).

Once plates were normalized, each sample had values for both the earlier integration time and the later integration time on all three triplicate plates. The higher value between the two integration times was selected for each plate. Of the three selected values (one from each triplicate plate), the two closest values were selected, and the least similar value was discarded. This resulted in duplicate activity measures for each sample. Mitochondrial enzymatic activities were determined by averaging the duplicates, and the specific activity of each sample was calculated as the total activity minus non-specific activity (negative control). Non-specific activity was also calculated by taking the two closest of three measures, except for CIV which only has one non-specific activity plate. For CIV the average non-specific activity value from each plate was applied to all samples on the plate.

### Respirometry assays

#### Complexes I, II and IV

The basic protocol optimized for previously frozen tissue samples was used as previously described^32,33^. In short, the brain homogenate samples were loaded in duplicate onto XF96 Seahorse plates to measure the maximal oxidative capacity of either ETC Complexes I and IV or Complexes II and IV on a Seahorse XFe96 instrument. Complex I and IV are measured by sequential injection of NADH, Antimycin A, TMPD/Ascorbate, and Azide, while Complex II and IV are measured by injecting Succinate/Rotenone, Antimycin A, TMPD/Ascorbate, and Azide.

Investigators running respirometry assays were blinded to the sample identity until all processing and analysis of the results were complete. Each Seahorse assay plate had 8 control wells: the GM, WM and mixed tissue controls that were identical to those used for colorimetric assays, mouse liver mitochondrion (a control for Seahorse and injection function), and a blank control. Samples were loaded at 15uL volume and normalized to protein concentration after data acquisition.

<u>Respirometry Data Analysis</u> was performed as previously described^32^. Briefly, oxygen consumption rate (OCR) data were extracted from duplicate Seahorse runs using the Agilent Wave Analysis software. Complex I OCR was calculated as the post-NADH injection OCR minus the post-Antimycin A injection OCR. Complex II OCR: post-succinate/rotenone injection OCR minus the post-antimycin A injection OCR. Complex IV OCR: post-TMPD/ascorbate injection OCR minus the post-azide injection OCR.

After values for Complex I-IV OCR were normalization for protein concentration, traces were analyzed for acceptance criteria. Minimal OCR thresholds (<2SD of the average readings from WM samples) were set to 0.5 pmol/min/μg for Complexes I and II, and 1.5 pmol/min/μg for Complex IV. If a sample had Complex I or II OCR readings that passed their acceptance criteria, Complex IV OCR values must exceed 80% of their respective Complex I OCR or Complex II OCR reading. Overall, one Complex I OCR value, one Complex II OCR value, and two Complex IV OCR values must pass all acceptance criteria. If a sample did not meet the criteria, the assays were rerun with 25μL sample instead of 15μL for the initial run. If a sample did not meet the acceptance criteria after the rerun, the data was not included in the final data set. Overall, out of 703 analyzed samples, 70 samples did not pass acceptance criteria for Complex I activity, 35 for Complex II activity, and 19 for Complex IV activity.

### Mitochondrial and nuclear DNA (mtDNA and nDNA)

mtDNA and nDNA densities as well as mtDNA copy number (mtDNAcn) were measured as previously described^5^ with minor modifications. Homogenates were lysed at 1:10 dilution in a lysis buffer (100 mM Tris HCl pH 8.5, 0.5% Tween 20, and 200 g/mL proteinase K) for 16 hours at 55°C, 10 minutes at 95°C, and then briefly at 4°C. Plates were then stored at −80°C and thawed to 4°C before performing qPCR. The reactions were run in triplicates in 384-well plates using a liquid handling station (ep-Motion5073, Eppendorf), with 12 μl of master mix (TaqMan Universal Master mix fast, Life Technologies #4444964) and 8 μl of the sample lysate, for a final volume of 20 μl. Each plate contained 90 samples, duplicates of the three loading control samples, a negative control (lysis buffer without homogenate), and 8 serial dilutions of HfB1 standard (1:4 dilutions), all of which were measured in triplicates. qPCR reaction with Taqman chemistry was used to simultaneously quantify mitochondrial and nuclear amplicons in the same reactions: ND1 (mtDNA) and β-2 microglobulin (B2M, nDNA). The Master Mix included 300 nM of primers and 100 nM probe: ND1 forward primer: GAGCGATGGTGAGAGCTAAGGT, reverse primer: CCCTAAAACCCGCCACATCT, and the probe: HEX-CCATCACCCTCTACATCACCGCCC-3IABkFQ. The B2M forward primer: CCAGCAGAGAATGGAAAGTCAA, reverse primer: TCTCTCTCCATTCTTCAGTAAGTCAACT, and the probe: FAM-ATGTGTCTGGGTTTCATCCATCCGACA-3IABkFQ.

The plates were sealed, briefly centrifuged at 2,000g, and subjected to 40x cycles on a QuantStudio 7 flex instrument (Applied Biosystems Cat# 448570): 50°C for 2min, 95°C for 20s, 95°C for 1min, and 60°C for 20s. To ensure comparable Ct values across plates and assays, thresholds for fluorescence detection for both ND1 and B2M were set to 0.08. Averages, standard deviations, and coefficients of variation (CV) were computed between qPCR triplicates and an exclusion cutoff of Ct>33 was applied. Control Ct values were averaged across all plates for each of the three controls, and each plates’ control values were then divided by the average to determine a normalization factor. The average normalization factor across the three controls was then used to normalize each plate. mtDNAcn was derived from the ΔCt calculated by subtracting the average mtDNA Ct from the average nDNA Ct and mtDNAcn was calculated by 2^(ΔCt)^×2. For measures of mtDNA density, the Ct value was linearized as 2^Ct^/(1/10^−12^) to derive relative mtDNA abundance per unit of tissue.

### MitoTracker Deep Red (MTDR) staining

Measurement of mitochondrial content in previously frozen sample homogenates using MTDR was performed as outlined in Refs: ^32,33^. Briefly, 2.5-80 µg of sample material (based on BCA protein calculations) was loaded to a clear, flat-bottom 96-well plate together with 100µL of MAS buffer containing 1µM MTDR. After 10min incubation at 37°C, MTDR was removed by centrifuging the plate for 5 min at 2000 x g at 4°C, with no brake for deceleration. The supernatant was carefully removed, and samples resuspended in 100µL of MAS buffer without MTDR. Fluorescence was measured using a Tecan Spark plate reader using 625 nm excitation and 670 nm emission. Relative mitochondrial content was determined after controlling for background fluorescence using a blank well. Note that because of high variability in MTDR assay values it correlated poorly with CS and mtDNA (Extended Data Fig. 6a) and was not used for the analysis.

### Overall number of samples

The following biochemical assays were performed to analyze 703 voxels (including replicates and controls): BCA (8 x 96-well plates, 768 samples), MTDR (8 plates, 768 samples), CI, CII and CS colorimetry (48 plates, 13,608 samples), CIV colorimetry (32 plates, 3024 samples), CI, CII and CIV respirometry (44 plates, 7,384 samples), and mtDNA/nDNA (24 plates, 2268 samples). Overall, 164 x 96-well plates (some not fully loaded) and 27,820 individual readouts, including replicates and controls were analyzed. Of the 703 original samples, 633 voxels had all mitochondrial features (CI, CII, CIV, CS and mtDNA) determined, of which 539 voxels mapped accurately into the MNI stereotaxic space.

### Single nucleus RNA sequencing (snRNAseq)

Frozen cell homogenates from four human brain areas (cortex, hippocampus, putamen, and corpus callosum, Fig. 3a) were used for single nucleus RNA sequencing (snRNAseq). Briefly, samples in homogenization buffer (Clontech, Cat: 2313A), were thawed on ice, then centrifuged at 500 x g for 5 minutes at 4°C, washed with 4 ml ice-cold EZ prep buffer (Sigma, Cat: NUC-101) with 0.2% RNAse inhibitor and incubated on ice for 5 minutes. After centrifugation, the nuclei were washed in 4 ml Nuclei Suspension Buffer (NSB, 1x PBS, 0.01% BSA and 0.1% RNAse inhibitor). Isolated nuclei were resuspended in 100-500 µl NSB depending on pellet size and filtered through a 35 μm cell strainer (Corning, Cat: 352235). Nuclei were counted using the Nexcelom Cellometer Vision 10x objective and an Acridine Orange & Propidium Iodide (AO/PI) stain. 20 µl of the AO/PI was pipet mixed with 20 µl of the nuclei suspension and 20 µl was loaded onto a Cellometer cell counting chamber of standard thickness (Nexcelom, Cat: CHT4-SD100-002) and counted with the dilution factor set to 2.

The single-nuclei library preparation was constructed using 10x Chromium Next GEM Single Cell 3’ Reagent Kits v3.1 (Dual Index) (10x Genomics, Pleasanton, CA) according to the manufacturer’s protocol. Briefly, a total of ∼10,000 nuclei were loaded on the 10x Genomics chromium controller single-cell instrument. Reverse transcription reagents, barcoded gel beads, and partitioning oil were mixed with the cells for generating single-cell gel beads in emulsions (GEM). After reverse transcription and barcoding of the cDNA, GEMs were broken, and pooled fractions were recovered and amplified via PCR. The amplified cDNA was then separated by SPRI size selection into cDNA fractions containing mRNA derived cDNA (>400bp), which were further purified by additional rounds of SPRI selection. GEX sequencing libraries were generated from the cDNA fractions, which were analyzed and quantified using TapeStation D5000 screening tapes (Agilent, Santa Clara, CA) and Qubit HS DNA quantification kit (Thermo Fisher Scientific). Libraries were pooled and sequenced together on a NovaSeq 6000 with S4 flow cell (Illumina, San Diego, CA) using paired-end, dual-index sequencing with 28 cycles for read 1, 10 cycles for i7 index, 10 cycles for i5 index, and 90 cycles for read 2. Samples from the four different brain regions were processed for snRNAseq on the same day to avoid batch effects.

The FASTQ files were processed using the 10x Genomics CellRanger package (v.6.0.0) with human reference data “GRCh38-2020-A” and a command line option “--include-introns”. Raw UMI count matrices were further processed using the Cellbender package (v.0.1.0) with command line options of 2,000 expected cells and 20,000 total droplets to remove debris and doublets. Cells with a range of 200-5000 genes, at least 500 counts, and less than 11% mitochondrial DNA genes were kept, and ribosomal genes were removed. The data was log-normalized, scaled, and doublets were removed using DoubletFinder^65^. The four datasets were integrated using Harmony^66^. Broad clusters were identified using the human motor cortex dataset from Azimuth^67^ and unspecific clusters were removed. Each dataset was down-sampled to 5000 nuclei per dataset for downstream mitochondrial analyses.

For mitochondrial analysis of the snRNAseq data, we first filtered for all human MitoCarta3.0-annotated mitochondrial genes (MitoCarta3.0, Ref ^19^). To calculate MitoPathway scores, we used the average expression of all mitochondrial genes of each respective pathway, as annotated in MitoCarta3.0, either for pseudobulk expression of each voxel, or each voxel and cell type. A detailed description of the data analysis can be found in a supplementary markdown file, linking each figure and analysis with the respective code. Detailed markdown of the RNAseq analysis can be found in Supplementary File 1.

### Uniform Manifold Approximation and Projection for dimension reduction

UMAP method^68^ was used to reduce the dimensions of mitochondrial features. UMAP is a non-linear embedding method that distributes data variability along major axes. It preserves the original pairwise distance between the input data structure over the global distance by projecting data into a newly constructed manifold. The UMAP manifold follows the theoretical framework of Riemannian geometry and algebraic topology. Cells with similar mitochondrial values cluster together in the UMAP morphospace, while those with different features are further apart.

### Voxel registration to Stereotaxic Montreal Neurological Institute (MNI) Space

The image of the brain slab before milling (Extended Data Fig. 1b) was converted to Neuroimaging Informatics Technology Initiative (NIfTI) format using a homemade code (Supplementary File 4). Then, a corresponding grid overlay (Extended Data Fig. 8) was delineated manually onto the converted brain slice using FSLeyes as part of the FMRIB Software Library v6.0 (https://fsl.fmrib.ox.ac.uk/) to produce a digital version of the grid saved as an independent NIfTI file. Each voxel of the digital grid was assigned a unique number and labeled as being gray, white or mixed matter by an expert anatomist (MTS). Additionally, each voxel was labeled with an anatomical label derived from the Atlas of Human Brain Connections^69^.

Next, we registered the coronal slice to the MNI space (MNI152 nonlinear 6th generation; http://www.bic.mni.mcgill.ca) using affine and elastic deformation provided in the MIPAV v10.0 software package (http://mipav.cit.nih.gov). After visual inspection for the alignment, the same deformation was applied to the digital grid to obtain a digital grid in the MNI space (Extended Data Fig. 8). The grid was subsequently split and binarized so that each voxel corresponded to one binary NIfTI file. To create a representation of the mitochondrial values in the MNI stereotaxic space, values collected for each sample were attributed to each corresponding NIfTI file that were subsequently summed to produce individual maps of the 6 mitochondrial parameters (CI, CII, CVI, MitoD, TRC and MRC).

To produce comparisons between these values and neuroimaging, we used neuroimaging values derived from averaged images computed from a sample of 1870 university students^11^ (Extended Data Table 1) and the human connectome project^60^. First, mapping of each voxel within the MNI space was manually verified and samples with a sum of GM and WM probabilities less than 70% (95 samples) were discarded to avoid partial volume effect contamination. The remaining 539 voxels with all mitochondrial features measured and correctly mapped into the stereotaxic space were randomly divided into 80% ‘learning’ (431 samples) and 20% ‘testing’ (108 samples) datasets (Extended Data Fig. 11). Predictive model was built by means of 6 stepwise backward linear regressions with neuroimaging values used as independent variables to predict each of the 6 mitochondrial values (i.e. CI, CII, CIV, MitoD, TRC and MRC) in the learning dataset. In the backward hierarchical linear regression, predictors are eliminated one by one, starting from the least significant. The procedure of elimination continues until the comparison between two consecutive models is not statistically significant. To test a null distribution of the values, the same analysis was performed on a randomized version of the same dataset (Extended Data Fig. 11c). Next, the models were applied to the 20% ‘testing’ dataset to verify out-of-sample quality of predictions. Then, the model was extended to all the brain voxel of the MNI space at 1mm^3^ resolution to produce a whole brain map for each mitochondrial value. 3D maps of predicted mitochondrial features for the whole human brain are available at https://identifiers.org/neurovault.collection:16418.

### Evolution and mitochondrial values

To determine whether increased mitochondrial OxPhos follows a phylogenetic gradient, we used two maps to estimate cortical evolution (Extended Data Fig. 12). The GM variability map has been shown to reflect evolution and individual differences in humans and our closest evolutionary relatives^51^. The second map compares macaque-to-human cortical expansion using a transformation map from Caret^70^. This map was created by aligning 23 landmarks in both species and measuring the geodesic distance between homologous regions (http://brainvis.wustl.edu/).

### Statistical analysis

Data was analyzed and graphed using Igor Pro (Wavemetrics, Lake Oswego, OR; RRID:SCR_000325), Python Programming Language (RRID: SCR_008394), R Project for Statistical Computing (RRID: SCR_001905) and Prism 8.0 software (GraphPad Software Inc, San Diego, CA; RRID:SCR_002798). Data sets with normal distributions were analyzed for significance using unpaired Student’s two-tailed t-test or analysis of variance (ANOVA) followed by Tukey’s post hoc test. Data sets with non-normal distributions were analyzed using the Mann-Whitney U test or Kruskal-Wallis test with Dunn’s adjustment for multiple comparisons. The violin plots are displayed as median ± quartiles; line graphs and data in the text are expressed as mean ± standard error of the mean (SEM). The statistical tests are listed in the figure legends.

## Supplementary Files

<u>Supplementary Video 1</u> | Human brain slab voxelization using a computer numerical control (CNC) cutter. Milling was done over ∼5h at −25°C (freezer room) with routing of the drill bit controlled by a computer in the next room (+4°C).

<u>Supplementary File 1</u> | Detailed markdown of the single nucleus RNA sequencing (snRNAseq) analysis, including the code for data pre-processing, visualization, cluster identification and mitochondrial analysis. This file also shows additional figures supporting the analysis.

<u>Supplementary File 2&3</u> | Igor Pro procedures for 1) defining the parameters to clean and partition a frozen brain slab, 2) generating G-codes used to control the CNC router, 3) randomizing samples and assigning them to specific assay plates, 4) derandomizing the samples after the assays are complete, and 5) plotting the assay data on the topological map of the brain.

<u>Supplementary File 4</u> | Python code used for the conversion of the image of the brain slab to Neuroimaging Informatics Technology Initiative (NIfTI) format.

## Data availability statement

All data supporting the findings of this study are available within the paper and its Supplementary Information.

## Author contributions

E.V.M. and M.P. designed the study and supervised data collection, analysis and interpretation of results. G.B.R., A.J.D., M.B.M., M.B., J.M., M.U. and M.B. (Quantitative Brain Biology, Brain QUANT Institute) collected and stored the donor brain, psychiatric autopsy, and imaging of thin brain slices. E.V.M. oversaw the hardware and software for physical partitioning (voxelization) of the frozen human brain. A.M.R and S.B. collected samples and performed biochemical and molecular assays. C.A.O., L.S., and O.S.S. performed respirometry assays. A.S.M, Y.Z., M.F., V.M. and P.L.D.J. performed snRNAseq analysis. M.T.D.S. registered the data to MNI coordinates, developed the regression model with MRI parameters and created the brain maps. E.V.M., M.T.d.S., A.S.M. and A.M.R. prepared the figures. E.V.M., M.T.d.S. and M.P. drafted the manuscript. All authors reviewed the final version of the manuscript.

## Acknowledgements

This project was supported by NIH grant RF1AG076821 and the Baszucki Brain Research Fund to M.P., a pilot award from the Columbia University Department of Psychiatry to M.P., E.V.M, and M.B., and the JPB Foundation and ASAP to E.V.M. and D.S. M.T.d.S is supported by the European Union’s Horizon 2020 research and innovation programme under the European Research Council (ERC) Consolidator grant agreement No. 818521 (DISCONNECTOME), the University of Bordeaux’s IdEx ‘Investments for the Future’ program RRI ‘IMPACT’, and the IHU ‘Precision & Global Vascular Brain Health Institute – VBHI’ funded by the France 2030 initiative. P.L.D.J. acknowledges the RF1AG057473 grant.

---

**Extended Data Fig. 1.**
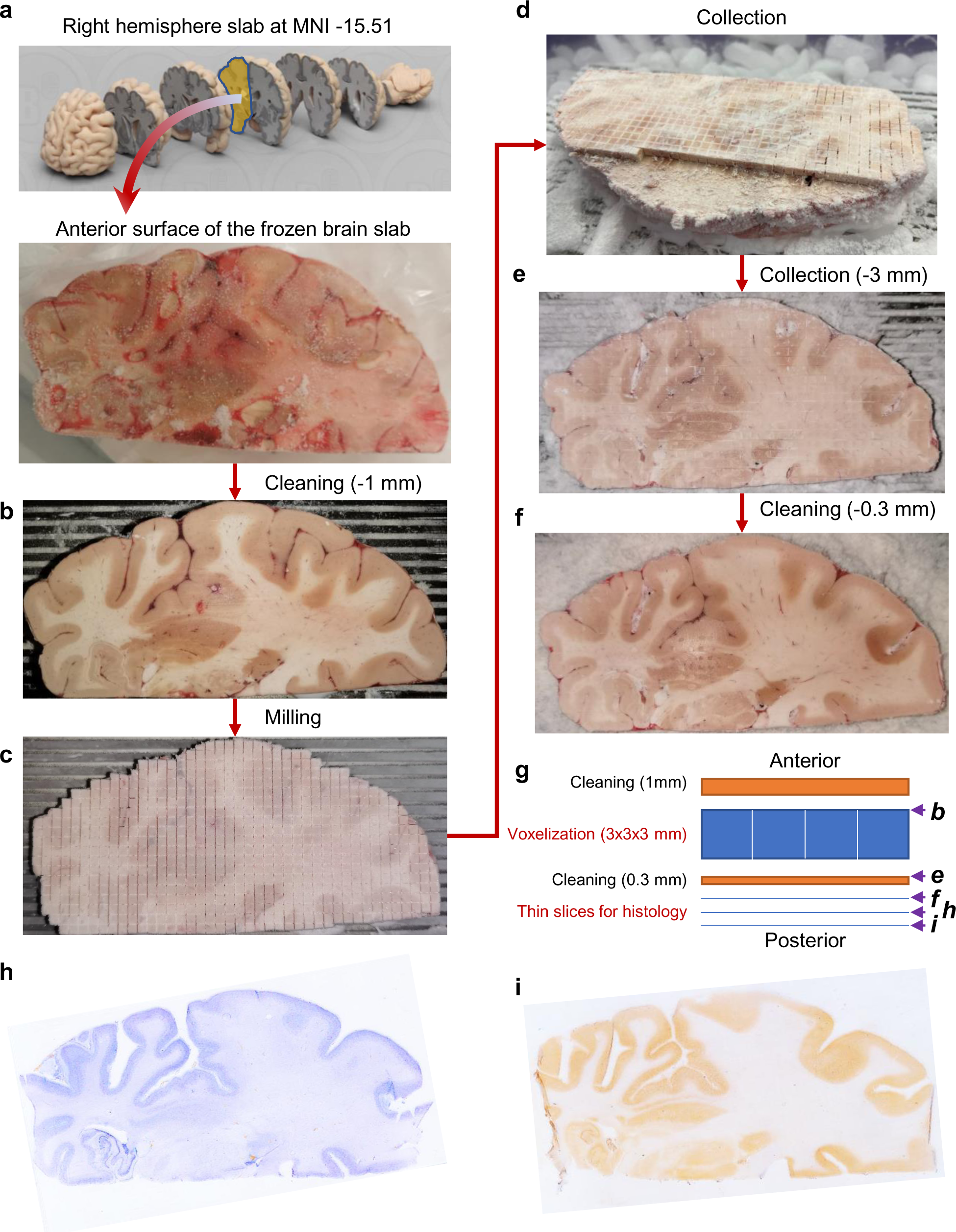
Workflow of tissue collection. **a,** The right hemisphere coronal slab was mounted on a metal plate with an OCT compound with the top (anterior) surface of the slab parallel to the plate. **b,** After affixing the plate to the computer numerical control (CNC) cutting area, the top surface was cleaned with a 12.7 mm flat-tip drill bit rotating at 100 RPM and moving horizontally at 300 mm/sec. After cleaning 1 mm from the top, morphological brain structures were clearly visible and the surface was parallel to the plane of drill bit movement, ensuring that voxels will be of a uniform height. **c,** A 3×3 mm grid was milled with a 0.4 mm drill bit rotating at 10,000 RPM and moving horizontally at 250 mm/sec. During a single pass, 0.2 mm of the depth was milled, requiring 15 passes to reach the desired 3 mm of cut depth. Total milling time for a 130.8 x 61.25 mm hemisphere was ∼5h. **d,** Fully milled slab was placed on dry ice. First, four samples of shavings were collected from the surface above one white and three gray areas (Extended Data Fig. 2). Next, shavings were gently removed from the surface with a brush and a pre-chilled scalpel and forceps were used to collect 703 individual voxels. **e,** Following sample collection, slab surface was cleaned once more with a 12.7 mm drill bit. **f,** The metal plate with the slab was then mounted on a freezing microtome and several 50 µm-thick cryosections were collected for histological evaluation. **g,** Summary of the steps during the collection of brain voxels and thin sections. Letters on the right refer to images shown on the corresponding panels. **h and i,** Thin brain section stained with Nissl to show neurons and glia either alone (h) or in combination with an immunostaining against neuronal nuclear antigen NeuN to highlight neuron-enriched areas (i).

**Extended Data Fig. 2.**
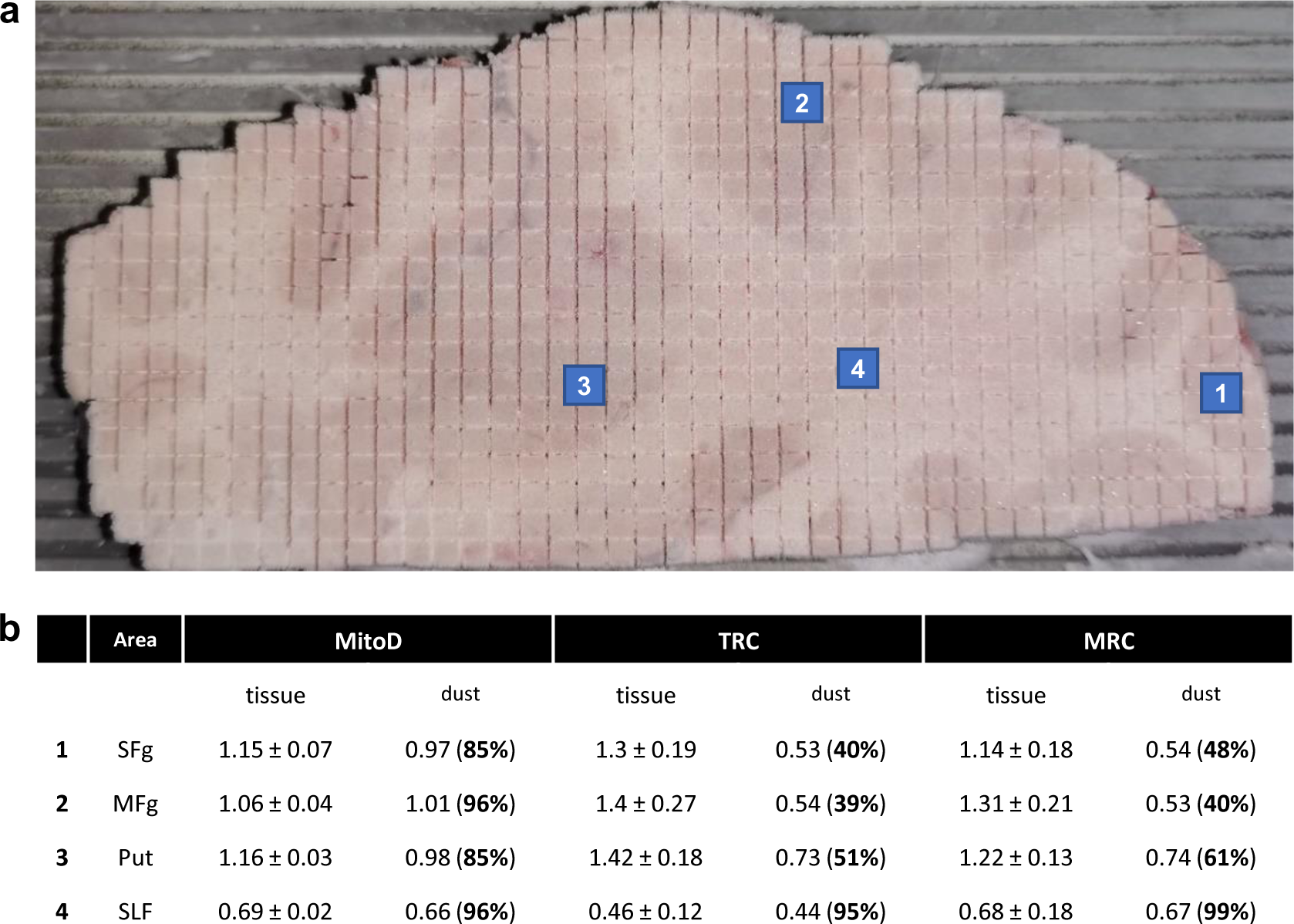
Collection and analysis of tissue shavings. **a,** Positions of collection sites of the four tissue shavings. Samples 1-3 were above the grey matter areas, while 4 was above a white matter area. **b,** Comparison of mitochondrial activities between the four ‘dust’ samples and corresponding tissue samples collected below after tissue voxelization. Values are mean ± SD. While all mitochondrial features (MitoD, TRC and MRC, see Fig 2c for definitions) were lower in all dust samples compared to tissue collected below, MitoD was less different between brain dust and tissue block. Moreover, in sample 4, brain dust and voxels features appear to be very similar, but this likely indicates that colorimetry and respirometry assays are at their detection limit when measuring mitochondria complex activities in the white matter.

**Extended Data Fig. 3.**
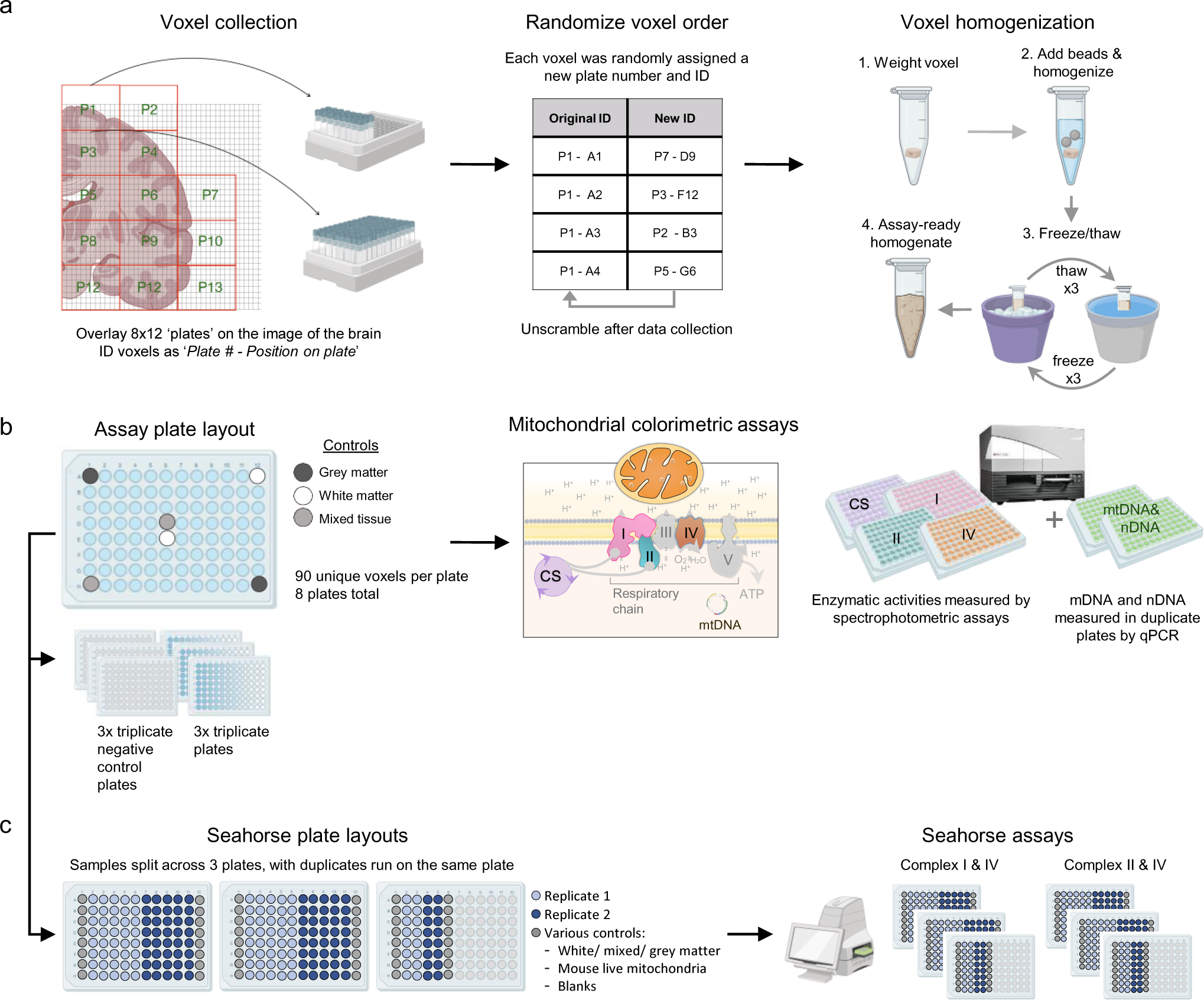
Workflow for mitochondrial colorimetric and respiratory assays. **a,** Brain voxel collection and preparation. **b,** Voxel plating and data collection for colorimetric mitochondrial assay and qPCR. **c,** Frozen tissue respirometry plate layouts derived from the assay plates in *b* to accommodate all samples and ran in duplicate. Samples were loaded into Seahorse plates at a constant volume rather than constant protein content.

**Extended Data Fig. 4.**
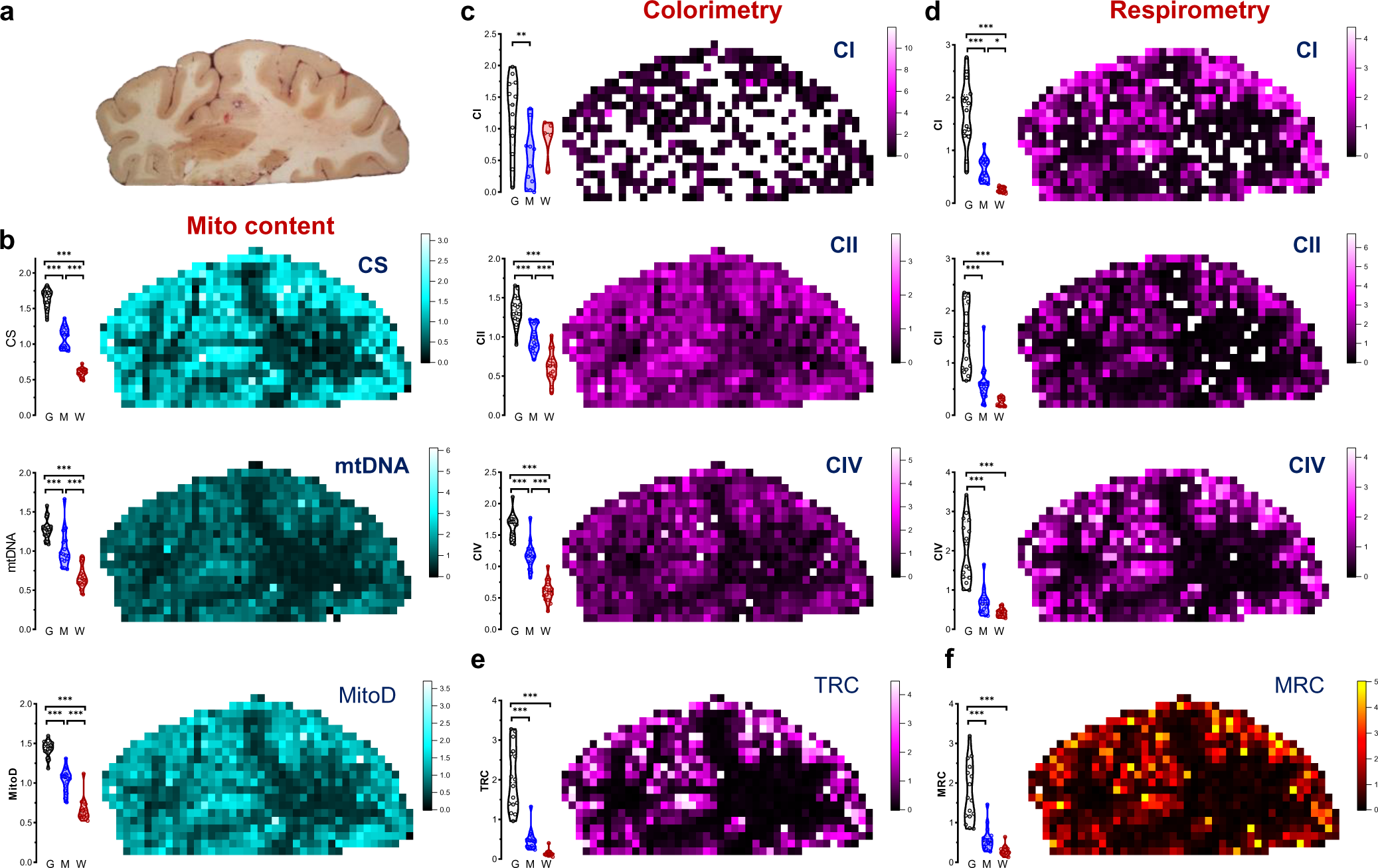
Raw mitochondrial features measured by colorimetric and respirometry assays. **a,** Image of the brain slab before voxelization. **b,** Distributions of non-transformed values (i.e., values are in a linear scale) of citrate synthase (CS) activity, mitochondrial DNA (mtDNA) density and mitochondrial density (MitoD). **c and d,** Mapping of mitochondria complexes I, II and IV (CI, CII and CIV) activities measured by colorimetry (c) and respirometry (d) assays. **e and f,** Tissue Respiratory Capacity (TRC) and Mitochondria Respiratory Capacity (MRC) calculated from raw (non-transformed) values of enzymatic activities. Same maps derived from power transformed values of enzymatic activities are shown on Fig. 2e-f. Activity readouts were normalized to each group’s mean. Bar graphs on the left of the panel show distributions of repeated measures of control gray, white and mixed matter samples on different assay plates (i.e., data points are technical replicates of the same samples).

**Extended Data Fig. 5.**
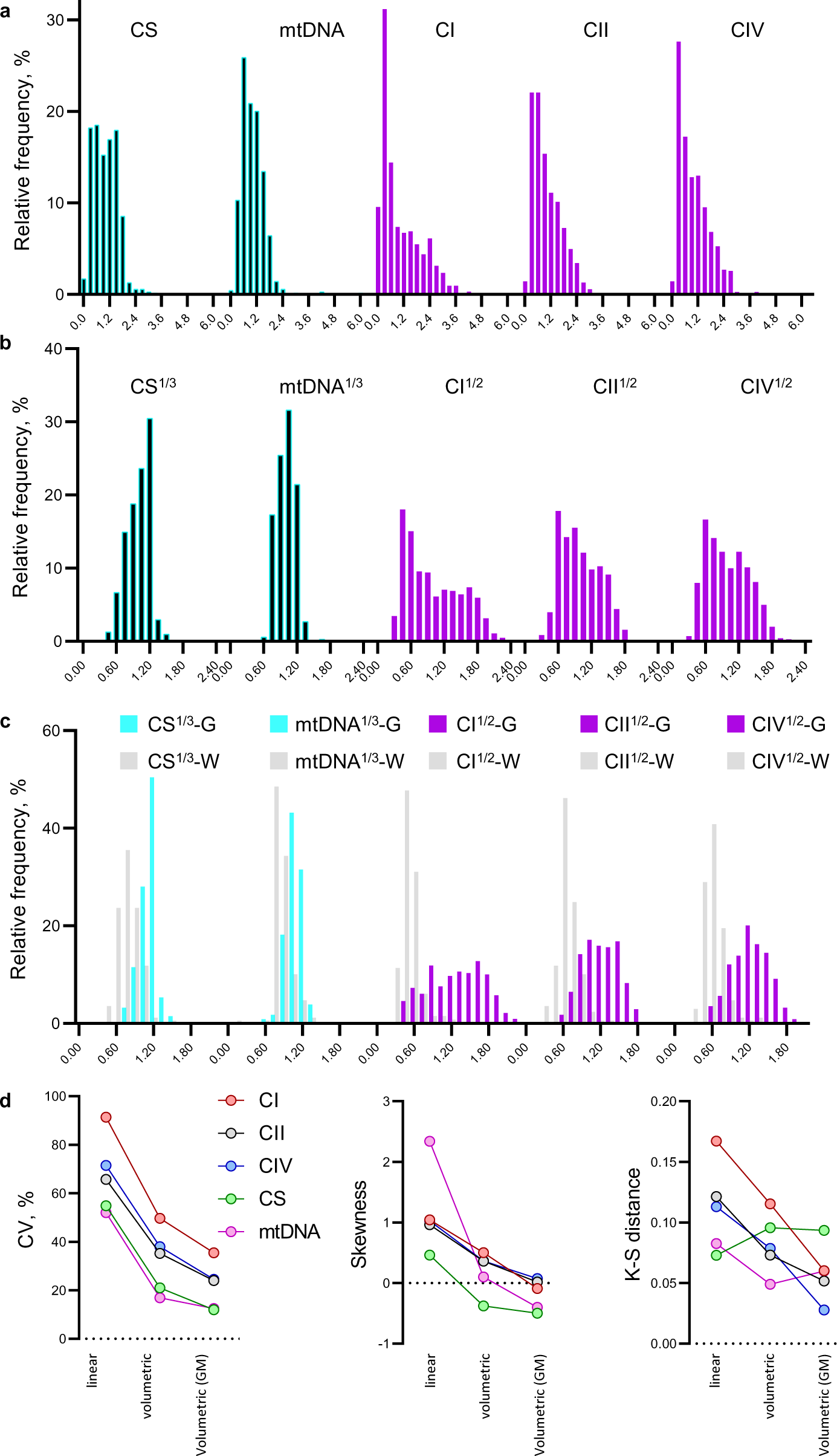
Volumetric transformation of mitochondrial parameters normalizes their distributions. **a,** Distributions of complexes I, II and IV activities, CS activity and mtDNA density. Complex activities are averages from colorimetry and respirometry assays. **b,** Volumetric transformations of the data on *a*. Mitochondria density metrics and OxPhos complexes activities were calculated as averages of cube root and square root values from colorimetric and respirometry assays. **c,** Same data as *b* with voxels assigned to Gray matter (G, n=339) and White matter (W, n=169) clusters based on their anatomical location. Voxels with mixed identity (n=194) are not shown for simplicity.

**Extended Data Fig. 6.**
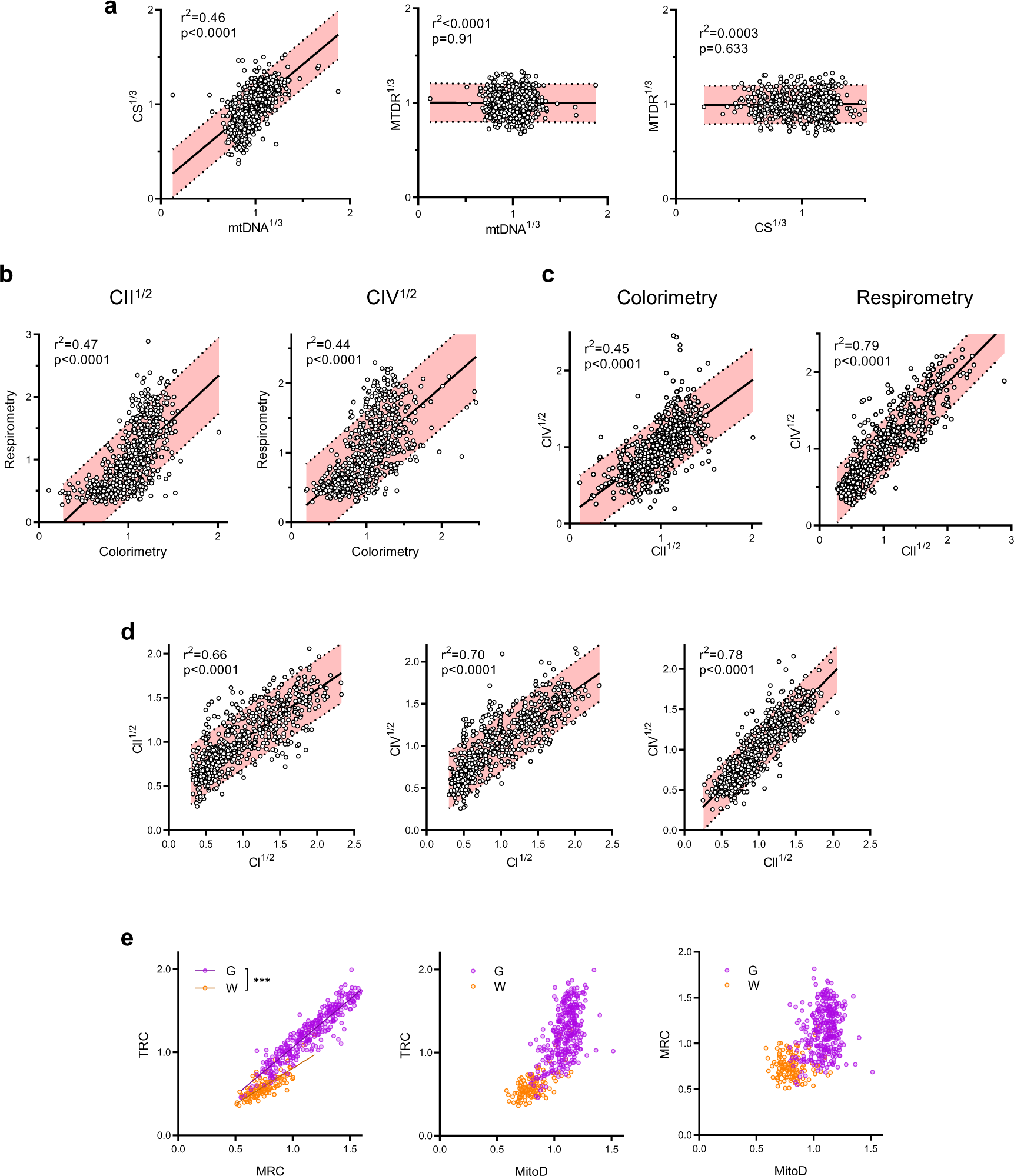
Correlations between mitochondrial parameters assayed using different techniques. **a,** CS activity and mtDNA density, which are related to mitochondria mass, are positively correlated. Because of the high variability in MitoTracker Deep Red (MTDR, a far-red fluorescent probe used to chemically mark and visualize mitochondria in cells) values, they correlated poorly with CS and mtDNA, thus MTDR was not used for the analysis. **b and c,** Relationship between CII and CIV activities measured by colorimetric and respirometry assays. **d,** Correlations between CI, CII and CIV activities (an average between colorimetric and respirometry assays). **a-d,** Pearson’s *r^2^* values show how well datapoints follow the linear regression; *p* values indicate if the slope of the regression line is significantly different from zero; shaded areas represent 90% confidence interval. **e,** Relationship between TRC, MRC and MitoD values (see Fig. 2c for definitions) in gray and white matter voxels. *** - the slopes of linear regressions are different with p<0.001.

**Extended Data Fig. 7.**
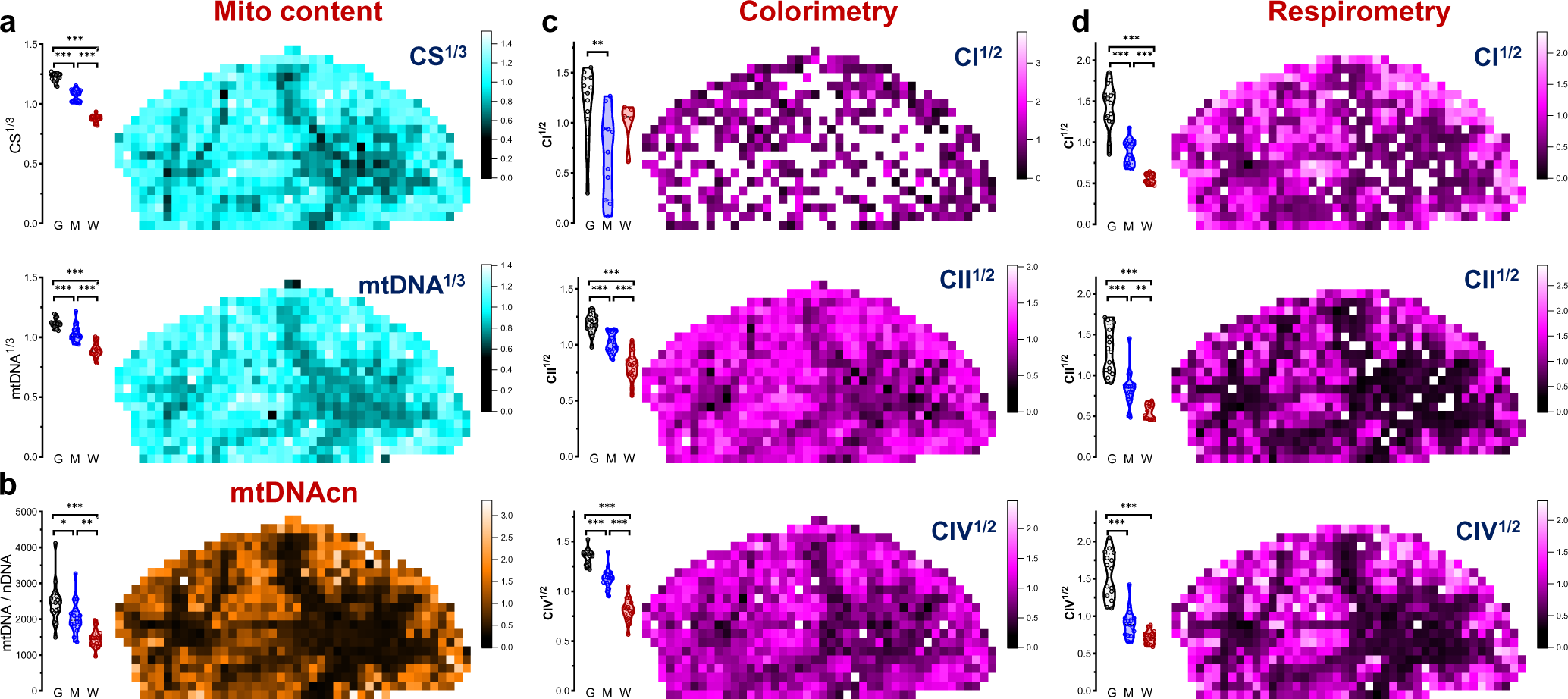
Mitochondrial features measured by colorimetry and respirometry assays. **a,** Distributions of CS activity and mtDNA density. **b,** Measurements of mtDNA copy number (mtDNAcn) derived as a ratio of mitochondrial over nuclear DNA (mtDNA/nDNA). **c and d**. Mapping of CI, CII and CIV activities measured by colorimetry (c) and respirometry (d) assays. Because of the high variability of CI activities measured by colorimetry (c, top), these data were not used for TRC calculation. All values except mtDNAcn are normalized to each group’s mean. Bar graphs on the left of each panel show distributions of repeated measures of control Gray, White and Mixed matter samples on different assay plates (i.e., data points are technical replicates).

**Extended Data Fig. 8.**
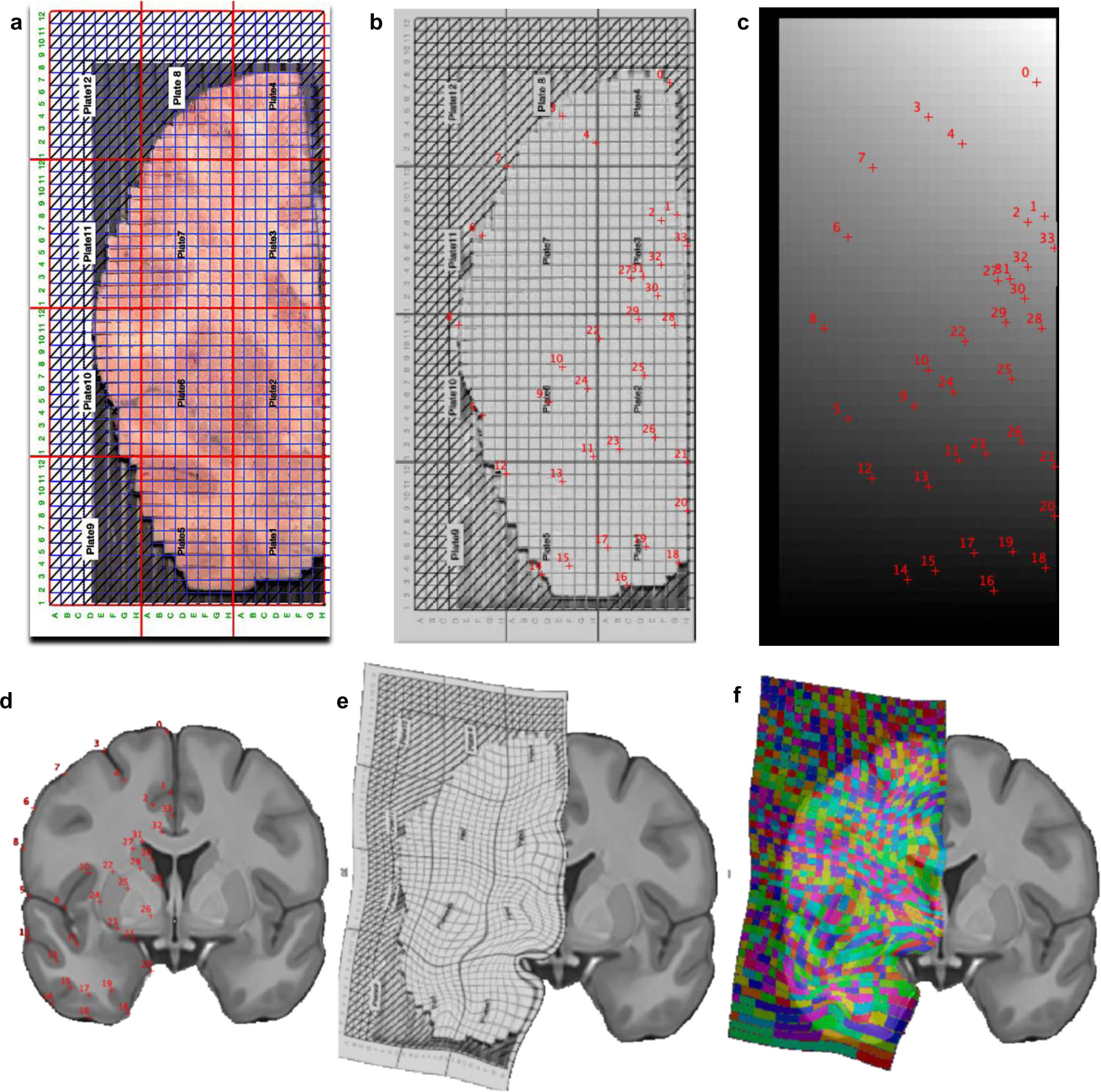
Morphing the partitioned brain slice into Montreal Neurological Institute (MNI) space. **a,** Image of the partitioned brain slice with an overlaid milling grid. **b,** Conversion of the brain slice into Neuroimaging Informatics Technology Initiative (NIfTI) format and identification of 34 anatomical landmarks. **c,** Manual creation of NIfTI grid corresponding to the milling grid with the same coordinates. **d,** Identification of the anatomical landmarks as on *b* in the magnetic resonance imaging stereotaxic space (MNI152). **e,** Warping and deformation of the coordinates from *b* to *d*. **f,** Application of this deformation to *c* displayed in color onto the MNI152 for visual convenience. The following 34 anatomical landmarks were identified: top of the interhemispheric fissure (0), surface (1) and deepest (2) point of the cingulate sulcus, surface (3) and deepest point (4) of the superior frontal sulcus (6), middle of the middle frontal gyrus (7), surface of the inferior frontal sulcus, middle surface of the inferior frontal gyrus (8), surface (5) and deepest point (9) of the lateral fissure, highest (10) and lowest (11) point of the circular insular sulcus, surface (12) and deepest point (13) of the superior temporal sulcus, surface (14) and deepest point (15) of the inferior temporal sulcus, surface (16) and deepest point (17) of the accessory temporopolar sulcus, surface (18) and deepest point (19) of the entorhinal sulcus, middle of the entorhinal cortex (20), most medial points of the amygdala (21), superior (22), inferior (23), lateral (24), and medial (25) putamen, most medial internal globus pallidus inferior (26), superior (27), inferior (28) lateral (29) and medial (30) caudate, highest point of the lateral ventricles (31), deepest point of the callosal sulcus (32) and middle of the cingulum gyrus (33).

**Extended Data Fig. 9.**
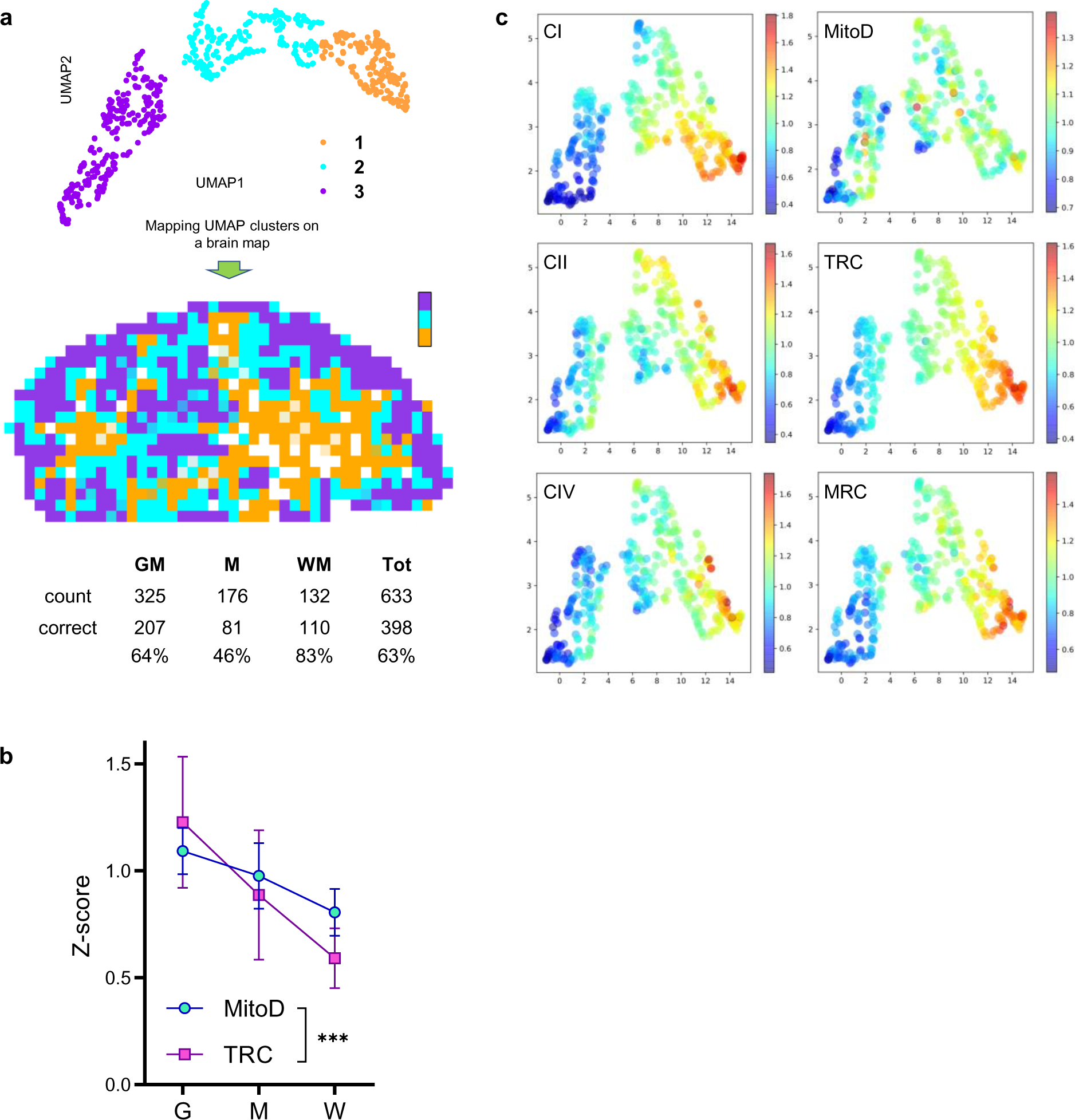
Clustering of brain voxels by similarities of mitochondrial density and OxPhos activities. **a,** Three visually defined UMAP clusters (upper) obtained after dimensionality reduction of mitochondrial features of all brain voxels (Fig. 2g) were used to predict whether voxels originated from GM, WM or a mixed sample (middle). The table lists prediction accuracy with the number of voxels of each type and the fractions that were identified correctly. Χ^2^(2, N=1031)=10.3, p<0.01. **b,** Both MitoD and TRC values were higher in GM than the WM voxels (same data as on Fig 2h are used; mean±SD), but the TRC decreases more than MitoD. ***-p<0.0001 by 2-way ANOVA (F_1,1260_=18.67). **c,** Distribution of z-score values of each mitochondrial feature on the UMAP plot of GM voxels only (Fig. 2i).

**Extended Data Fig. 10.**
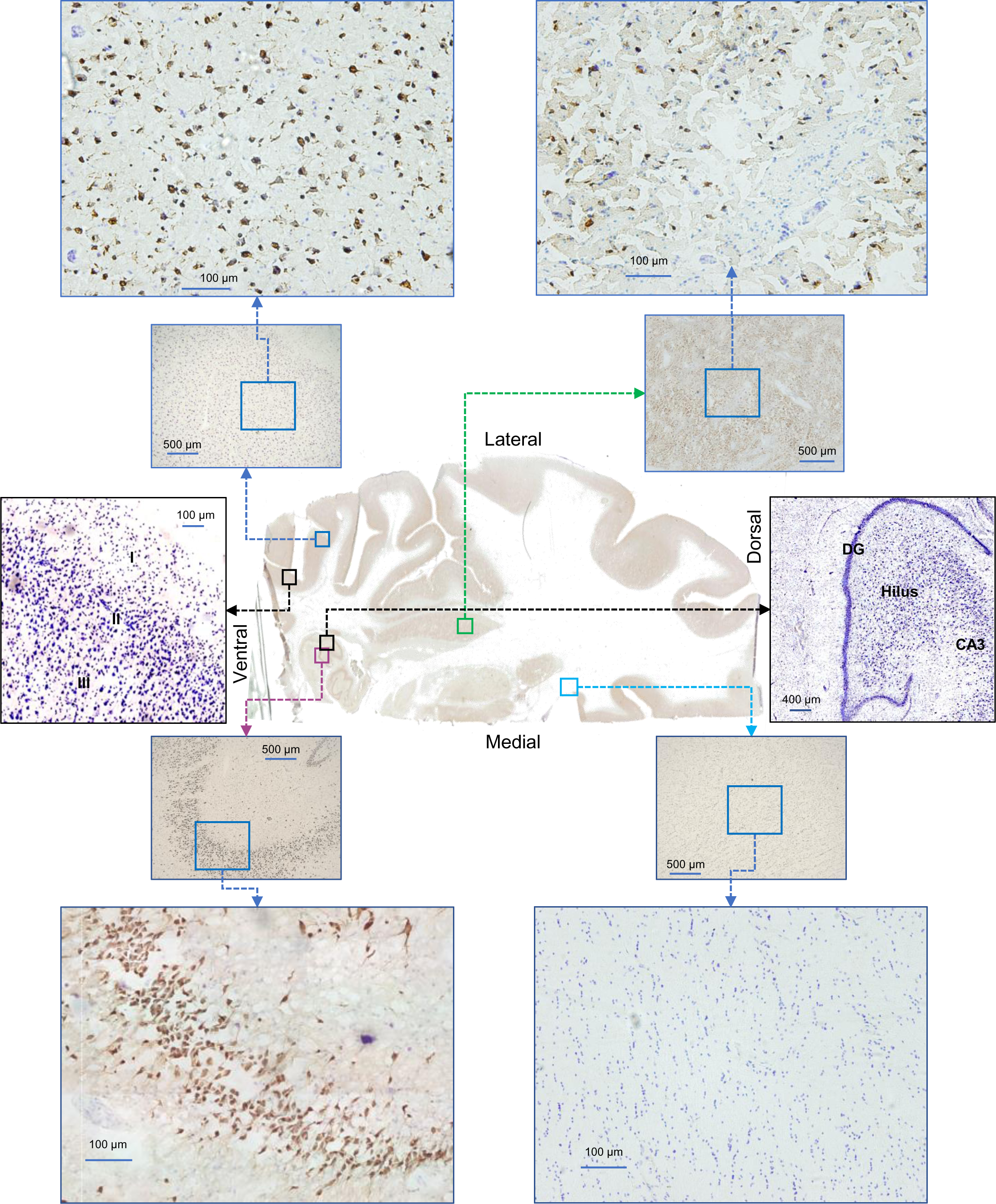
Histological evaluation of brain areas used for snRNAseq analysis. Staining of 50 µm sections obtained after brain voxelization (see Extended Data Fig 1) with Nissl stain (blue, neurons and glial cells) and NeuN (brown, neuronal cells). Two panels in black frames show Nissl-only stained sections with annotated cortical (layers I-II) and hippocampal (dentate gyrus, hilus and CA3) areas.

**Extended Data Fig. 11.**
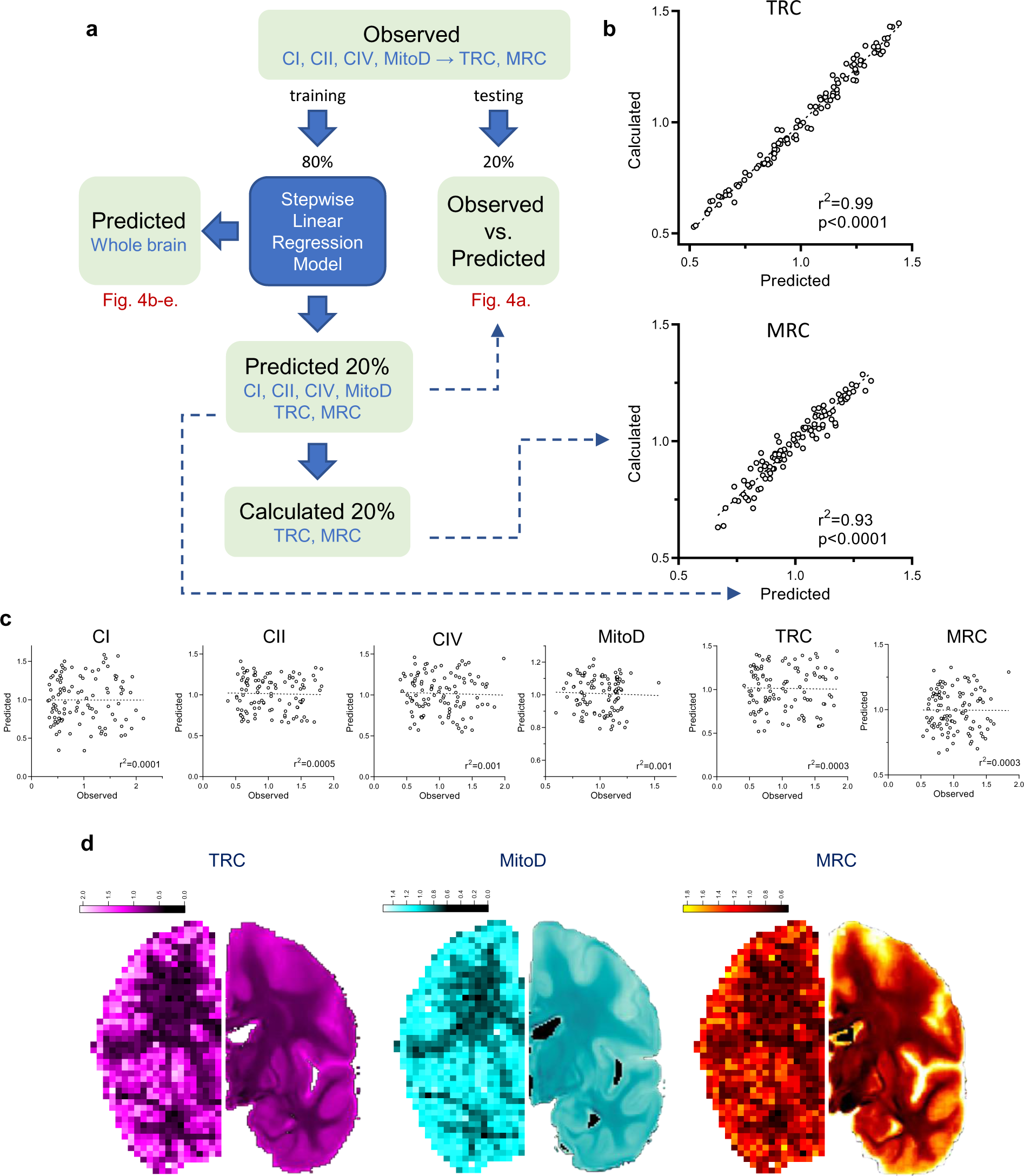
Workflow of data processing with stepwise linear regression model. **a,** Voxels that had all 6 mitochondrial parameters (4 independent and 2 derived) and with a sum of GM and WM probabilities more than 70% (n = 539 ‘observed’ voxels) were randomly split into 80% learning (n = 431) and 20% testing (n = 108) datasets. A model was built by predicting each of the 6 mitochondrial values in the learning dataset using stepwise backward linear regression of neuroimaging values (Extended Data Table 1). The model was then applied to the testing dataset to verify predictions accuracy. Correlation between observed and predicted values of the testing dataset are shown on Fig. 4a. Next, the predictive models were extended to all the brain voxel of the MNI space to produce a whole brain estimated map for each of the mitochondrial values at 1 mm^3^ resolution (Fig. 4b-e). **b,** Correlation between TRC and MRC values either predicted by the model or calculated from predicted CI, CII, CIV and MitoD values (see formulas on Fig. 2c). Strong correlation between predicted and calculated values for TRC and MRC shows the robustness of the model as it’s able to accurately predict derived parameters (TRC and MRC) without the knowledge of the independent readouts they were derived from (CI, CII, CIV and MitoD). **c,** Scatterplots of 20% out-of-sample prediction of mitochondrial profiles (same data as on Fig. 4a) after the pairing between observed and predicted values was scrambled. Slopes of all linear regressions are not different from zero (p>0.7; Pearson’s r^2^ is shown on each graph). **d,** Predicted (upper row) vs. observed (lower row, same data as on Fig 2d-f) maps of values for TRC, MitoD and MRC from the same coronal plane. Scale bars are in z-score units.

**Extended Data Fig. 12.**
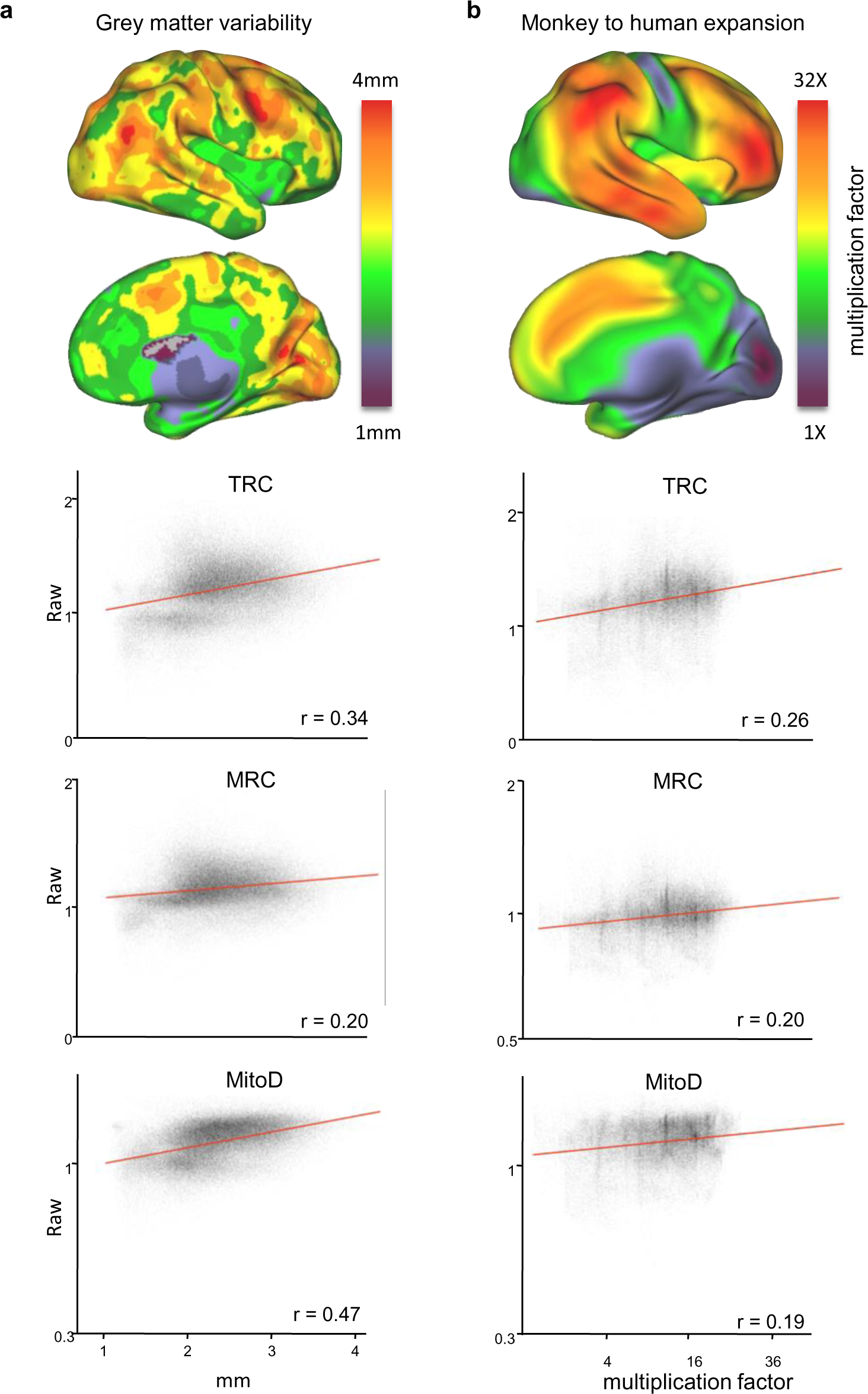
Higher mitochondrial activity in phylogenetically younger brain areas. **a-b,** Comparison between predicted values and indirect measures of brain evolution such as grey matter variability (a) and monkey-to-humans areal expansion (b).

**Extended Data Table 1.** List of neuroimaging metrics and their standardized beta coefficient relationship with the mitochondrial features.

|  | Abbreviation | MRI metric | CI | CII | CIV | MitoD | TRC | MRC |
| --- | --- | --- | --- | --- | --- | --- | --- | --- |
| Diffusion MRI | AD | axial water diffusivity |  | .242 |  |  | .246 |  |
|  | FA | white matter density |  |  | .681 | .820 |  |  |
|  | RD | radial water diffusivity | -.365 |  | .439 | .632 |  | -.175 |
|  | StreDensity | streamlines density |  | -.184 |  |  |  |  |
|  | ICVF | intra-cellular volume fraction | -.371 |  |  |  |  |  |
|  | ISOCF | extra-cellular volume fraction |  | -.158 | -.438 | -.445 | -.330 |  |
|  | MD | mean water diffusivity |  |  |  |  |  |  |
|  | OD | orientation dispersion index (neurite complexity) | .363 | .323 | .628 | .499 | .481 | .334 |
| Structural MRI | T1W | T1w imaging |  |  |  |  |  |  |
|  | T2W | T2w imaging | -.245 | -.304 | -.458 | -.410 | -.330 | -.244 |
|  | T1w/T2w | T1w/T2w ratio | -.656 | -.580 | -.723 | -.622 | -.617 | -.593 |
|  | FLAIR | imaging approach to see the anatomy of the brain |  | -.221 |  | -.339 |  |  |
|  | CT | cortical thickness |  |  |  |  |  |  |
|  | inner_CSA | local surface for inner cortical ribbon |  |  |  |  |  |  |
|  | plial_CSA | local surface for outer cortical ribbon |  |  |  |  |  |  |
|  | GM | probability of gray matter | -1.038 |  | -1.382 |  | -1.177 | -.939 |
|  | WM | probability of white matter | -1.082 |  | -1.263 | -.312 | -1.040 | -.906 |
| Functional MRI | Max_activity | maximum bold derived from fMRI |  |  | .380 |  | .333 | .501 |
|  | Reho | regional homogeneity |  | .145 |  | .135 |  |  |
|  | Entropy | synaptic complexity derived from fMRI |  |  | -.386 |  | -.365 | -.560 |
|  | ALFF | amplitude of Low-Frequency Fluctuation |  |  |  |  |  |  |
|  | fALFF | ratio between low and and high frequency fluctuations |  |  |  |  |  |  |

