## Supplementary Files1-4 and video1 for "A Human Brain Map of Mitochondrial Respiratory Capacity and Diversity": Supplementary File1 - Markdown of RNAseq analysis.html

Supplementary File 1 - Markdown of RNAseq analysis


Code 

- Show All Code
- Hide All Code

### Supplementary File 1 - Markdown of RNAseq analysis

#### Supplementary File 1 - Markdown of RNAseq analysis

- Data pre-processing
  - Quality control measures
  - Selecting cells
  - Normalization, feature selection, and scaling
  - Dimensionality reduction
  - Doublet removal
  - Harmony integration
- Cluster identification
  - UMAP, neighbors and clustering
  - Azimuth prediction
- Assigning cell type identity to clusters
  - UMAP annotated
  - New UMAP annotated
  - Marker expression
  - Cell type proportions
- Data downsampling
- Mitochondrial analysis
- R session
- References

Compiled: February 16, 2024

This markdown file links the code to the figures shown in Figure 3, as well as additional information (tables, interactive plots) supporting the analysis. Code and images expand when clicked on. The original markdown file can be found on github https://github.com/mitopsychobio. The raw data can be downloaded from -TBD-. Single nuclei RNA sequencing data was generated on a 10x platform and analyzed using the Seurat package^4^.

```
options(mc.cores = parallel::detectCores() - 1)
library(tidyverse)
library(Seurat)
library(ComplexHeatmap)
library(RColorBrewer)
library(DoubletFinder)
library(Azimuth)
library(fontawesome)
library(parallel)


col_cells =  c(
  "unspecific" = "grey",
  "Inhibitory" = "#FF7F00",# 
  "Excitatory" = "#007200",
  "OPC" = "#A6CEE3", 
  "Oligodendrocytes" = "#1F78B4",
  "Astrocytes" = "#E15A57",
  "Microglia" ="#FFFF99",
  "VLMCs" =  "#6A3D9A",
  "Pericytes" = "#D0BEDA",
  "Endothelial" = "#B789DA"
  )

col_vox = list("Cortex" = "#4F5088",
            "Hippocampus" = "#A5458D",
            "Putamen" =  "#5D9E4F",
            "WhiteMatter"= "#4E77BC")

theme_density <- theme(axis.title = element_text(size = 20),
                       axis.text = element_text(size = 16),
                       legend.text = element_text(size = 20),
                       legend.title = element_text(size = 20),
                       plot.title = element_blank(),
                       legend.position = "right")
theme_umap <- theme(axis.title = element_text(size = 20),
                    axis.text = element_text(size = 18),
                    legend.text = element_text(size = 20),
                    plot.title = element_blank())


theme_predicted <- theme(title = element_text(size = 12, face = "plain"),
        legend.text  = element_text(size = 10))

theme_dotplot <- theme(axis.text.x = element_text(angle = 45, hjust =1,size = 8),
        axis.text.y = element_text(size = 8),
        axis.title = element_blank() ,
        strip.text = element_text(angle = 90, hjust =1, size = 7))
```

---

### Data pre-processing

**Import data** from cellbender^5^

The FASTQ files were processed with the 10x software Cell Ranger (QCs on github) and further processed using cellbender^5^ to remove debris and doublets (not part of this markdown file).

```
vox1 <- readRDS(paste0(mainpath, "/CellBender/cellbender_vox1.rds"))
vox2 <- readRDS(paste0(mainpath, "/CellBender/cellbender_vox2.rds"))
vox3 <- readRDS(paste0(mainpath, "/CellBender/cellbender_vox3.rds"))
vox4 <- readRDS(paste0(mainpath, "/CellBender/cellbender_vox4.rds"))
data <- merge(vox1, y = c(vox2, vox3, vox4), add.cell.ids = c("MTG", "HC", "PT", "WM"), project = "MitoBrainMap")

data$mitoRatio <- PercentageFeatureSet(data, pattern = "^MT-")
data$log10GenesPerUMI <- log10(data$nFeature_RNA) / log10(data$nCount_RNA)

metadata <- data@meta.data
metadata <- metadata %>%
  mutate(Brainregion = case_when(
    orig.ident == "voxel1" ~ "Cortex",
    orig.ident == "voxel2" ~ "Hippocampus",
    orig.ident == "voxel3" ~ "Putamen",
    orig.ident == "voxel4" ~ "WhiteMatter",
  )) %>%
  dplyr::rename(
    nUMI = nCount_RNA,
    nGene = nFeature_RNA)
```

#### Quality control measures

##### UMIs per cell

```
metadata %>% 
  ggplot(aes(color=Brainregion, x=nUMI, fill= Brainregion)) + 
  geom_density(alpha = 0.2) + 
  scale_x_log10() + 
  scale_color_manual(values = unlist(col_vox)) + 
  scale_fill_manual(values = unlist(col_vox)) + 
  theme_classic() +
  ylab("Cell density") +
  geom_vline(xintercept = 500) + 
  theme_density
```

##### Genes per cell

```
metadata %>% 
  ggplot(aes(color=Brainregion, x=nGene, fill= Brainregion)) + 
  geom_density(alpha = 0.2) + 
  theme_classic() +
  scale_x_log10() + 
  geom_vline(xintercept = 200) +
  geom_vline(xintercept = 5000) +
  ylab("Cell density") +
  scale_color_manual(values = unlist(col_vox)) + 
  scale_fill_manual(values = unlist(col_vox)) + 
  theme_density
```

##### Genes per UMI

```
metadata %>%
  ggplot(aes(x=log10GenesPerUMI, color = Brainregion, fill=Brainregion)) +
  geom_density(alpha = 0.2) +
  theme_classic() +
  geom_vline(xintercept = 0.8) +
  ylab("Cell density") +
  scale_color_manual(values = unlist(col_vox)) + 
  scale_fill_manual(values = unlist(col_vox)) + 
  theme_density
```

##### Mitochondrial genes

```
metadata %>% 
  mutate(xintercept = case_when(
    orig.ident == "voxel1" ~ 6,
    orig.ident == "voxel2" ~ 8,
    orig.ident == "voxel3" ~ 11,
    orig.ident == "voxel4" ~ 4.5,
  )) %>%
  mutate(xintercept = as.numeric(xintercept))%>%
  ggplot(aes(color=Brainregion, x=mitoRatio, fill=Brainregion)) + 
  geom_density(alpha = 0.2) + 
  theme_classic() +
  ylab("Cell density") +
  xlab("% mitochondrial counts") +
  scale_color_manual(values = unlist(col_vox)) + 
  scale_fill_manual(values = unlist(col_vox)) + 
  theme_density+
  xlim(0,20)
```

##### Summary

```
metadata %>% 
  ggplot(aes(x=nUMI, y=nGene, color=mitoRatio)) + 
  geom_point() + 
  scale_colour_gradient(low = "gray90", high = "black") +
  stat_smooth(method=lm) +
  scale_x_log10() + 
  scale_y_log10() + 
  theme_classic() +
  facet_wrap(~Brainregion) +
  theme(axis.title = element_text(size = 20),
        axis.text = element_text(size = 14),
        legend.text = element_text(size = 20),
        legend.title = element_text(size = 20),
        plot.title = element_blank(),
        strip.text = element_text(size =20),
        legend.position = "right")
```

#### Selecting cells

Cells were selected based on the following QC metrics:

- 200 - 5000 Genes
- >500 counts
- <11% mitochondrial genes
- Removal of ribosomal genes

```
vox1 <- PercentageFeatureSet(vox1, pattern = "^MT-", col.name = 'percent.mt')
vox1 <- subset(vox1, subset = 
                 nFeature_RNA > 200 & nFeature_RNA < 5000 & 
                 nCount_RNA > 500 & 
                 percent.mt < 11)
counts <- GetAssayData(vox1, assay = "RNA")
mt <- counts[-grep("^MT-", rownames(counts)),]
rp <- counts[-grep("^RP[SL]", rownames(counts)),]
vox1 <- subset(vox1, features = rownames(rp))

vox2 <- PercentageFeatureSet(vox2, pattern = "^MT-", col.name = 'percent.mt')
vox2 <- subset(vox2, subset = 
                 nFeature_RNA > 200 & nFeature_RNA < 5000 & 
                 nCount_RNA > 500 & 
                 percent.mt < 11)
counts <- GetAssayData(vox2, assay = "RNA")
mt <- counts[-grep("^MT-", rownames(counts)),]
rp <- counts[-grep("^RP[SL]", rownames(counts)),]
vox2 <- subset(vox2, features = rownames(rp))

vox3 <- PercentageFeatureSet(vox3, pattern = "^MT-", col.name = 'percent.mt')
vox3 <- subset(vox3, subset = 
                 nFeature_RNA > 200 & nFeature_RNA < 5000 & 
                 nCount_RNA > 500 &
                 percent.mt < 11)
counts <- GetAssayData(vox3, assay = "RNA")
mt <- counts[-grep("^MT-", rownames(counts)),]
rp <- counts[-grep("^RP[SL]", rownames(counts)),]
vox3 <- subset(vox3, features = rownames(rp))

vox4 <- PercentageFeatureSet(vox4, pattern = "^MT-", col.name = 'percent.mt')
vox4 <- subset(vox4, subset = 
                 nFeature_RNA > 200 & nFeature_RNA < 5000 & 
                 nCount_RNA > 500 & 
                 percent.mt < 11)
counts <- GetAssayData(vox4, assay = "RNA")
mt <- counts[-grep("^MT-", rownames(counts)),]
rp <- counts[-grep("^RP[SL]", rownames(counts)),]
vox4 <- subset(vox4, features = rownames(rp))
```

For subsequent analyses, all 4 seurat objects were combined and processed together.

```
data <- merge(vox1, y = c(vox2, vox3, vox4), add.cell.ids = c("MTG", "HC", "PT", "WM"), project = "MitoBrainMap")
rm(vox1, vox2, vox3, vox4, dataloc, counts, mt, rp)

# voxel_annotations <- list(
#  'voxel1' = 'Cortex',
#   'voxel2' = 'Hippocampus',
#   'voxel3' = 'Putamen',
#   'voxel4' = 'WhiteMatter'
# )
# data$Brainregion <- unlist(voxel_annotations[data$orig.ident])
data$Brainregion <- NA
data$Brainregion[data$orig.ident == "voxel1"] <- 'Cortex'
data$Brainregion[data$orig.ident == "voxel2"] <- 'Hippocampus'
data$Brainregion[data$orig.ident == "voxel3"] <- 'Putamen'
data$Brainregion[data$orig.ident == "voxel4"] <- 'WhiteMatter'
```

#### Normalization, feature selection, and scaling

The data was log-normalized using the NormalizeData() function, and 3000 highly variable features between the cells were mapped. Next, the data was scaled and centered around 0 (linear transformation) using the previously identified 3000 features.

```
data <- NormalizeData(data, normalization.method = "LogNormalize", scale.factor = 1e4)
data <- FindVariableFeatures(data, selection.method = 'vst', nfeatures = 3000)
data <- ScaleData(data)
```

#### Dimensionality reduction

##### PCA

To determine how many principal components (PCs) will be used in downstream analyses, the standard deviations of PCs were visualized using an ElbowPlot. Next, we used a quantitative method as suggested by https://github.com/hbctraining/scRNA-seq. First, we determined the PCs that cumulatively contribute 90% of the standard deviation (PCs 1 to 41), with less than 5% variation associated with the PC (PCs 1, 2, and 3). We further matched the PC where the percent change in variation between consecutive PCs is less than 0.1% (PC 21). The minimum value of both thresholds determined the maximal PC for downstream analyses (21 PCs in total).

```
data <- RunPCA(data, features = VariableFeatures(object = data))
ElbowPlot(data, ndims = 40, reduction = "pca")
```

```
ggsave(paste0(mainpath,"/", Sys.Date(), "/Plots/PCA_elbow.png"), width = 5, height =4.5)
pct <- data[["pca"]]@stdev / sum(data[["pca"]]@stdev) * 100
cumu <- cumsum(pct)
co1 <- which(cumu > 90 & pct < 5)[1]
co2 <- sort(which((pct[1:length(pct) - 1] - pct[2:length(pct)]) > 0.1), decreasing = T)[1] + 1
pcs <- min(co1, co2)
```

##### UMAP

UMAP of the non-integrated data showing clear separation between the datasets.

```
temp <- RunUMAP(data, dims = 1:pcs, seed.use = 953)
DimPlot(temp,label=F, label.size = 5,reduction = "umap", 
        group.by ="Brainregion", cols = unlist(col_vox)) +
  guides(color = guide_legend(override.aes = list(size = 6))) +
  theme_umap
```

```
ggsave(paste0("Plots/", Sys.Date(), "UMAP1_origIdent.png"), width = 7.5, height = 6.3)
rm(temp)
```

#### Doublet removal

In addition to cellbender, the function DoubletFinder^6^ was used to identify and remove cell doublets. We assumed a 7.5% doublet formation rate. The pK value (required by the function) was determined to be at 5e-04 (see figure below).

```
bcmvn %>%
  ggplot(aes(x = pK, y = MeanBC)) +
  geom_point() +
  theme(axis.text.x = element_text(angle = 45, hjust = 1))
```

```
ggsave(paste0(mainpath,"/", Sys.Date(), "/Plots/DF_bcmvn.png"), width = 6, height =4.5)
nExp_poi <- round(0.075*nrow(data_df@meta.data))  ## Assuming 7.5% doublet formation rate - tailor for your dataset
data_df <- doubletFinder_v3(data_df, PCs = 1:pcs, pN = 0.25, pK = 5e-04, nExp = nExp_poi, reuse.pANN = FALSE, sct = FALSE)
colnames(data_df@meta.data) <- c("orig.ident",
                                      "nCount_RNA",
                                      "nFeature_RNA",
                                      "percent.mt",
                                      "Brainregion",
                                      "pANN",
                                      "DF_class")
```

The following UMAP shows singlets and doublets. All doublets were filtered out afterwards.

```
tmp <- RunUMAP(data_df, dims = 1:pcs, seed.use = 4321)
DimPlot(tmp,label=F, label.size = 5, reduction = "umap",
        group.by ="DF_class", cols = c("coral3","darkcyan")) +
  guides(color = guide_legend(override.aes = list(size = 6))) +
  theme_umap
```

```
ggsave(paste0(mainpath,"/", Sys.Date(), "/Plots/UMAP2_DoubledFinder.png"), width = 5, height =4.5)
data_singlets <-data_df %>% subset(subset = DF_class == "Singlet")
rm(tmp)
```

#### Harmony integration

As previously shown, UMAP revealed clustering by dataset. Hence, the data was integrated using harmony^7^ prior to cell type identification and clustering.

```
set.seed(2)
data_harmony <- harmony::RunHarmony(data_singlets,"orig.ident", plot_convergence = TRUE, num.cores = detectCores() - 1)
```

```
rm(data, data_singlets)
```

---

### Cluster identification

#### UMAP, neighbors and clustering

- Reduction = harmony
- Number of PCs: 21
- Resolution = 0.2

```
data <- data_harmony
Idents(data) <- "Brainregion"

data <- data %>%
  RunUMAP(reduction = "harmony", dims = 1:pcs, verbose = F, seed.use = 8) %>% #123 #1
  FindNeighbors(reduction = "harmony", dims = 1:pcs) %>%
  FindClusters(resolution =0.2, random.seed = 2)
```

```
## Modularity Optimizer version 1.3.0 by Ludo Waltman and Nees Jan van Eck
## 
## Number of nodes: 48793
## Number of edges: 1542315
## 
## Running Louvain algorithm...
## Maximum modularity in 10 random starts: 0.9521
## Number of communities: 12
## Elapsed time: 8 seconds
```

UMAP of the integrated data with clusters:

```
DimPlot(data,label=T, reduction = "umap")+
  NoLegend() +
  theme_umap
```

```
ggsave(paste0(mainpath,"/", Sys.Date(), "/Plots/UMAP3_clusters.png"), width = 5, height =4.5)
```

##### Cells per cluster

```
cells_per_cluster <- as.data.frame.matrix(
  table(data@meta.data[["RNA_snn_res.0.2"]],
        data@meta.data[["Brainregion"]])) %>%
  rownames_to_column("Cluster") %>%
  mutate(Cluster = as.numeric(Cluster)) %>%
  arrange(Cluster)

kableExtra::kable(cells_per_cluster)
```

| Cluster | Cortex | Hippocampus | Putamen | WhiteMatter |
| --- | --- | --- | --- | --- |
| 0 | 5573 | 7591 | 670 | 2444 |
| 1 | 502 | 1412 | 2868 | 3077 |
| 2 | 125 | 396 | 767 | 5969 |
| 3 | 3486 | 1657 | 1208 | 29 |
| 4 | 1217 | 1419 | 477 | 500 |
| 5 | 299 | 775 | 113 | 436 |
| 6 | 436 | 439 | 256 | 172 |
| 7 | 461 | 360 | 237 | 123 |
| 8 | 1021 | 105 | 24 | 1 |
| 9 | 947 | 192 | 0 | 0 |
| 10 | 353 | 172 | 172 | 32 |
| 11 | 98 | 117 | 54 | 11 |

#### Azimuth prediction

**Azimuth reference dataset:** Human motorcortex^8^. The code for the prediction was downloaded from Azimuth (https://app.azimuth.hubmapconsortium.org/app/human-motorcortex).

```
reference <- Azimuth:::LoadReference(path = "/Users/annamonzel/Desktop/R/R_MitoBrainMap/Run15_Final_Markdown", seconds = 10L)
query <- data
if (!all(c("nCount_RNA", "nFeature_RNA") %in% c(colnames(x = query[[]])))) {
    calcn <- as.data.frame(x = Seurat:::CalcN(object = query))
    colnames(x = calcn) <- paste(
      colnames(x = calcn),
      "RNA",
      sep = '_'
    )
    query <- AddMetaData(
      object = query,
      metadata = calcn
    )
    rm(calcn)
}

# Calculate percent mitochondrial genes if the query contains genes
# matching the regular expression "^MT-"
if (any(grepl(pattern = '^MT-', x = rownames(x = query)))) {
  query <- PercentageFeatureSet(
    object = query,
    pattern = '^MT-',
    col.name = 'percent.mt',
    assay = "RNA"
  )
}

# Filter cells based on the thresholds for nCount_RNA and nFeature_RNA
# you set in the app
cells.use <- query[["nCount_RNA", drop = TRUE]] <= 26235 &
  query[["nCount_RNA", drop = TRUE]] >= 485 &
  query[["nFeature_RNA", drop = TRUE]] <= 4984 &
  query[["nFeature_RNA", drop = TRUE]] >= 207

# If the query contains mitochondrial genes, filter cells based on the
# thresholds for percent.mt you set in the app
if ("percent.mt" %in% c(colnames(x = query[[]]))) {
  cells.use <- cells.use & (query[["percent.mt", drop = TRUE]] <= 12 &
    query[["percent.mt", drop = TRUE]] >= 0)
}

# Remove filtered cells from the query
query <- query[, cells.use]

# Preprocess with SCTransform
query <- SCTransform(
  object = query,
  assay = "RNA",
  new.assay.name = "refAssay",
  residual.features = rownames(x = reference$map),
  reference.SCT.model = reference$map[["refAssay"]]@SCTModel.list$refmodel,
  method = 'glmGamPoi',
  ncells = 2000,
  n_genes = 2000,
  do.correct.umi = FALSE,
  do.scale = FALSE,
  do.center = TRUE, verbose = F
)

# Find anchors between query and reference
anchors <- FindTransferAnchors(
  reference = reference$map,
  query = query,
  k.filter = NA,
  reference.neighbors = "refdr.annoy.neighbors",
  reference.assay = "refAssay",
  query.assay = "refAssay",
  reference.reduction = "refDR",
  normalization.method = "SCT",
  features = intersect(rownames(x = reference$map), VariableFeatures(object = query)),
  dims = 1:50,
  n.trees = 20,
  mapping.score.k = 100
)

# Transfer cell type labels and impute protein expression
#
# Transferred labels are in metadata columns named "predicted.*"
# The maximum prediction score is in a metadata column named "predicted.*.score"
# The prediction scores for each class are in an assay named "prediction.score.*"
# The imputed assay is named "impADT" if computed

refdata <- lapply(X = c("subclass", "class", "cluster"), function(x) {
  reference$map[[x, drop = TRUE]]
})
names(x = refdata) <- c("subclass", "class", "cluster")
if (FALSE) {
  refdata[["impADT"]] <- GetAssayData(
    object = reference$map[['ADT']],
    slot = 'data'
  )
}
query <- TransferData(
  reference = reference$map,
  query = query,
  dims = 1:50,
  anchorset = anchors,
  refdata = refdata,
  n.trees = 20,
  store.weights = TRUE, verbose = F
)

# Calculate the embeddings of the query data on the reference SPCA
query <- IntegrateEmbeddings(
  anchorset = anchors,
  reference = reference$map,
  query = query,
  reductions = "pcaproject",
  reuse.weights.matrix = TRUE,
  verbose = F
)

# Calculate the query neighbors in the reference
# with respect to the integrated embeddings
query[["query_ref.nn"]] <- FindNeighbors(
  object = Embeddings(reference$map[["refDR"]]),
  query = Embeddings(query[["integrated_dr"]]),
  return.neighbor = TRUE,
  l2.norm = TRUE
)

# The reference used in the app is downsampled compared to the reference on which
# the UMAP model was computed. This step, using the helper function NNTransform,
# corrects the Neighbors to account for the downsampling.
query <- Azimuth:::NNTransform(
  object = query,
  meta.data = reference$map[[]]
)

# Project the query to the reference UMAP.
query[["proj.umap"]] <- RunUMAP(
  object = query[["query_ref.nn"]],
  reduction.model = reference$map[["refUMAP"]],
  reduction.key = 'UMAP_',verbose = F
)


# Calculate mapping score and add to metadata
query <- AddMetaData(
  object = query,
  metadata = MappingScore(anchors = anchors),
  col.name = "mapping.score"
)
```

##### Predicted class and subclass

###### Original Identity

```
DimPlot(object = query, reduction = "umap", group.by = "Brainregion", label = F, 
        cols = unlist(col_vox)) +
  guides(color = guide_legend(override.aes = list(size = 6))) +
  labs(title = "Color-coded by dataset") +
  theme_umap
```

###### Predicted class

```
id <- c("class")[1]
predicted.id <- paste0("predicted.", id)
DimPlot(object = query, reduction = "umap", group.by = predicted.id, label = TRUE, label.size = 3, label.box = TRUE ) +
  labs(title = "Predicted class") +
  theme_predicted
```

###### Predicted subclass

```
id <- c("subclass")[1]
predicted.id <- paste0("predicted.", id)
DimPlot(object = query, reduction = "umap", group.by = predicted.id, label = TRUE, label.size = 3, label.box = TRUE ) +
  labs(title = "Predicted subclass") +
  theme_predicted
```

Cells from the white matter sample (voxel 4) were all predicted as oligodendrocytes and temporarily excluded for cell type identification.

```
Idents(query) <- "orig.ident"
query_sub <- subset(query, idents = c("voxel4"), invert = T)
```

##### Scores of predicted cell types

###### Glutamatergic

```
id <- c("class")[1]
predicted.id <- paste0("predicted.", id)
FeaturePlot(object = query_sub, features = "Glutamatergic", reduction = "umap") +
  labs(title = "Glutamatergic") +
  theme_predicted
```

```
VlnPlot(object = query_sub, features = "Glutamatergic", group.by = predicted.id) + NoLegend() +
  labs(title = "Glutamatergic") +
  theme_predicted
```

###### GABAergic

```
FeaturePlot(object = query_sub, features = "GABAergic", reduction = "umap")+
  labs(title = "GABAergic") +
  theme_predicted
```

```
VlnPlot(object = query_sub, features = "GABAergic", group.by = predicted.id) + NoLegend()+
  labs(title = "GABAergic") +
  theme_predicted
```

###### Astrocytes

```
id <- c("subclass")[1]
predicted.id <- paste0("predicted.", id)

FeaturePlot(object = query_sub, features = "Astro", reduction = "umap")+
  labs(title = "Astrocytes") +
  theme_predicted
```

```
VlnPlot(object = query_sub, features = "Astro", group.by = predicted.id) + NoLegend()+
  labs(title = "Astrocytes") +
  theme_predicted
```

###### Oligodendrocytes

```
FeaturePlot(object = query_sub, features = "Oligo", reduction = "umap")+
  labs(title = "Oligodendrocytes") +
  theme_predicted
```

```
VlnPlot(object = query_sub, features = "Oligo", group.by = predicted.id) + NoLegend()+
  labs(title = "Oligodendrocytes") +
  theme_predicted
```

###### Microglia

```
FeaturePlot(object = query_sub, features = "Micro-PVM", reduction = "umap")+
  labs(title = "Microglia") +
  theme_predicted
```

```
VlnPlot(object = query_sub, features = "Micro-PVM", group.by = predicted.id) + NoLegend()+
  labs(title = "Microglia") +
  theme_predicted
```

###### OPCs

```
FeaturePlot(object = query_sub, features = "OPC", reduction = "umap")+
  labs(title = "OPCs") +
  theme_predicted
```

```
VlnPlot(object = query_sub, features = "OPC", group.by = predicted.id) + NoLegend()+
  labs(title = "OPCs") +
  theme_predicted
```

###### VLMCs

```
FeaturePlot(object = query_sub, features = "VLMC", reduction = "umap")+
  labs(title = "VLMCs") +
  theme_predicted
```

```
VlnPlot(object = query_sub, features = "VLMC", group.by = predicted.id) + NoLegend()+
  labs(title = "VLMCs") +
  theme_predicted
```

###### Endothelial

```
FeaturePlot(object = query_sub, features = "Endo", reduction = "umap")+
  labs(title = "Endothelial") +
  theme_predicted
```

```
VlnPlot(object = query_sub, features = "Endo", group.by = predicted.id) + NoLegend()+
  labs(title = "Endothelial") +
  theme_predicted
```

###### Original clusters (for comparison)

```
DimPlot(data,label=T, reduction = "umap")+
  NoLegend() +
  theme_umap
```

Clusters 0 and 3 have no clear predicted cell type class. We used a set of manually-curated markers to find the cell type identity. Moving forward, we added the previously excluded voxel 4 (white matter) back to the dataset.

##### Missing clusters

To identify the missing cell type identities, we derived a list of differentially expressed genes (DEGs) between the missing clusters. This list was inspected online using the cortical cell types database from the allen brain atlas https://celltypes.brain-map.org/rnaseq/^10^. We further visualized the expression of a manually-curated set of markers using dotplots.

###### UMAP subset

```
Idents(query_sub) <- "seurat_clusters"
query_missing <- subset(query_sub, idents = c("0", "3", "10"), invert = F)
id <- c("subclass")[1]
predicted.id <- paste0("predicted.", id)

DimPlot(object = query_missing, reduction = "umap", group.by = predicted.id, label = F)  +
  guides(color = guide_legend(override.aes = list(size = 6))) +
  theme_umap
```

###### Top20 DEGs

Top20 differentially expressed genes between the clusters

```
DefaultAssay(query_missing) <- "RNA"
markers <- FindAllMarkers(query_missing, only.pos = TRUE, min.pct = 0.25, logfc.threshold = 0.25)
top20 <- markers %>% group_by(cluster) %>% top_n(n = 20, wt = avg_log2FC)
top10 <- markers %>% group_by(cluster) %>% top_n(n = 10, wt = avg_log2FC)

kableExtra::kable(top20 %>%
                    select(cluster, gene) %>%
                    pivot_wider(names_from = "cluster", values_from = "gene"))
```

| 0 | 3 | 10 |
| --- | --- | --- |
| GFAP, OG…. | AHI1, DL…. | EBF1, RG…. |

###### Heatmap of DEGs

Heatmap of differentially expressed genes for each identity class

```
query_missing <- ScaleData(query_missing, features = top10$gene)
DoHeatmap(query_missing, features = top10$gene)
```

```
ggsave(paste0(mainpath,"/", Sys.Date(), "/Plots/Heatmap_Missing_Clusters_Top10DEGs.png"))
```

###### Dotplot marker expression

```
Idents(query) <- "seurat_clusters"
All_Marker <- list("Neurons" = c("RBFOX3"),
                   "Glut" = c("SLC17A6", "SLC17A7", "SLC17A8"),
                   "GABA" = c("GAD1","GAD2", "SST", "RELN"),
                  "Astro" = c("MEIS2",  "EIF1B",  "AQP4", "S100B", "SLC1A3","ALDH1L1"),
                  "OPC" = c( 'PDGFRA'),
                  "Oligo" = c("MOBP", "MOG", "MAG", "PLP1"),
                  "Microglia" = c("AIF1","TMEM119"),
                   "Endo" = c("CLDN5"),
                   "Peri" = c("PDGFRB", "RGS5")
)
DotPlot(query_missing, features = unlist(All_Marker), assay = "RNA", dot.scale = 3,  cols = "RdYlBu") +
  theme_classic() +
  theme_dotplot
```

```
ggsave(paste0(mainpath,"/", Sys.Date(), "/Plots/Dotplot_missing.png"), width = 7, height =3)
rm(query, query_sub, query_missing, data_df, data_harmony, refdata, reference, data_singlets)
```

**Cluster 0** shows a mixed expression of several markers, such as markers of astrocytes, oligodendrocytes, as well as mitochondrial genes. This suggests that cluster 0 is a population of hybrids of several compromised nuclei, and it will be excluded for downstream analyses. **Cluster 3** expresses markers indicative of glutamatergic neurons, despite background expression of other (non-neuronal) markers. It appears to contain a fraction of low quality nuclei, as some cells also express astrocytic and mitochondrial genes. Nevertheless, cluster 3 will be included in downstream analysis because it primarily has a neuronal identity, and most of the DEGs are higher expressed in glutamatergic neurons in other reference datasets. **Cluster 10** expresses markers of pericytes, previously not predicted in azimuth (missing in reference data).

```
## Going back to data (without predictions but with voxel 4)
Idents(data) <- "seurat_clusters"
data <- subset(data, idents = c("0"), invert = T)
DimPlot(object = data, reduction = "umap", group.by = "seurat_clusters", label = TRUE) +
  labs(title = "UMAP after cleaning")
```

---

### Assigning cell type identity to clusters

Clusters were annotated as follows:

```
cluster_annotations <- list(
  '0' = "unspecific",
  '1' = "Oligodendrocytes",
  '2' = "Oligodendrocytes",
  '3' = "Excitatory",
  '4' = "Astrocytes",
  '5' = "Endothelial",
  '6' = "Microglia",
  '7' = "OPC",
  '8' = "Inhibitory",
  '9' = "Excitatory",
  '10' = "Pericytes",
  '11' = "VLMCs"
)

table <- unlist(cluster_annotations)
table <- data.frame(Cluster = names(table), Annotation = table) %>%
  rownames_to_column("rowname") %>%
  select(-rowname)
kableExtra::kable(table)
```

| Cluster | Annotation |
| --- | --- |
| 0 | unspecific |
| 1 | Oligodendrocytes |
| 2 | Oligodendrocytes |
| 3 | Excitatory |
| 4 | Astrocytes |
| 5 | Endothelial |
| 6 | Microglia |
| 7 | OPC |
| 8 | Inhibitory |
| 9 | Excitatory |
| 10 | Pericytes |
| 11 | VLMCs |

#### UMAP annotated

```
data$CellType <- NA
data$CellType[data$seurat_clusters == 0] <- "unspecific"
data$CellType[data$seurat_clusters == 1] <- "Oligodendrocytes"
data$CellType[data$seurat_clusters == 2] <- "Oligodendrocytes"
data$CellType[data$seurat_clusters == 3] <- "Excitatory"
data$CellType[data$seurat_clusters == 4] <- "Astrocytes"
data$CellType[data$seurat_clusters == 5] <- "Endothelial"
data$CellType[data$seurat_clusters == 6] <- "Microglia"
data$CellType[data$seurat_clusters == 7] <- "OPC"
data$CellType[data$seurat_clusters == 8] <- "Inhibitory"
data$CellType[data$seurat_clusters == 9] <- "Excitatory"
data$CellType[data$seurat_clusters == 10] <- "Pericytes"
data$CellType[data$seurat_clusters == 11] <- "VLMCs"
data$CellType_cluster <- paste0(data$CellType, '-',data$seurat_clusters)
# cluster_annotations <- list(
#   '0' = "unspecific",
#   '1' = "Oligodendrocytes",
#   '2' = "Oligodendrocytes",
#   '3' = "Excitatory",
#   '4' = "Astrocytes",
#   '5' = "Endothelial",
#   '6' = "Microglia",
#   '7' = "OPC",
#   '8' = "Inhibitory",
#   '9' = "Excitatory",
#   '10' = "Pericytes",
#   '11' = "VLMCs"
# )
# data$CellType <- unlist(cluster_annotations[data$seurat_clusters])
# 
# 
# data$CellType_cluster <- paste0(data$CellType, '-',data$seurat_clusters)
# 
Idents(data) <- "CellType"

DimPlot(data,label=T, reduction = "umap", group.by = "CellType",
        cols = unlist(col_cells))+
  NoLegend() +
  guides(color = guide_legend(override.aes = list(size = 6))) +
  theme_umap
```

#### New UMAP annotated

We re-ran UMAP on the annotated and filtered dataset (for visualization purposes).

```
data <- data %>%
  RunUMAP(reduction = "harmony", dims = 1:pcs, verbose = F, seed.use = 3) 

DimPlot(data,label=T, reduction = "umap", group.by = "CellType",
        cols = unlist(col_cells))+
  NoLegend() +
  guides(color = guide_legend(override.aes = list(size = 6))) +
  theme_umap
```

```
ggsave(paste0(mainpath,"/", Sys.Date(), "/Plots/Figure3B_UMAP_annotated.png"), width = 5, height =4.5)


p <- DimPlot(data,label=F, reduction = "umap", group.by = "CellType",
        cols = unlist(col_cells))+
  NoLegend() +
  guides(color = guide_legend(override.aes = list(size = 6))) +
  theme_umap
ggsave(paste0(mainpath,"/", Sys.Date(), "/Plots/Figure3B_UMAP_blank.png"),p, width = 5, height =4.5)
```

#### Marker expression

##### Dotplot

```
DotPlot(data, features = unlist(All_Marker), assay = "RNA", dot.scale = 3,  cols = "RdYlBu") +
 theme_classic() +
  theme_dotplot
```

```
ggsave(paste0(mainpath,"/", Sys.Date(), "/Plots/Dotplot_All_Final.png"), width = 8, height =4)
```

##### Top10 DEGs

Top10 differentially expressed genes between all clusters

```
DefaultAssay(data) <- "RNA"
markers <- FindAllMarkers(data, only.pos = TRUE, min.pct = 0.25, logfc.threshold = 0.25, verbose =F)
top20 <- markers %>% group_by(cluster) %>% top_n(n = 20, wt = avg_log2FC)
top10 <- markers %>% group_by(cluster) %>% top_n(n = 10, wt = avg_log2FC)

kableExtra::kable(top10 %>%
                    select(cluster, gene) %>%
                    pivot_wider(names_from = "cluster", values_from = "gene"))
```

| Excitatory | Inhibitory | Astrocytes | Microglia | Oligodendrocytes | OPC | Pericytes | Endothelial | VLMCs |
| --- | --- | --- | --- | --- | --- | --- | --- | --- |
| AC011287…. | GRIK1, B…. | LINC0029…. | LHFPL3, …. | ST18, AC…. | FLT1, AT…. | SLC38A11…. | APBB1IP,…. | CEMIP, T…. |

##### Heatmap of DEGs

Heatmap of differentially expressed genes for each identity class

```
Idents(data) <- "CellType"
data@meta.data[["CellType"]] <- factor(data@meta.data[["CellType"]], 
                            levels=c(
                              "Excitatory",
                              "Inhibitory",
                              "Astrocytes",
                              "Oligodendrocytes",
                              "OPC",
                              "Microglia",
                              "VLMCs",  
                              "Endothelial", 
                              "Pericytes"))
data <- ScaleData(data, features = top10$gene)

DoHeatmap(data, features = top10$gene,size = 2) + 
 theme(axis.text.y = element_text(size = 4),
       legend.text = element_text(size = 8),
       panel.spacing.x = unit(0.2, "cm"))
```

```
ggsave(paste0(mainpath,"/", Sys.Date(), "/Plots/Heatmap_All_Clusters_Top10DEGs.png"), height = 5, width=10)
```

#### Cell type proportions

```
## Add cells per cluster annotation 
cells_per_cluster <- as.data.frame.matrix(table(data@meta.data[["RNA_snn_res.0.2"]],
                                                data@meta.data[["Brainregion"]])) %>%
  rownames_to_column("Cluster") %>%
  mutate(CellType = case_when(
    Cluster == '0'~ "unspecific",
  Cluster == '1'~ "Oligodendrocytes",
  Cluster == '2' ~ "Oligodendrocytes",
  Cluster == '3' ~ "Excitatory",
  Cluster == '4' ~ "Astrocytes",
  Cluster == '5' ~ "Endothelial",
  Cluster == '6' ~ "Microglia",
  Cluster == '7' ~ "OPC",
  Cluster == '8' ~ "Inhibitory",
  Cluster == '9' ~ "Excitatory",
  Cluster == '10' ~ "Pericytes",
  Cluster == '11' ~ "VLMCs")) %>%
  filter(!Cluster %in% c('0')) %>%
  pivot_longer(cols = -c(CellType, Cluster)) %>%
  unite(Group, CellType, name) %>%
  arrange(value)
```

```
cells_per_cluster  %>%
  mutate(Cluster = as.numeric(Cluster)) %>%
  arrange(Cluster) %>%
  separate(Group, into = c("CellType", "Brainregion")) %>%
  mutate(Brainregion = case_when(
    Brainregion == "Cortex" ~ "Cort",
    Brainregion == "Hippocampus" ~ "Hipp",
    Brainregion == "Putamen" ~ "Put",
    Brainregion == "WhiteMatter" ~ "WM",
  )) %>%
  na.omit() %>%
  group_by(CellType, Brainregion) %>%
  mutate(count =sum(value)) %>%
  dplyr::select(-c(Cluster, value)) %>%
  unique() %>%
  ggplot(aes(x =Brainregion,
             y= count, fill = fct_relevel(CellType, "Pericytes", "Endothelial", "VLMCs", "Microglia", 
                                          "OPC", "Oligodendrocytes", "Astrocytes", "Inhibitory", "Excitatory"))) +
  geom_bar(position = "fill", stat="identity",
           size = 0.2, color = "black") +
  scale_fill_manual(values = unlist(col_cells)) +
  labs(y="Fraction of total") +
  theme_minimal() +
  theme(axis.text.x = element_text(angle = 45, hjust = 1, size = 12),
        axis.text.y = element_text(size = 12),
        axis.title.y = element_text(size = 12),
        legend.text = element_text(size = 12),
        legend.title = element_blank(),
        axis.title.x = element_blank())
```

```
ggsave(paste0(mainpath,"/", Sys.Date(), "/Plots/Figure3C_Proportions.png"), width = 4, height =3)#width = 7.5, height = 6.3)
```

---

### Data downsampling

The amount of nuclei is unbalanced between the datasets and was downsampled to 5000 nuclei per dataset/voxel (dashed line in plot) for any downstream analyses:

```
cells_per_cluster %>%
  separate(Group, into = c("CellType", "Brainregion")) %>%
  group_by(Brainregion) %>%
  mutate(sum = sum(value)) %>%
  select(Brainregion, sum) %>%
  unique() %>%
  ggplot(aes(x = Brainregion, y = sum, fill = Brainregion)) +
  geom_bar(stat = "identity") +
  scale_fill_manual(values = unlist(col_vox)) +
  labs(y = "Sum of nuclei") +
  labs(x = "Voxel/Brainregion") +
  labs(title = "Sum of nuclei per voxel") +
  geom_hline(yintercept = 5000, linetype = "dashed") + 
  theme_classic() +
  theme(
    legend.position ="none",
    axis.text = element_text(size = 10),
    axis.title = element_text(size = 12)
  )
```

```
set.seed(666)
data <- subset(x = data, downsample = 5000)

DimPlot(data,label=F, label.size = 5,reduction = "umap", 
        group.by ="Brainregion", cols = unlist(col_vox))  +
  labs(title = "Downsampled data") +
  guides(color = guide_legend(override.aes = list(size = 6))) +
  theme(axis.title = element_text(size = 20),
        axis.text = element_text(size = 18),
        legend.text = element_text(size = 20))
```

```
ggsave(paste0(mainpath,"/", Sys.Date(), "/Plots/Final_UMAP_downsampled.png"), width = 5, height =4.5)
```

We further removed poorly represented cell groups with <20 cells per voxel. The following groups were removed:

```
remove <- cells_per_cluster %>%
  filter((value <20 | value == 20) & value > 0)  %>%
  rename(total_number_cells = value)
remove
```

```
## # A tibble: 2 × 3
##   Cluster Group                  total_number_cells
##   <chr>   <chr>                               <int>
## 1 8       Inhibitory_WhiteMatter                  1
## 2 11      VLMCs_WhiteMatter                      11
```

```
data <- subset(x = data, subset = ((Brainregion == "WhiteMatter" & seurat_clusters == 8) |
                                     Brainregion == "WhiteMatter" & seurat_clusters == 11),
               invert = TRUE)
```

---

### Mitochondrial analysis

For the mitochondrial analysis, we used the list of 1136 mitochondrial genes from the human MitoCarta3.0 dataset^11^, as well as genes lists and pathway annotations for

- only OxPhos pathways
- all mitochondrial pathways

```
## Mitocarta
mitocarta_sheet4 <- readxl::read_xls(here::here(mainpath, "HumanMitoCarta3_0.xls"), sheet = 4) %>%
  dplyr::select(MitoPathway, Genes) %>%
  na.omit() 
gene_to_pathway <- splitstackshape::cSplit(mitocarta_sheet4, 'Genes', ',') %>%
  column_to_rownames("MitoPathway") %>%
  t() %>%
  as.data.frame() 
gene_to_pathway <- gene_to_pathway %>%
  pivot_longer(cols = colnames(gene_to_pathway), names_to = "Pathway", values_to = "Gene") %>%
  na.omit() 
mitogenes <- readxl::read_xls(here::here(mainpath, "HumanMitoCarta3_0.xls"), sheet = 2) %>%
  select(Symbol)
mitogenes <- mitogenes$Symbol


### OxPhos genes and pathways
oxphos_genes <- gene_to_pathway %>%
  filter(Pathway %in% "OXPHOS")
oxphos_genes <- oxphos_genes$Gene

oxphos_pathways <- c("CI assembly factors",
                        "CI subunits",
                        "CII assembly factors",
                        "CII subunits",
                        "CIII assembly factors",
                        "CIII subunits",
                        "CIV assembly factors",
                        "CIV subunits",
                        "Complex I",
                        "Complex II",
                        "Complex III",
                        "Complex IV",
                        "Complex V",
                        "CV assembly factors",
                        "CV subunits",
                        "Cytochrome C",
                        "OXPHOS",
                        "OXPHOS assembly factors",
                        "OXPHOS subunits" )
```

A total of 1118 mitochondrial genes was found in the dataset. The missing 18 genes (i.e either not found or not expressed) are listed below:

```
mito_data <- NormalizeData(data,normalization.method = "LogNormalize", scale.factor = 10000)
mito_data <- subset(mito_data, features = mitogenes)
pseudobulk_mito_voxel <- AverageExpression(mito_data, group.by = c("orig.ident"), slot = "data", return.seurat=F)
## Since slot="data", the data is assumed to be log transformed. Return.seurat = F, average non log values are returned
pseudobulk_mito_voxel  <-pseudobulk_mito_voxel[["RNA"]] %>%
  as.data.frame() %>%
  rownames_to_column("Gene") %>%
  pivot_longer(cols = -Gene) %>%
  group_by(Gene) %>%
  mutate(sum = sum(value)) %>%
  filter(!sum %in% 0) %>%
  select(-sum) %>%
  pivot_wider(names_from = "Gene", values_from = "value") 
genes_found <- sort(unique(colnames(pseudobulk_mito_voxel[-1])))
not_found <- sort(mitogenes[which(!mitogenes %in% genes_found)])
not_found
```

```
##  [1] "ACOD1"         "AGXT"          "ATP5MF-PTCD1"  "CYP11B1"       "CYP11B2"       "FTMT"          "GARS1"         "GPX1"          "KARS1"         "MARCHF5"       "MCCD1"         "MPC1L"        
## [13] "MTARC1"        "MTARC2"        "MYG1"          "PDHA2"         "RP11_469A15.2" "SLC25A47"      "SLC25A52"      "SPATA19"
```

##### Pseudobulk expression per brain region

For each **brain region** we calculated the pseudobulk expression of each mitochondrial gene using all cells from each respective voxel (downsampled data). We removed genes that were not expressed in any cell.

```
mito_data <- NormalizeData(data,normalization.method = "LogNormalize", scale.factor = 10000)
mito_data <- subset(mito_data, features = mitogenes)
pseudobulk_mito <- AverageExpression(mito_data, group.by = c("orig.ident"), slot = "data", return.seurat=F)
## Since slot="data", the data is assumed to be log transformed. Return.seurat = F, average non log values are returned 
pseudobulk_mito  <- pseudobulk_mito[["RNA"]] %>%
  as.data.frame() %>%
  rownames_to_column("Gene") %>%
  pivot_longer(cols = -Gene) %>%
  mutate(name = case_when(
    name == "voxel1" ~ "Cort",
    name == "voxel2" ~ "Hipp",
    name == "voxel3" ~ "Put",
    name == "voxel4" ~ "WM"
  )) %>%
  rename(Voxel = name) %>%
  group_by(Gene) %>%
  mutate(sum = sum(value)) %>%
  filter(!sum %in% 0) %>%
  select(-sum) %>%
  pivot_wider(names_from = "Gene", values_from = "value")
```

We next calculated pathway scores for each mitochondrial pathway using its respective gene list as annotated in MitoCarta3.0^11^, and kept all OxPhos pathways. The data was z-score transformed between voxels (relative pathway scores between the brain regions)

```
pseudobulk_mitopathway_scores <- pseudobulk_mito %>%
  pivot_longer(cols = -c(Voxel), names_to = "Gene") %>%
  full_join(gene_to_pathway, by = "Gene") %>%
  filter(!is.na(Pathway)) %>%
  group_by(Voxel,Pathway) %>%
  mutate(score = mean(value, na.rm= T)) %>%
  select(-c(Gene, value)) %>%
  unique() %>%
  na.omit() 

pseudobulk_oxphos_scores <- pseudobulk_mitopathway_scores %>%
  filter(Pathway %in% oxphos_pathways) %>%
  mutate(Pathway =case_when(
    Pathway == "Complex I" ~ "CI exprs",
    Pathway == "Complex II" ~ "CII exprs",
    Pathway == "Complex III" ~ "CIII exprs",
    Pathway == "Complex IV" ~ "CIV exprs",
    Pathway == "Complex V" ~ "CV exprs",
    Pathway == "Cytochrome C"~"Cyt C",
    Pathway == "OXPHOS"~"OxPhos",
    Pathway == "OXPHOS assembly factors"~"OxPhos assembly factors",
    Pathway == "OXPHOS subunits"~"OxPhos subunits",
    TRUE ~ Pathway))  %>%
## Zscore
  group_by(Pathway) %>%
  mutate(mean = mean(score, na.rm = T)) %>%
  mutate(sd = sd(score, na.rm = T)) %>%
  mutate(score = (score - mean) / sd) %>%
  select(-c(mean, sd)) %>%
  pivot_wider(names_from = "Pathway", values_from = "score")
```

We further calculated the average mitochondrial expression across all mitochondrial genes (estimated transcription-based mitocontent).

```
pseudobulk_mito_content <- pseudobulk_mito %>%
  pivot_longer(cols = -Voxel) %>%
  group_by(Voxel) %>%
  mutate(estim_content = mean(value, na.rm = T)) %>%
  select(-c(name, value)) %>%
  unique()
```

Finally, we combined the gene expression pseudobulk mitopathway data with MRC activity measurements, calculated the relative expression between the 4 voxels (for comparability), and visualized all measures in a heatmap (Figure 3d).

```
additional_data <- read.csv(here::here(mainpath, "voxel_MHI_data.csv")) %>%
  select(Voxel, MRC, TRC, CI, CII, CIV, MitoD) %>%
   mutate(Voxel = case_when(
    Voxel == "voxel1" ~ "Cort",
    Voxel == "voxel2" ~ "Hipp",
    Voxel == "voxel3" ~ "Put",
    Voxel == "voxel4" ~ "WM"
  )) %>%
  full_join(pseudobulk_mito_content) %>%
  pivot_longer(cols = -Voxel) %>%
  mutate(name = case_when(
    name == "CI" ~ "CI-activity",
    name == "CII" ~ "CII-activity",
    name == "CIV" ~ "CIV-activity",
    TRUE ~ name
  )) %>%
  group_by(name) %>%
  mutate(mean = mean(value, na.rm = T)) %>%
  mutate(sd = sd(value, na.rm = T)) %>%
  mutate(value = (value - mean) / sd) %>%
  select(-c(mean, sd)) %>%
  pivot_wider(names_from = "name", values_from = "value") 

pseudobulk_oxphos_scores <- pseudobulk_oxphos_scores %>%
  full_join(additional_data) %>%
  select(Voxel, 
  OxPhos, `OxPhos assembly factors`, `OxPhos subunits`,
  `CI exprs`, `CI subunits`,`CI assembly factors`,`CI-activity`,
  `CII exprs`,`CII subunits`, `CII assembly factors`,`CII-activity`,
  `CIII exprs`,`CIII subunits`, `CIII assembly factors`,
  `CIV exprs`,`CIV subunits`, `CIV assembly factors`,`CIV-activity`,
  `CV exprs`,`CV subunits`, `CV assembly factors`, 
  `Cyt C`, estim_content, 
  MitoD, MRC, TRC  )


## Transpose data
exprs <- t(pseudobulk_oxphos_scores%>%column_to_rownames("Voxel"))

## Heatmap annotation
anno_df <- as.data.frame(pseudobulk_oxphos_scores) %>%
  pivot_longer(cols = -c(Voxel), names_to= "Gene") %>%
  na.omit() %>%
  pivot_wider(names_from = "Gene", values_from = "value") %>%
  dplyr::select(Voxel)
col_anno = columnAnnotation(
  Voxel = anno_df$Voxel, 
  col=list(
  Voxel = c(
    "Cort" = "#4F5088", 
    "Hipp" = "#A5458D", 
    "Put" = "#5D9E4F", 
    "WM" = "#4E77BC"
  )
  ),
  show_annotation_name = F,
  show_legend =  T,
  annotation_legend_param = list(nrow=12)) 

## Build the heatmap
hm <- Heatmap(exprs, 
        name = "Rel. expression",
        show_column_dend=F,
        show_column_names =T,
        show_row_names =T,
        row_title="",
        cluster_columns =F,
        cluster_rows =F,
        row_names_gp = grid::gpar(fontsize = 8),
        row_title_gp = grid::gpar(fontsize = 10),
        column_names_rot = 45,
        column_names_gp = grid::gpar(fontsize = 12),
        column_title_gp = grid::gpar(fontsize = 10),
        top_annotation=col_anno,
        width = unit(5, "cm")
           )
pdf(paste0(mainpath,"/", Sys.Date(), "/Plots/Figure3D_Heatmap_oxphos_pseudobulk_per_voxel.pdf"), width = 8, height = 4)
draw(hm)
invisible(dev.off())
draw(hm)
```

```
rm(pseudobulk_mito, pseudobulk_mitopathway_scores, pseudobulk_oxphos_scores, pseudobulk_mito_content, additional_data, exprs, anno_df, col_anno, hm)
```

##### Raw expression (scaled across the dataset)

For each **brain region and celltype** we calculated the pseudobulk expression using all cells from each respective cell type in each voxel (downsampled data).

```
mito_data <- NormalizeData(data,normalization.method = "LogNormalize", scale.factor = 10000)
mito_data <- subset(mito_data, features = mitogenes)
pseudobulk_mito_vox_cell <- AverageExpression(mito_data, group.by = c("orig.ident", "CellType"),
                                              slot = "data", return.seurat=F)

pseudobulk_mito_vox_cell  <-pseudobulk_mito_vox_cell[["RNA"]] %>%
  as.data.frame() %>%
  rownames_to_column("Gene") %>%
  pivot_longer(cols = -Gene) %>%
  separate(name, into = c("Voxel", "CellType"), remove = T) %>%
  mutate(Voxel = case_when(
    Voxel == "voxel1" ~ "Cort",
    Voxel == "voxel2" ~ "Hipp",
    Voxel == "voxel3" ~ "Put",
    Voxel == "voxel4" ~ "WM"
  )) %>%
  group_by(Gene) %>%
  mutate(sum = sum(value)) %>%
  filter(!sum %in% 0) %>%
  select(-sum)
```

**OxPhos complex bivariate plots**

```
pseudobulk_mito_vox_cell_pathways = pseudobulk_mito_vox_cell %>%
  full_join(gene_to_pathway, by = "Gene") %>%
  filter(!is.na(Pathway)) %>%
  group_by(Voxel,CellType, Pathway) %>%
  mutate(score = mean(value, na.rm= T)) %>%
  na.omit() %>%
  select(-c(Gene, value)) %>%
  unique() 

mito_raw <- pseudobulk_mito_vox_cell_pathways %>%
  pivot_wider(names_from = "Pathway", values_from = "score") %>%
  unite(name, Voxel, CellType, remove =F) %>%
  column_to_rownames("name")

col_vox <- list(  
    "Cort" = "#4F5088", 
    "Hipp" = "#A5458D", 
    "Put" = "#5D9E4F", 
    "WM" = "#4E77BC")

theme <-   theme(
    legend.title = element_blank(),
    axis.title = element_text(size = 10), 
    axis.text = element_text(size =8)) 

p <- mito_raw %>%
  ggplot(aes(x = `Complex II`, y = `Complex I`)) +
  geom_point(aes(fill = Voxel, color = Voxel), alpha = 0.6, size = 4) +
  scale_color_manual(values = col_vox) +
  scale_fill_manual(values = col_vox) +
  labs(title = "CI + CII") + 
  theme_bw() +
  theme(legend.position = "none")
p
```

```
ggsave(paste0(mainpath,"/", Sys.Date(), "/Plots/Figure3E_Bivariate_CI_CIV_Raw.png" ), height = 2.8, width = 2.8)
```

```
p <- mito_raw %>%
  ggplot(aes(x = `Complex III`, y = `Complex I`)) +
  geom_point(aes(fill = Voxel, color = Voxel), alpha = 0.6, size = 4) +
  scale_color_manual(values = col_vox) +
  scale_fill_manual(values = col_vox) +
  labs(title = "CI + CIII") + 
  theme_bw() +
  theme(legend.position = "none")
p
```

```
p <- mito_raw %>%
  ggplot(aes(x = `Complex IV`, y = `Complex I`)) +
  geom_point(aes(fill = Voxel, color = Voxel), alpha = 0.6, size = 4) +
  scale_color_manual(values = col_vox) +
  scale_fill_manual(values = col_vox) +
  labs(title = "CI + CIV (Figure 3e)") + 
  theme_bw() +
  theme(legend.position = "none")
p
```

```
p <- mito_raw %>%
  ggplot(aes(x = `Complex V`, y = `Complex I`)) +
  geom_point(aes(fill = Voxel, color = Voxel), alpha = 0.6, size = 4) +
  scale_color_manual(values = col_vox) +
  scale_fill_manual(values = col_vox) +
  labs(title = "CI + CV") + 
  theme_bw() +
  theme(legend.position = "none")
p
```

**MitoPathway Heatmap (Figure 3f)**

Next, we calculated pathway scores for each mitochondrial pathway (as annotated in MitoCarta3.0^11^) using its respective gene list, and scaled the data across the entire dataset.

```
pseudobulk_mito_vox_cell_pathways = pseudobulk_mito_vox_cell %>%
  full_join(gene_to_pathway, by = "Gene") %>%
  filter(!is.na(Pathway)) %>%
  group_by(Voxel,CellType, Pathway) %>%
  mutate(score = mean(value, na.rm= T)) %>%
  na.omit() %>%
  select(-c(Gene, value)) %>%
  unique() 

mito_raw <- pseudobulk_mito_vox_cell_pathways %>%
  pivot_wider(names_from = "Pathway", values_from = "score") %>%
  unite(name, Voxel, CellType, remove =F) %>%
  column_to_rownames("name")

## Transpose and scale matrix
exprs <- t(scale(mito_raw[,-c(1:2)]))

## Heatmap annotation
anno_df <- as.data.frame(mito_raw) %>%
  pivot_longer(cols = -c(Voxel, CellType), names_to= "Pathway") %>%
  na.omit() %>%
  pivot_wider(names_from = "Pathway", values_from = "value") %>%
  dplyr::select(Voxel, CellType)
col_anno = columnAnnotation(
  Celltype=anno_df$CellType,
  Voxel = anno_df$Voxel, 
  col=list(
  Celltype  =unlist(col_cells),
  Voxel = c(
    "Cort" = "#4F5088", 
    "Hipp" = "#A5458D", 
    "Put" = "#5D9E4F", 
    "WM" = "#4E77BC")
  ),
  show_annotation_name = F,
  show_legend =  T,
  annotation_legend_param = list(nrow=12)) 
col_dist    = dist(t(as.matrix(exprs)), method="euclidean" )
coldend     = hclust(col_dist, method="ward.D2")  
row_dist    = dist(as.matrix(exprs), method="euclidean" )
rowdend     = hclust(row_dist, method="ward.D2")

hm <- Heatmap(exprs, 
        name = "Rel. expression",
        show_column_dend=T,
        show_column_names =T,
        show_row_names =F,
        show_row_dend=F,
        row_title="",
        cluster_columns =coldend,
        cluster_rows =rowdend,
        column_names_rot = 45,
        row_names_gp = grid::gpar(fontsize = 8),
        row_title_gp = grid::gpar(fontsize = 10),
        column_names_gp = grid::gpar(fontsize = 8),
        column_title_gp = grid::gpar(fontsize = 10),
        top_annotation=col_anno,
        width = unit(12, "cm"))
pdf(paste0(mainpath,"/", Sys.Date(), "/Plots/Figure3F_Heatmap_pathways_scaled.pdf"), width = 25, height = 5)
draw(hm)
invisible(dev.off())
Heatmap(exprs,
        name = "Rel. expression",
        show_column_dend=T,
        show_column_names =T,
        show_row_names =T,
        show_row_dend=T,
        row_title="Mitopathways",
        cluster_columns =coldend,
        cluster_rows =rowdend,
        column_names_rot = 45,
        row_names_gp = grid::gpar(fontsize = 2),
        row_title_gp = grid::gpar(fontsize = 10),
        column_names_gp = grid::gpar(fontsize = 6),
        column_title_gp = grid::gpar(fontsize = 10),
        top_annotation=col_anno,
        width = unit(8, "cm")
           )
```

```
rm(pseudobulk_mito_vox_cell_zscore, exprs, anno_df, col_anno, hm)
```

##### Normalized data (by voxel)

The data was z-score transformed within each voxel.

**OxPhos complex bivariate plots**

```
mito_zscore <- pseudobulk_mito_vox_cell_pathways %>%
  group_by(Voxel, Pathway) %>%
  mutate(mean = mean(score, na.rm = T)) %>%
  mutate(sd = sd(score, na.rm =T)) %>%
  mutate(score = (score-mean)/sd) %>%
  select(-c(mean, sd)) %>%
  pivot_wider(names_from = "Pathway", values_from = "score") %>%
  unite(name, Voxel, CellType, remove =F) %>%
  column_to_rownames("name")

p <- mito_zscore %>%
  ggplot(aes(x = `Complex II`, y = `Complex I`)) +
  geom_point(aes(fill = CellType, color = CellType), alpha = 0.6, size = 4) +
  scale_color_manual(values = col_cells) +
  scale_fill_manual(values = col_cells) +
  labs(title = "CI + CII") + 
  theme_bw() +
  theme(legend.position = "none")
p
```

```
 ggsave(paste0(mainpath,"/", Sys.Date(), "/Plots/Figure3G_Bivariate_CI_CIV_Zscore.png" ), height = 2.8, width = 2.8)
```

```
p <- mito_zscore %>%
  ggplot(aes(x = `Complex III`, y = `Complex I`)) +
  geom_point(aes(fill = CellType, color = CellType), alpha = 0.6, size = 4) +
  scale_color_manual(values = col_cells) +
  scale_fill_manual(values = col_cells) +
  labs(title = "CI + CIII") + 
  theme_bw() +
  theme(legend.position = "none")
p
```

```
p <- mito_zscore %>%
  ggplot(aes(x = `Complex IV`, y = `Complex I`)) +
  geom_point(aes(fill = CellType, color = CellType), alpha = 0.6, size = 4) +
  scale_color_manual(values = col_cells) +
  scale_fill_manual(values = col_cells) +
  labs(title = "CI + CIV (Figure 3g)") + 
  theme_bw() +
  theme(legend.position = "none")
p
```

```
p <- mito_zscore %>%
  ggplot(aes(x = `Complex V`, y = `Complex I`)) +
  geom_point(aes(fill = CellType, color = CellType), alpha = 0.6, size = 4) +
  scale_color_manual(values = col_cells) +
  scale_fill_manual(values = col_cells) +
  labs(title = "CI + CV") + 
  theme_bw() +
  theme(legend.position = "none")
p
```

**MitoPathway Heatmap (Figure 3h)**

```
pseudobulk_mito_vox_cell_zscore <- pseudobulk_mito_vox_cell %>%
  full_join(gene_to_pathway, by = "Gene") %>%
  filter(!is.na(Pathway)) %>%
  group_by(Voxel,CellType, Pathway) %>%
  mutate(score = mean(value, na.rm= T)) %>%
  select(-c(Gene, value)) %>%
  unique() %>%
  na.omit() %>%
  ## zscore
  group_by(Pathway, Voxel) %>%
  mutate(mean = mean(score, na.rm = T)) %>%
  mutate(sd = sd(score, na.rm =T)) %>%
  mutate(score = (score-mean)/sd) %>%
  select(-c(mean, sd)) %>%
  pivot_wider(names_from = "Pathway", values_from = "score") %>%
  unite(name, Voxel, CellType, remove =F) %>%
  column_to_rownames("name")

## Transpose matrix
exprs <- t(pseudobulk_mito_vox_cell_zscore[,-c(1:2)])

## Heatmap annotation
anno_df <- as.data.frame(pseudobulk_mito_vox_cell_zscore) %>%
  pivot_longer(cols = -c(Voxel, CellType), names_to= "Pathway") %>%
  na.omit() %>%
  pivot_wider(names_from = "Pathway", values_from = "value") %>%
  dplyr::select(Voxel, CellType)
col_anno = columnAnnotation(
  Celltype=anno_df$CellType,
  Voxel = anno_df$Voxel, 
  col=list(
  Celltype  =unlist(col_cells),
  Voxel = c(
    "Cort" = "#4F5088", 
    "Hipp" = "#A5458D", 
    "Put" = "#5D9E4F", 
    "WM" = "#4E77BC")
  ),
  show_annotation_name = F,
  show_legend =  T,
  annotation_legend_param = list(nrow=12)) 
col_dist    = dist(t(as.matrix(exprs)), method="euclidean" )
coldend     = hclust(col_dist, method="ward.D2")  
row_dist    = dist(as.matrix(exprs), method="euclidean" )
rowdend     = hclust(row_dist, method="ward.D2")

hm <- Heatmap(exprs, 
        name = "Rel. expression",
        show_column_dend=T,
        show_column_names =T,
        show_row_names =F,
        show_row_dend=F,
        row_title="",
        cluster_columns =coldend,
        cluster_rows =rowdend,
        column_names_rot = 45,
        row_names_gp = grid::gpar(fontsize = 8),
        row_title_gp = grid::gpar(fontsize = 10),
        column_names_gp = grid::gpar(fontsize = 8),
        column_title_gp = grid::gpar(fontsize = 10),
        top_annotation=col_anno,
        width = unit(12, "cm"))
pdf(paste0(mainpath,"/", Sys.Date(), "/Plots/Figure3H_Heatmap_pathways_zscore.pdf"), width = 25, height = 5)
draw(hm)
invisible(dev.off())
Heatmap(exprs,
        name = "Rel. expression",
        show_column_dend=T,
        show_column_names =T,
        show_row_names =T,
        show_row_dend=T,
        row_title="Mitopathways",
        cluster_columns =coldend,
        cluster_rows =rowdend,
        column_names_rot = 45,
        row_names_gp = grid::gpar(fontsize = 2),
        row_title_gp = grid::gpar(fontsize = 10),
        column_names_gp = grid::gpar(fontsize = 6),
        column_title_gp = grid::gpar(fontsize = 10),
        top_annotation=col_anno,
        width = unit(8, "cm")
           )
```

```
rm(pseudobulk_mito_vox_cell_zscore, exprs, anno_df, col_anno, hm)
```

### R session

```
sessionInfo()
```

```
## R version 4.2.2 (2022-10-31)
## Platform: aarch64-apple-darwin20 (64-bit)
## Running under: macOS 14.1.1
## 
## Matrix products: default
## BLAS:   /Library/Frameworks/R.framework/Versions/4.2-arm64/Resources/lib/libRblas.0.dylib
## LAPACK: /Library/Frameworks/R.framework/Versions/4.2-arm64/Resources/lib/libRlapack.dylib
## 
## locale:
## [1] en_US.UTF-8/en_US.UTF-8/en_US.UTF-8/C/en_US.UTF-8/en_US.UTF-8
## 
## attached base packages:
## [1] parallel  grid      stats     graphics  grDevices utils     datasets  methods   base     
## 
## other attached packages:
##  [1] ROCR_1.0-11           KernSmooth_2.23-22    fields_15.2           viridisLite_0.4.2     spam_2.10-0           fontawesome_0.5.2     Azimuth_0.5.0         shinyBS_0.61.1       
##  [9] DoubletFinder_2.0.3   RColorBrewer_1.1-3    ComplexHeatmap_2.12.1 Seurat_5.0.1          SeuratObject_5.0.1    sp_2.1-3              lubridate_1.9.3       forcats_1.0.0        
## [17] stringr_1.5.1         dplyr_1.1.4           purrr_1.0.2           readr_2.1.5           tidyr_1.3.1           tibble_3.2.1          ggplot2_3.4.4         tidyverse_2.0.0      
## 
## loaded via a namespace (and not attached):
##   [1] ica_1.0-3                         svglite_2.1.3                     RcppRoll_0.3.0                    Rsamtools_2.14.0                  foreach_1.5.2                    
##   [6] lmtest_0.9-40                     rprojroot_2.0.4                   crayon_1.5.2                      MASS_7.3-60.0.1                   rhdf5filters_1.10.1              
##  [11] nlme_3.1-164                      rmdformats_1.0.4                  rlang_1.1.3                       readxl_1.4.3                      XVector_0.38.0                   
##  [16] irlba_2.3.5.1                     limma_3.52.4                      filelock_1.0.3                    BiocParallel_1.32.6               rjson_0.2.21                     
##  [21] CNEr_1.34.0                       bit64_4.0.5                       glue_1.7.0                        harmony_1.2.0                     poweRlaw_0.80.0                  
##  [26] sctransform_0.4.1                 vipor_0.4.7                       spatstat.sparse_3.0-3             AnnotationDbi_1.60.2              SeuratDisk_0.0.0.9021            
##  [31] BiocGenerics_0.44.0               shinydashboard_0.7.2              dotCall64_1.1-1                   spatstat.geom_3.2-8               tidyselect_1.2.0                 
##  [36] SummarizedExperiment_1.28.0       fitdistrplus_1.1-11               XML_3.99-0.16.1                   zoo_1.8-12                        GenomicAlignments_1.34.1         
##  [41] xtable_1.8-4                      RcppHNSW_0.5.0                    magrittr_2.0.3                    evaluate_0.23                     cli_3.6.2                        
##  [46] zlibbioc_1.44.0                   rstudioapi_0.15.0                 miniUI_0.1.1.1                    bslib_0.6.1                       fastmatch_1.1-4                  
##  [51] ensembldb_2.22.0                  fastDummies_1.7.3                 maps_3.4.2                        shiny_1.8.0                       xfun_0.41                        
##  [56] clue_0.3-65                       cluster_2.1.6                     caTools_1.18.2                    KEGGREST_1.38.0                   ggrepel_0.9.5                    
##  [61] listenv_0.9.1                     TFMPvalue_0.0.9                   Biostrings_2.66.0                 png_0.1-8                         future_1.33.1                    
##  [66] withr_3.0.0                       bitops_1.0-7                      plyr_1.8.9                        cellranger_1.1.0                  AnnotationFilter_1.22.0          
##  [71] pracma_2.4.4                      SeuratData_0.2.2.9001             pillar_1.9.0                      GlobalOptions_0.1.2               cachem_1.0.8                     
##  [76] GenomicFeatures_1.50.4            fs_1.6.3                          hdf5r_1.3.9                       GetoptLong_1.0.5                  vctrs_0.6.5                      
##  [81] ellipsis_0.3.2                    generics_0.1.3                    tools_4.2.2                       beeswarm_0.4.0                    munsell_0.5.0                    
##  [86] DelayedArray_0.24.0               fastmap_1.1.1                     compiler_4.2.2                    abind_1.4-5                       httpuv_1.6.14                    
##  [91] rtracklayer_1.58.0                plotly_4.10.4                     GenomeInfoDbData_1.2.9            gridExtra_2.3                     lattice_0.22-5                   
##  [96] deldir_2.0-2                      utf8_1.2.4                        later_1.3.2                       BiocFileCache_2.6.1               jsonlite_1.8.8                   
## [101] scales_1.3.0                      pbapply_1.7-2                     lazyeval_0.2.2                    promises_1.2.1                    doParallel_1.0.17                
## [106] splitstackshape_1.4.8             R.utils_2.12.3                    goftest_1.2-3                     spatstat.utils_3.0-4              reticulate_1.35.0                
## [111] rmarkdown_2.25                    cowplot_1.1.3                     textshaping_0.3.7                 Rtsne_0.17                        BSgenome_1.66.3                  
## [116] Biobase_2.58.0                    uwot_0.1.16                       igraph_2.0.1.1                    survival_3.5-7                    yaml_2.3.8                       
## [121] systemfonts_1.0.5                 htmltools_0.5.7                   memoise_2.0.1                     BiocIO_1.8.0                      IRanges_2.32.0                   
## [126] here_1.0.1                        digest_0.6.34                     RhpcBLASctl_0.23-42               mime_0.12                         rappdirs_0.3.3                   
## [131] httr2_1.0.0                       RSQLite_2.3.5                     future.apply_1.11.1               data.table_1.15.0                 blob_1.2.4                       
## [136] JASPAR2020_0.99.10                S4Vectors_0.36.2                  R.oo_1.26.0                       TFBSTools_1.36.0                  ragg_1.2.7                       
## [141] splines_4.2.2                     labeling_0.4.3                    Rhdf5lib_1.20.0                   googledrive_2.1.1                 ProtGenerics_1.30.0              
## [146] RCurl_1.98-1.14                   hms_1.1.3                         rhdf5_2.42.1                      colorspace_2.1-0                  ggbeeswarm_0.7.2                 
## [151] GenomicRanges_1.50.2              shape_1.4.6                       Signac_1.12.0                     ggrastr_1.0.2                     sass_0.4.8                       
## [156] Rcpp_1.0.12                       BSgenome.Hsapiens.UCSC.hg38_1.4.5 bookdown_0.37                     RANN_2.6.1                        circlize_0.4.15                  
## [161] fansi_1.0.6                       tzdb_0.4.0                        parallelly_1.36.0                 R6_2.5.1                          ggridges_0.5.6                   
## [166] lifecycle_1.0.4                   curl_5.2.0                        googlesheets4_1.1.1               leiden_0.4.3.1                    jquerylib_0.1.4                  
## [171] Matrix_1.6-5                      RcppAnnoy_0.0.22                  iterators_1.0.14                  spatstat.explore_3.2-6            htmlwidgets_1.6.4                
## [176] polyclip_1.10-6                   biomaRt_2.59.1                    timechange_0.3.0                  seqLogo_1.64.0                    mgcv_1.9-1                       
## [181] globals_0.16.2                    patchwork_1.2.0                   spatstat.random_3.2-2             progressr_0.14.0                  codetools_0.2-19                 
## [186] matrixStats_1.2.0                 GO.db_3.16.0                      gtools_3.9.5                      prettyunits_1.2.0                 dbplyr_2.4.0                     
## [191] RSpectra_0.16-1                   R.methodsS3_1.8.2                 GenomeInfoDb_1.34.9               gtable_0.3.4                      DBI_1.2.1                        
## [196] stats4_4.2.2                      tensor_1.5                        httr_1.4.7                        highr_0.10                        stringi_1.8.3                    
## [201] presto_1.0.0                      progress_1.2.3                    reshape2_1.4.4                    farver_2.1.1                      annotate_1.76.0                  
## [206] magick_2.8.2                      DT_0.31                           xml2_1.3.6                        EnsDb.Hsapiens.v86_2.99.0         shinyjs_2.1.0                    
## [211] kableExtra_1.4.0                  restfulr_0.0.15                   scattermore_1.2                   bit_4.0.5                         MatrixGenerics_1.10.0            
## [216] spatstat.data_3.0-4               pkgconfig_2.0.3                   gargle_1.5.2                      DirichletMultinomial_1.40.0       knitr_1.45
```
