## Supplementary Files1-4 and video1 for "A Human Brain Map of Mitochondrial Respiratory Capacity and Diversity": Supplementary File4 - Python code for NIfTI file conversion.py

```
#!/usr/bin/env python3
# -*- coding: utf-8 -*-
"""
Created on Mon Aug 29 14:44:56 2022

@author: victor

Convert all the .DNG files from a folder to the Nifti (.nii.gz) files,
saving them with the same name and in the same folder
# don' forget to pip install rawpy
"""

import rawpy  # To install with pip
import nibabel as nib
import numpy as np
import os
import sys

if len(sys.argv) != 2:
    print('Bad input.\n'
          'dng2nii.py is expecting 1 argument: the path to the folder containing the .DNG files.\n'
          'For example: python dng2nii.py /path/to/myPictures/')
else:
    folder = sys.argv[1]
    if not os.path.isdir(folder):
        raise FileNotFoundError('The input folder was not found (or is not a folder).')
    dngL = [f for f in os.listdir(folder) if os.path.splitext(f)[1].lower() == '.dng']
    for i, dngName in enumerate(dngL):
        print(f'Processing file {i + 1}/{len(dngL)} ({dngName})')
        niiPath = os.path.join(folder, os.path.splitext(dngName)[0] + '.nii.gz')
        if os.path.exists(niiPath):
            print(f'{niiPath} already exists. Skipping.')
            continue
        dngPath = os.path.join(folder, dngName)
        with rawpy.imread(dngPath) as raw:
            im = raw.postprocess()
        if len(im.shape) == 3:  # RGB image
            im = np.expand_dims(im, -2)  # Putting the RGB channel in the 4th dim with an empty 3rd dim
        im = np.moveaxis(im, [0, 1, 2], [2, 1, 0])  # Putting X and Y from DNG convention to Nifti convention
        im = np.flip(im, 2)
        affine = np.eye(4, dtype=int)
        affine[:3, -1] = -1 * np.ceil(np.array(im.shape[:3]) / 2).astype(int)  # Centering the 0 of the image
        im_nii = nib.Nifti1Image(im, affine)
        nib.save(im_nii, niiPath)
```
